# Transcription-Coupled Chromatin Reinforcement Maintains the Mature Cardiomyocyte State

**DOI:** 10.64898/2026.09.04.749300

**Authors:** Han-Hsuan Tang, Hsin-Yi Tseng, Chia-Yeh Lin, Cheng-Fu Kao

## Abstract

Chromatin homeostasis is fundamental to maintaining transcriptional programs that stabilize cell states while permitting plasticity for timely state transitions. However, mechanisms that actively sustain mature cell states after their establishment remain poorly defined despite being essential for lifelong maintenance of specialized cellular functions. Here we demonstrate that the transcription-coupled chromatin regulator RNF20 is crucial for maintaining chromatin accessibility at identity-associated promoters that support mature cardiomyocyte transcriptional programs in adult murine cardiomyocytes. Depletion of RNF20 progressively erodes the transcriptional, structural, and functional integrity of adult cardiomyocytes and activates AP-1-associated stress-responsive enhancer elements without cell-cycle re-entry. Spatial transcriptomics further revealed a subendocardial localization of this stress-responsive state. Together, these findings support a model in which transcription-coupled chromatin reinforcement continuously stabilizes the mature cardiomyocyte attractor state while constraining transitions toward pathological cell states.

**Teaser:** Mature cardiomyocyte identity requires continuous chromatin reinforcement to resist transitions toward pathological states.

## INTRODUCTION

A defining feature of multicellular organisms is the ability of a single genome to support a diversity of cell states with distinct functions. This diversity is orchestrated by transcription factors (TFs) and epigenetic mechanisms that establish state-specific transcriptional programs (*1–5*). Maintenance of cell state requires sustained expression of identity-defining genes together with repression of alternative cellular programs, thereby preserving specialized functions (*6, 7*). In dynamical systems, such stable cellular configurations can be viewed as attractor states, in which gene regulatory networks constrain cells within a defined transcriptional and functional state despite perturbations (*8, 9*). Chromatin homeostasis may contribute to the stability of such attractor states by preserving identity-defining transcriptional programs while permitting controlled transitions in response to developmental or environmental cues (*10*). Such active preservation of cell state is particularly important in long-lived postmitotic cells, including mature cardiomyocytes and neurons, whose specialized functions cannot be readily restored through endogenous cell replacement (*11*).

Cardiomyocytes are highly specialized contractile cells responsible for pumping blood throughout the body. To achieve their optimal function, cardiomyocytes undergo extensive postnatal maturation, including sarcomere organization, a metabolic switch from glycolysis to oxidative phosphorylation, and electrical coupling enabled by gap junctions through widespread transcriptional and chromatin remodeling (*12, 13*). Acquisition of the mature cardiomyocyte state is accompanied by gradual loss of proliferative capacity, thereby limiting myocardial regeneration after injury in mammals (*14*). Despite being postmitotic, mature cardiomyocytes retain a degree of plasticity and can undergo dedifferentiation in response to stress, an adaptive process marked by partial loss of mature transcriptional programs and reactivation of fetal gene expression (*15, 16*). Such dedifferentiation, however, does not necessarily restore proliferative or regenerative competence (*17–20*).

Despite being exempt from the extensive chromatin repackaging accompanying cell division, postmitotic cardiomyocytes remain subject to continuous transcription-associated histone turnover, especially in cis-regulatory elements (*21*). Histone turnover can passively dilute histone modifications, suggesting the need for faithful re-establishment of the histone-mark repertoire to sustain appropriate transcriptional programs and thereby cell-state stability. Consistent with this requirement, disruption of histone modifiers in adult cardiomyocytes perturbs chromatin homeostasis, transcriptional programs, and cardiac function (*22, 23*). Beyond the regulation of cis-regulatory elements, transcription itself imposes continuous demands on chromatin homeostasis by requiring repeated nucleosome disruption and restoration during RNA polymerase II (Pol II) passage, processes facilitated by elongation factors and histone chaperones, and coupled to histone modifications across transcribed gene bodies (*24*). Despite their fundamental roles in transcription, however, whether these transcription-coupled mechanisms are required to preserve the mature cardiomyocyte state remains unclear.

RNF20 and RNF40 form the principal E3 ligase for monoubiquitination of histone H2B at lysine 120 (H2Bub), a transcription-coupled chromatin modification enriched across active gene bodies (*25, 26*). H2Bub promotes transcriptional elongation by regulating nucleosome dynamics, including FACT-dependent nucleosome reassembly, providing a mechanism for preserving chromatin architecture during ongoing transcription (*27–29*). These properties position RNF20 at the interface between ongoing transcription and chromatin maintenance. Its established roles in differentiation (*30, 31*), cellular reprogramming (*32*), and postnatal cardiomyocyte maturation (*33, 34*), further provide the biological rationale to test whether this transcription-coupled mechanism remains required once a terminally differentiated state has been established. To directly test this possibility, we induced cardiomyocyte-specific *Rnf20* deletion in adult mice.

By integrating single-nucleus transcriptomics, cardiomyocyte chromatin accessibility profiling, spatial transcriptomics, cytoarchitectural analyses, and cardiac physiology, we found that RNF20 loss progressively eroded the mature cardiomyocyte transcriptional state, elicited spatially patterned injury-like remodeling without productive proliferation, and impaired cardiac function. Unlike canonical repressive chromatin mechanisms such as G9a (*22*), RNF20 is coupled to transcriptionally active chromatin, suggesting a distinct mode of epigenetic state maintenance. These findings support a model in which a continuous transcription-coupled chromatin regulation sustain the mature cardiomyocyte attractor state and prevent transitions toward pathological cell states, thereby preserving cardiac homeostasis.

## RESULTS

### RNF20 stabilizes the mature cardiomyocyte transcriptional state

If RNF20/H2Bub provides continuous chromatin reinforcement of the established mature cardiomyocyte state, H2Bub should remain preferentially associated with genes defining that state. We therefore examined H2Bub distribution across mature-state genes. Gene-body H2Bub was selectively enriched at mature genes relative to immature genes (Fig. 1A, Fig. S1, and Data S1; Please refer to supplementary text for detail of the construction of gene sets.). Notably, this preferential enrichment was observed despite a progressive decline in *Rnf20* expression during postnatal maturation (Fig. 1B and Fig. S2A), indicating that maintenance of the mature transcriptional program cannot be explained simply by global RNF20 abundance. Instead, these observations suggest that RNF20-dependent H2Bub may contribute to maintenance of the mature cardiomyocyte state by selectively supporting transcription of mature-state genes.

**Figure 1.**
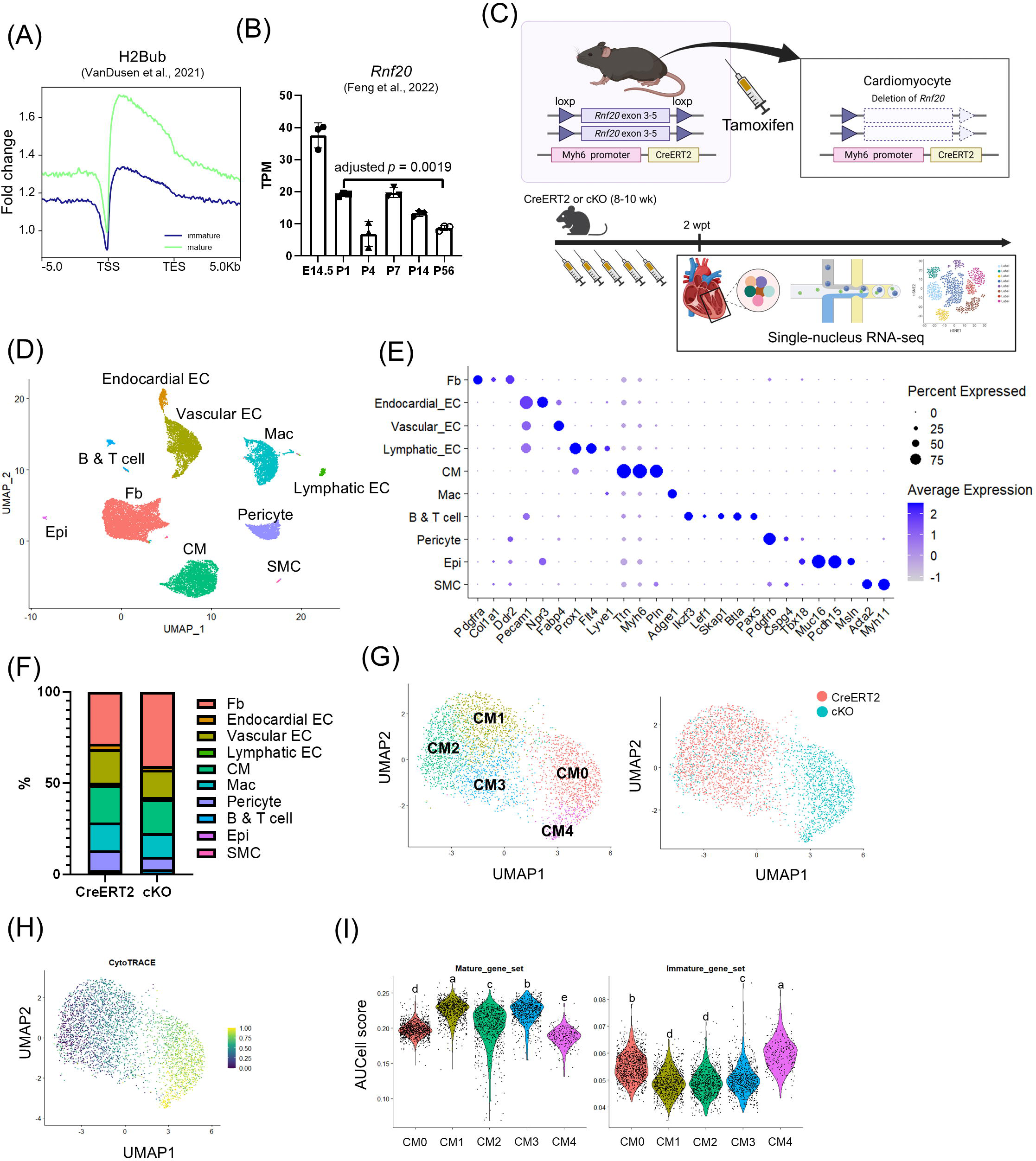
RNF20 stabilizes mature cardiomyocyte transcriptional state. (A) Metagene profile plot of H2Bub across maturation-associated genes in postnatal day 28 murine hearts (*34*), spanning gene bodies with 5 kb upstream and downstream flanking regions. The y-axis represents fold change of signal relative to input. TSS, transcription start site; TES, transcription end site. (B) Expression of *Rnf20* from isolated cardiomyocytes during cardiomyocyte maturation. The data were generated with n = 3 biological replicates and are presented as mean ± s.d.. TPM, transcripts per million. One-way ANOVA with Tukey’s test. (C) Experimental design for *Rnf20* conditional knockout mice (top) and snRNA-seq (bottom). (D) Uniform Manifold Approximation Projection (UMAP) visualization of clusters identified by Louvain algorithm with nuclei from both genotypes at 2 wpt. Fb, fibroblast; EC, endothelial cell; CM, cardiomyocyte; Mac, macrophage; Epi, epicardial cell; SMC, smooth muscle cell. (E) Dot plot of expression of known marker genes among cardiac cell types. (F) Proportion of cell types. (G) UMAP visualization of subclustered cardiomyocytes colored by cell states (left) or by genotype (right). (H) CytoTRACE scores in cardiomyocytes. (I) AUCell scores for mature (left) and immature (right) gene set in cardiomyocytes. Violin plots show the distribution of expression levels across individual nucleus. Pairwise Wilcoxon rank-sum tests with Benjamini–Hochberg correction, states with different lowercase letter differ significantly (adjusted *p*-value < 0.05). See also Figure S1, S2, and S3.

To directly test this hypothesis, we generated cardiomyocyte–specific *Rnf20* knockout mice (*αMHC-MerCreMer* (*35*); *Rnf20^flox/flox^* (*^33^*), cKO) (Fig. 1C, S2B, S2C, and S2D). Given that cell states are shaped by complex interactions among neighboring cells, we performed single-nucleus RNA-sequencing (snRNA-seq) to resolve the transcriptional dynamics of cardiac cell states following cardiomyocyte-specific RNF20 deficiency (Fig. 1C). Cardiac nuclei of both control (*αMHC-MerCreMer*, CreERT2) and cKO were isolated at 2 weeks post-tamoxifen (wpt) from the left ventricular (LV) tissues, which play a predominant role in cardiac contraction.

After quality control, a total of 17,599 cardiac nuclei were retained for analysis, including 9,582 nuclei from the CreERT2 heart and 8,017 nuclei from the cKO heart (Fig. 1D and S3A). Unsupervised clustering identified 10 transcriptionally distinct clusters, which were identified in both genotypes (Fig. 1D and S3A) and annotated based on the expression of established marker genes (Fig. 1E and S3B). Fibroblasts were identified by expression of *Pdgfra*, *Col1a1*, and *Ddr2*. Endothelial cells (ECs) were identified by *Pecam1* expression and further subdivided into endocardial ECs (*Npr3*), vascular ECs (*Fabp4*), and lymphatic ECs (*Prox1*, *Flt4*, and *Lyve1*). Cardiomyocytes were identified by expression of *Ttn*, *Myh6*, and *Pln*. Macrophages were identified by *Adgre1*. B and T cells were identified by expression of *Ikzf3*, *Lef1*, *Skap1*, *Btla*, and *Pax5*. Pericytes were identified by *Pdgfrb* and *Cspg4*. Epicardial cells were identified by *Tbx18*, *Muc16*, *Pcdh15*, and *Msln*. Smooth muscle cells were identified by *Acta2* and *Myh11*.

Comparison of cell-type composition showed that the cKO heart exhibited an increased fraction of fibroblasts and a reduced fraction of pericytes; however, the cardiac cellular compartments were largely preserved following RNF20 deficiency (Fig. 1F).

The preserved proportion of cardiomyocytes in the cKO heart allowed us to ask whether RNF20 deficiency alters cardiomyocyte at the level of transcriptional state rather than cell abundance (Fig. 1F). Subclustering of cardiomyocytes resolved five cell states (CM0-CM4) (Fig. 1G). CM1, CM2, and CM3 were enriched in the CreERT2 heart wile CM0 and CM4 were enriched in the cKO heart (Fig. 1G) suggesting a cell-state transition. Correlation analysis based on highly variable genes revealed a high degree of transcriptional similarity among CM1, CM2, and CM3 (Fig. S3C), indicating that these states represent closely related cardiomyocyte populations. Despite this overall similarity, differential gene expression analysis revealed distinct features among these states (Fig. S3D). CM1 was characterized by higher expression of genes involved in fatty acid β-oxidation (e.g., *Acca2* and *Cpt2*), CM2 showed enrichment of genes associated with transcriptional regulation (e.g., *Ezh2* and *Pcgf3*), and CM3 exhibited increased expression of genes related to myofibril assembly (e.g., *Actc1* and *Csrp3*) (Fig. S3D). Together, these findings reveal functional heterogeneity among homeostatic cardiomyocytes, with distinct metabolic, regulatory, and structural programs coexisting within the adult myocardium.

We next asked whether cKO-enriched states represented less mature populations. Applying CytoTRACE, a potency-scoring method that estimates cellular differentiation status based on transcriptional diversity (*36*), to our cardiomyocyte population, we found that cKO-enriched cell states CM0 and CM4 exhibited higher CytoTRACE scores (Fig. 1H), suggesting a less differentiated state. Consistent with this observation, AUCell analysis (*37, 38*) showed reduced enrichment of the mature cardiomyocyte gene set and increased enrichment of the immature gene set in CM0 and CM4 (Fig. 1I). Together, these results indicate that continuous supply of RNF20 is required to stabilize the mature cardiomyocyte transcriptional state, and that RNF20 deficiency drives cardiomyocytes toward dedifferentiated states.

### RNF20 maintains the structural characteristics of mature cardiomyocytes

Given that RNF20-deficient cardiomyocytes shifted toward transcriptionally less mature states, we next asked whether this state transition is accompanied by alterations in canonical molecular and structural features of mature cardiomyocytes. We first examined the expression of two well-established pairs of cardiomyocyte maturation markers: *Myh6*/*Myh7* and *Tnni3*/*Tnni1*. *Myh7* and *Tnni1* represent the predominant isoforms expressed in murine neonatal cardiomyocytes, whereas a switch toward the adult-associated isoforms *Myh6* and *Tnni3* is a hallmark of cardiomyocyte maturation (*13, 39*). Real-time quantitative polymerase chain reaction (RT–qPCR) analysis revealed expression of *Myh7* and *Tnni1* was significantly increased, accompanied by a reduction in *Tnni3* expression, compared with CreERT2 hearts (Fig. S4A).

We next examined sarcomere organization, a defining structural feature of mature cardiomyocytes and a hallmark disrupted during dedifferentiation (*40–42*). Immunofluorescence staining for cardiac troponin T (cTnT) revealed well-organized sarcomeric striations in CreERT2 hearts, whereas cKO hearts showed marked disruption of sarcomere architecture (Fig. 2A). To quantitatively assess sarcomere organization, we applied SOTAtool(*40, 43*) to calculate the proportion of images with detectable periodic intensity peaks which reflect organized sarcomere. At 2 wpt, cKO hearts exhibited a significantly higher fraction of disorganized sarcomeres compared with CreERT2 hearts (41% in CreERT2 and 64% in cKO) (Fig. S4B). Notably, this defect became more pronounced by 4 wpt (44% in CreERT2 and 91% in cKO) (Fig. 2B). Consistently, immunofluorescence staining for α-actinin with isolated cardiomyocytes confirmed reduced sarcomere organization in cKO cardiomyocytes (Fig. S4C and S4D). Sarcomere length, however, was preserved (Fig. S4D), indicating disruption of sarcomere organization without gross alteration of sarcomere spacing.

**Figure 2.**
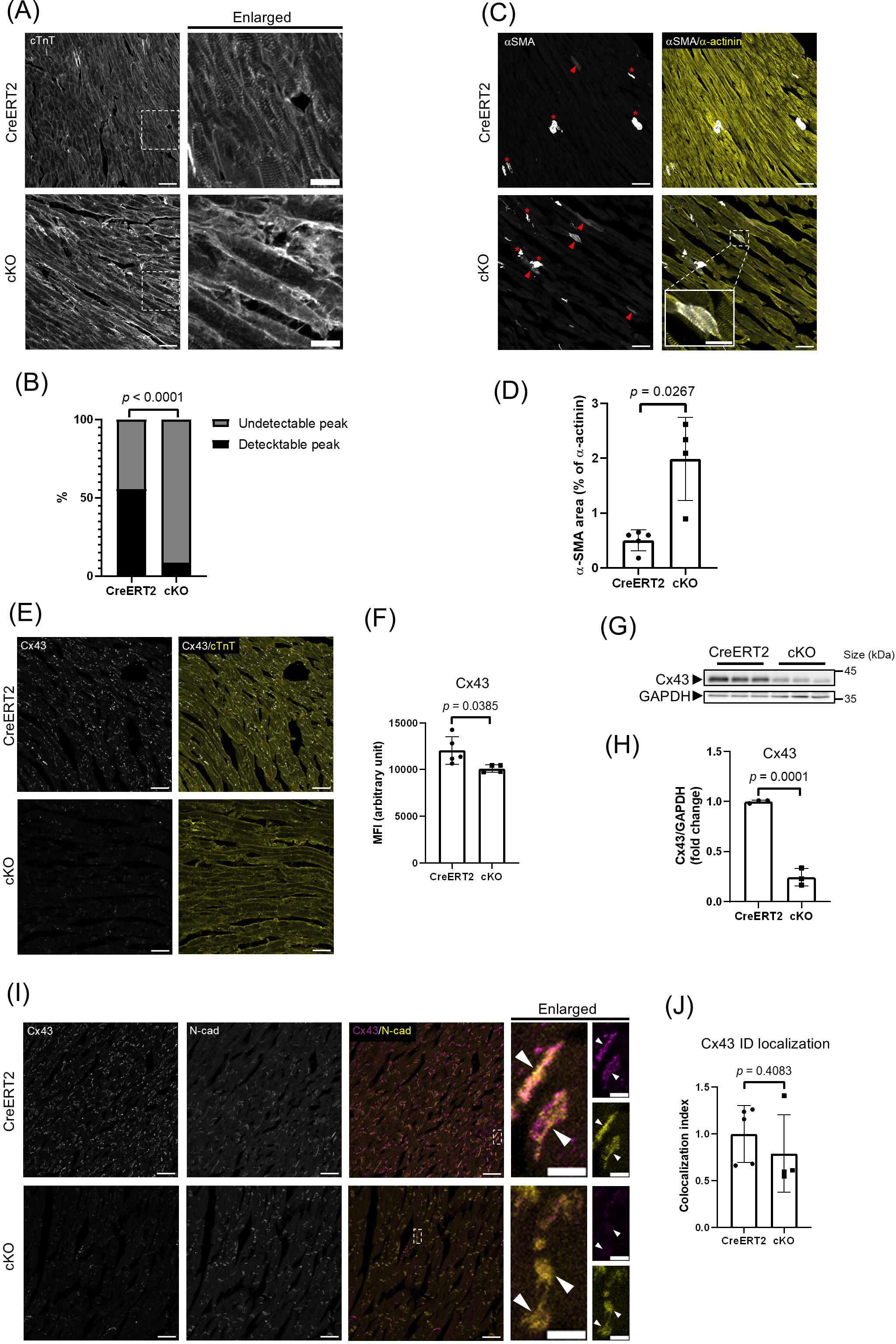
RNF20 maintains the structural characteristics of mature cardiomyocytes. (A) Sarcomere organization evaluation by immunofluorescence analysis of cTnT with tissue sections at 4 wpt. Insets are shown for organized (upper right) and disorganized (lower right) sarcomere. Scale bar = 50 μm (20 μm for insets). (B) Quantification of (A). The data were generated with n = 5 (CreERT2 with 255 sub-images) and 4 (cKO with 238 sub-images) biological replicates. Chi-square test. (C) Immunofluorescence analysis of α-SMA at 4 wpt. Arrows show α-SMA-positive cardiomyocytes and asterisks show vessels. Scale bar = 50 μm (20 μm for insets). (D) Quantification of (C). The data were generated with n = 5 (CreERT2) and 4 (cKO) biological replicates and are presented as mean ± s.d.. Unpaired two-tailed t test with Welch’s correction. (E) Immunofluorescence analysis of Cx43 at 4 wpt. Scale bar = 50 μm. (F) Quantification of (E). MFI, mean fluorescent intensity. The data were generated with n = 5 (CreERT2) and 4 (cKO) biological replicates and are presented as mean ± s.d.. Unpaired two-tailed t test. (G) Western blot analysis of Cx43 from LV tissues at 4 wpt. GAPDH was used as normalization control. (H) Quantification of (G). The data were generated with n = 3 biological replicates and are presented as mean ± s.d.. Unpaired two-tailed t test. (I) Localization of Cx43 at ID was evaluated by co-immunostaining of Cx43 and N-cadherin (N-cad). Arrows show Cx43 localizing ID. Scale bar = 50 μm (10 μm for insets). (J) Quantification of (I). The data were generated with n = 5 (CreERT2) and 4 (cKO) biological replicates and are presented as mean ± s.d.. Unpaired two-tailed t test. See also Figure S4.

Re-expression of α–smooth muscle actin (α-SMA), normally restricted to smooth muscle cells and embryonic cardiomyocytes (*16, 44*), is a hallmark of cardiomyocyte dedifferentiation (*16, 45*). In CreERT2 hearts, α-SMA signal was largely confined to vascular structures and was minimal in cardiomyocytes (Fig. 2C and 2D). In contrast, cKO hearts showed marked induction of α-SMA within cardiomyocytes (Fig. 2C and 2D), with an even stronger signal at 6 wpt (Fig. S4E). These findings indicate that RNF20 deficiency promotes progressive cytoskeletal remodeling and sarcomere disorganization.

In adult cardiomyocytes, the major gap junction protein connexin 43 (Cx43) is concentrated at intercalated discs (ID), where it facilitates intercellular electrical coupling and synchronized contraction (*46*). Reduced expression of Cx43 has been reported in dedifferentiated cardiomyocytes (*47*). Consistent with this, both immunofluorescence staining (Fig. 2E and 2F) and Western blot analysis (Fig. 2G, 2H, S4F, and S4G) revealed a significant reduction in Cx43 expression in cKO hearts. Notably, despite the overall reduction in Cx43 abundance, its localization at ID—identified by N-cadherin staining—was preserved in cKO hearts (Fig. 2I and 2J), indicating that RNF20 preferentially affects Cx43 expression rather than its targeting to ID.

Together, these validation experiments show that the transcriptional shift detected by snRNA-seq is accompanied by not only molecular but also structural regression of adult cardiomyocytes, indicating coordinated loss of mature-state features rather than an isolated transcriptional response. RNF20-deficient cardiomyocytes re-express immature isoforms, lose organized sarcomere architecture, induce α-SMA, and reduce Cx43 expression.

### RNF20 deficiency drives a trajectory away from the mature state

We next sought to order the transcriptional states associated with cardiomyocyte dedifferentiation by performing pseudotime analysis using Monocle3 (*48*) (Fig. 3A). CM2 was designated as the root state, as it showed the lowest CytoTRACE score and was predicted to represent the most differentiated population (Fig. 1H). Consistent with the CytoTRACE analysis (Fig. 1H), the CreERT2-enriched states CM1–CM3 occupied early positions along the trajectory, whereas the cKO-enriched states CM0 and CM4 were positioned later, with CM4 located at the terminal end (Fig. 3A).

**Figure 3.**
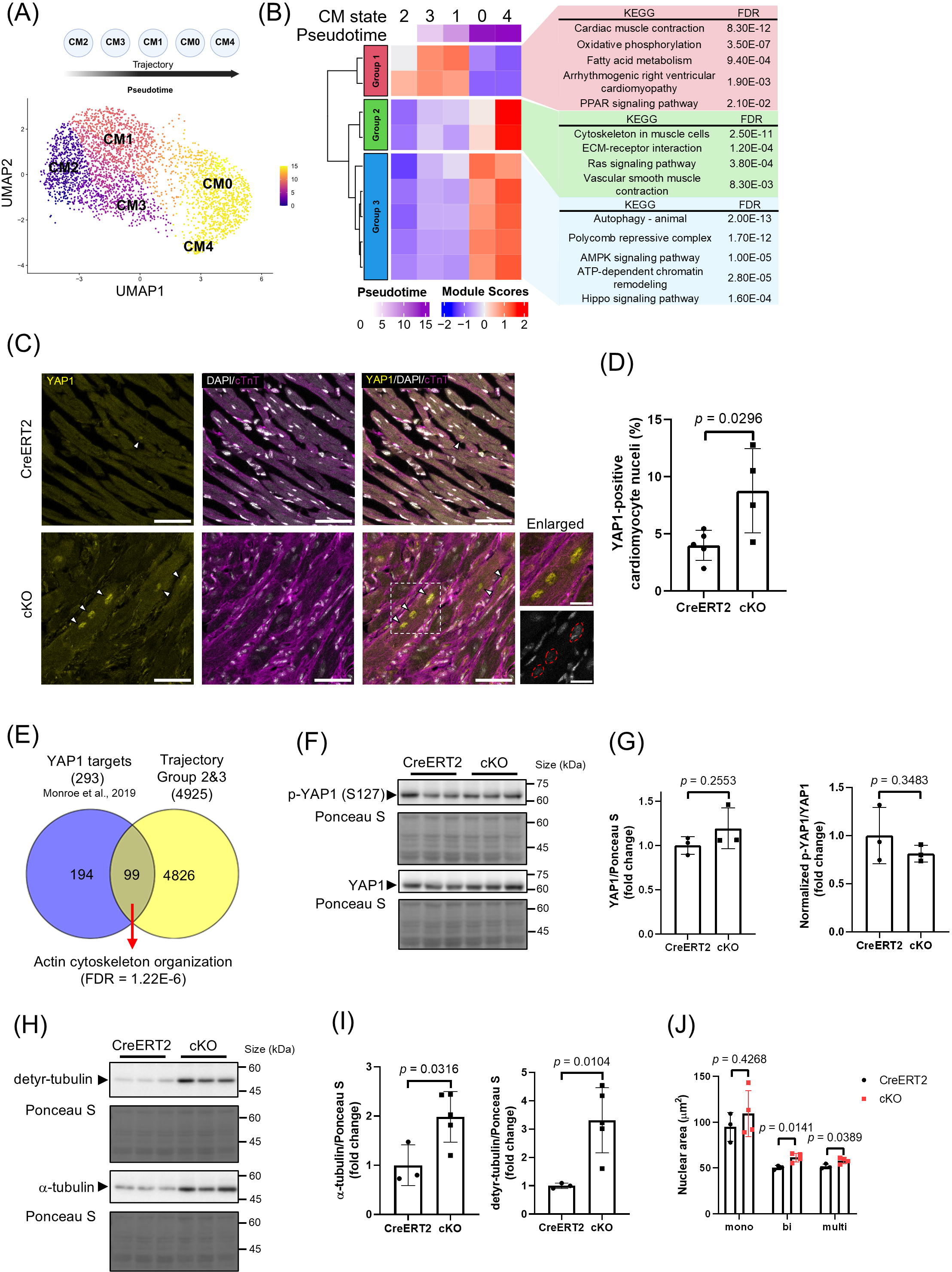
Dedifferentiation follows a stepwise transcriptional trajectory with activation of YAP1 signaling. (A) UMAP visualization of cardiomyocytes colored by pseudotime. (B) Heatmap showing pseudotime-dependent genes (left) and their corresponding overrepresented KEGG pathways. (C) Immunofluorescence analysis of YAP1 at 4 wpt. Arrows show nuclear YAP1. Cardiomyocyte nuclei are encircled in the insets for better identification. Scale bar = 50 μm (20 μm for insets). DAPI, 4’,6-diamidino-2-phenylindole. (D) Quantification of nuclear YAP1 from (C). Data were generated with n = 5 (CreERT2) and 4 (cKO) biological replicates and are presented as mean ± s.d.. Unpaired two-tailed t test. (E) Venn diagram (*117*) showing intersection of YAP1 targets and upregulated genes in RNF20-deficient cardiomyocytes derived from pseudotime analysis. Overrepresentation analysis of upregulated YAP1 targets showed actin cytoskeleton organization as the only significant term (FDR < 0.05) from the Gene Ontology: Biological Process (GO:BP) terms. (F) Western blot analysis of YAP1 and p-YAP1 from LV tissues at 4 wpt. Total protein stained with Ponceau S was used as normalization control. (G) Quantification of YAP1 and p-YAP1 from (F). Data were generated with n = 3 biological replicates and are presented as mean ± s.d.. Unpaired two-tailed t test. (H) Western blot analysis of α-tubulin and detyrosinated-tubulin (detyr-tubulin) from LV tissues at 4 wpt. Total protein stained with Ponceau S was used as normalization control. (I) Quantification of α-tubulin and detyrosinated-tubulin from (H). Data were generated with n = 3 (CreERT2) and 5 (cKO) biological replicates and are presented as mean ± s.d.. Unpaired two-tailed t tests. Welch’s correction was performed for detyr-tubulin. (J) Nuclear area measured with isolated cardiomyocytes at 4 wpt. Data were generated with n = 3 (CreERT2, each with 16-20, 208-291, and 28-94 nuclei from mononucleated, binucleated, and multinucleated cardiomyocytes, respectively) and 4 (cKO, each with 18-21, 151-288, 48-116 nuclei from mononucleated, binucleated, and multinucleated cardiomyocytes, respectively) biological replicates and are presented as mean ± s.d.. Unpaired two-tailed t test. See also Figure S5.

To identify transcriptional programs associated with this transition, we applied Moran’s I spatial autocorrelation analysis to identify genes whose expression varied along pseudotime (*48*). Co-regulated genes were first grouped into modules and were subsequently clustered into three major gene groups (Fig. 3B). Group 1, comprising genes whose expression decreased with increasing pseudotime, was enriched for mature cardiomyocyte functional programs, including cardiac muscle contraction and oxidative phosphorylation (Fig. 3B).

In contrast, Groups 2 and 3 contained genes that were activated as cardiomyocytes deviated from the mature state (Fig. 3B). Group 3 showed increased expression beginning at intermediate pseudotime, particularly in CM0, and was enriched for stress-responsive signaling pathways—including autophagy, AMPK signaling pathway, and Hippo signaling pathway—together with transcriptional regulatory processes, such as ATP-dependent chromatin remodeling (Fig. 3B). Group 2 showed the highest expression at later pseudotime, prominently in CM4, and was enriched for pathways related to cell–extracellular matrix (ECM) interactions and cytoskeletal organization (Fig. 3B). Together, these results define an inferred transcriptional trajectory away from the mature cardiomyocyte state, characterized by decreasing mature cardiomyocyte contractile and metabolic programs, increased stress-responsive and chromatin-regulatory pathways at intermediate pseudotime, and increased cell–ECM interactions and cytoskeletal remodeling at later pseudotime.

### RNF20 deficiency induces a selective YAP1-associated response

Sarcomere disorganization (Fig. 2A, 2B, S4B, S4C, and S4D), reduced expression of Cx43 (Fig. 2E, 2F, 2G, 2H, S4F, and S4G), and activation of cytoskeletal remodeling program (Fig. 2C, 2D, 3B, and S4E) suggested altered cellular architecture and mechanotransductive signaling in RNF20-deficient cardiomyocytes. Given that YAP1, a downstream regulator of Hippo signaling pathway, functions as a mechanosensitive transcriptional coactivator (*49*) and has been implicated in cytoskeletal remodeling, dedifferentiation, and regeneration in cardiomyocytes (*47, 50–53*), we next examined whether YAP1 was associated with the transition away from the mature cardiomyocyte state.

Immunofluorescence analysis revealed increased nuclear YAP1 localization in cKO cardiomyocytes as early as 2 wpt (Fig. S5A and S5B), and this increase persisted at 4 wpt (Fig. 3C and 3D). To determine whether nuclear YAP1 localization was associated with activation of YAP1-dependent transcription, we intersected previously defined YAP1 target genes (*51*) with pseudotime-dependent genes activated during dedifferentiation, corresponding to Groups 2 and 3 from Fig. 3B. Approximately one third of YAP1 target genes were upregulated along the dedifferentiation trajectory (Fig. 3E). Among these activated targets, actin cytoskeleton organization was the only significantly enriched term (false discovery rate, FDR < 0.05), including genes such as *Rock2, Actn1*, and *Fmn1* (Fig. 3E and S5C). Thus, RNF20 deficiency is associated with selective induction of a YAP1-linked cytoskeletal organization program rather than broad activation of YAP1 target genes.

Since canonical Hippo signaling restricts YAP1 nuclear localization through Ser127 phosphorylation (*54*), we next examined total and phosphorylated YAP1 levels. Despite increased nuclear localization of YAP1 (Fig. 3C and 3D), neither total YAP1 abundance nor Ser127 phosphorylation was significantly altered between genotypes (Fig. 3F and 3G). We then assessed cytoskeletal features that can influence YAP1 localization independently of canonical Hippo signaling (*49, 55*). cKO hearts showed increased levels of both total α-tubulin and detyrosinated tubulin (Fig. 3H and 3I), consistent with microtubule remodeling (*47, 56*). Isolated cKO cardiomyocytes also exhibited increased nuclear area (*47, 55, 57*) compared with CreERT2 controls (Fig. 3J). Taken together, these findings associate nuclear YAP1 accumulation to cytoskeletal remodeling and altered nuclear morphology in RNF20-deficient cardiomyocytes.

### Mature state destabilization is accompanied by an injury-like state but not cell-cycle re-entry

Because cardiomyocyte dedifferentiation (*15*) and YAP1 activation (*50, 51*) have been linked to regenerative proliferation, we next sought to determine whether RNF20-deficient cardiomyocytes acquire a regeneration-associated state. Comparison with published snRNA-seq dataset from regenerative postnatal day 1 and non-regenerative postnatal day 8 injured murine hearts (*58*) (Fig. 4A and 4B) revealed that the least mature cardiomyocyte population in our dataset, CM4, was more similar to the injury-associated nCM5 state than to the regeneration-competent nCM4 state (Fig. 4C).

**Figure 4.**
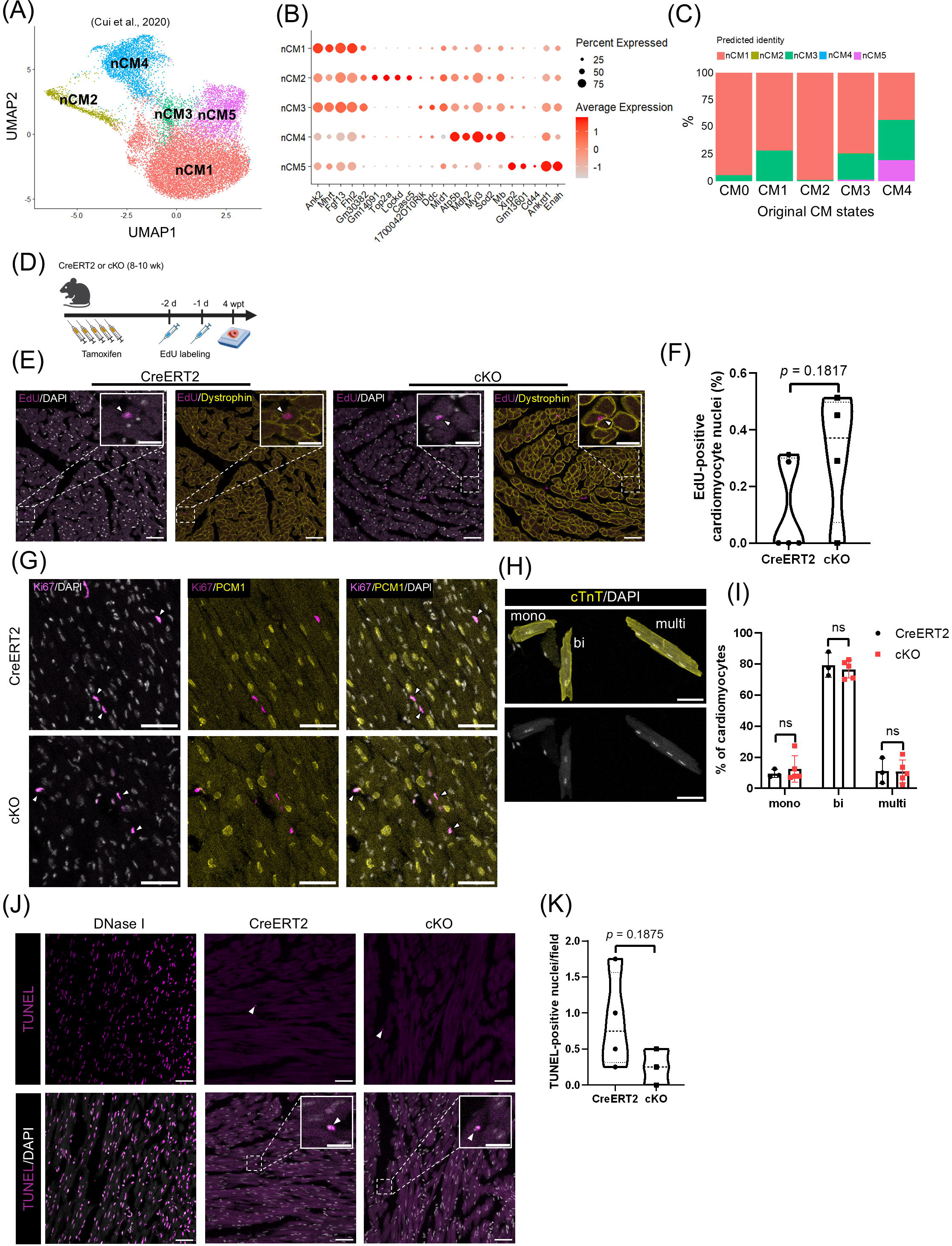
Mature state destabilization is accompanied by an injury-like state but not cell-cycle re-entry. (A) UMAP visualization of neonatal cardiomyocytes post-myocardial infarction. (B) Dot plot of expression of pre-defined marker genes. (C) Unbiased label transfer from reference cells (nCM1-nCM5) to query cells (CM0-CM4). (D) Workflow of EdU incorporation assay. (E) EdU incorporation assay at 4 wpt. Arrows show EdU-positive cardiomyocyte nuclei. Scale bar = 50 μm (20 μm for insets). (F) Quantification of (E). Data were generated with n = 5 (CreERT2) and 4 (cKO) biological replicates and are presented as mean ± s.d.. Unpaired two-tailed t test. (G) Immunofluorescence analysis of Ki67 at 4 wpt. Arrows show Ki67-positive nonmyocyte nuclei. Scale bar = 50 μm. (H) Nucleation status evaluation with isolated cardiomyocytes at 4 wpt. Mono, bi, and multi indicates cardiomyocytes with 1, 2, and ≥ 3 nuclei, respectively. Scale bar = 50 μm. (I) Quantification of (H). ns, nonsignificant. *p* = 0.591 (mono), 0.5963 (bi), and 0.9566 (multi). Data were generated with n = 3 (CreERT2, each with 212-330 cells) and 5 (cKO, each with 241-351 cells) biological replicates and are presented as mean ± s.d.. Unpaired two-tailed t test. (J) TUNEL assay at 4 wpt. The section treated with DNase I was used as a positive control. Arrows show TUNEL-positive nuclei. Scale bar = 50 μm (20 μm for insets). (K) Quantification of TUNEL-positive nuclei from (J). Violin plots show the distribution of individual data points. Data were generated with n = 4 (CreERT2) and 3 (cKO) biological replicates and are presented as mean ± s.d.. Unpaired two-tailed t test.

Consistent with this observation, neither 5-Ethynyl-2’-deoxyuridine (EdU) incorporation assay (Fig. 4D, 4E, and 4F) nor Ki67 staining (Fig. 4G) detected increased cardiomyocyte proliferation in cKO hearts, and the proportion of mononucleated cardiomyocytes (*59*) remained unchanged (Fig. 4H and 4I). Since nCM5 has been linked to apoptotic signaling after injury (*58*), we next assessed cell death by terminal deoxynucleotidyl transferase dUTP nick end labeling (TUNEL) assay. However, no significant increase in TUNEL-positive nuclei was found in cKO hearts (Fig. 4J and 4K). These results indicate that destabilization of the mature state redirects cardiomyocytes toward an injury-like state without productive cell-cycle re-entry or increased apoptosis, rather than restoring a regeneration-competent neonatal state.

### RNF20 maintains the mature chromatin landscape by preserving promoter accessibility and restraining AP-1-associated enhancers

Having defined the transcriptional features of mature-state erosion, we next asked whether the state transition was reflected in the cardiomyocyte cis-regulatory landscape. To address this, we performed assay for transposase-accessible chromatin with sequencing (ATAC-seq) on fluorescence-activated cell sorting-purified PCM1-positive cardiomyocyte nuclei (*12, 60*) (Fig. 5A), generating a consensus set of 48,796 accessible peaks (Fig. S6A).

**Figure 5.**
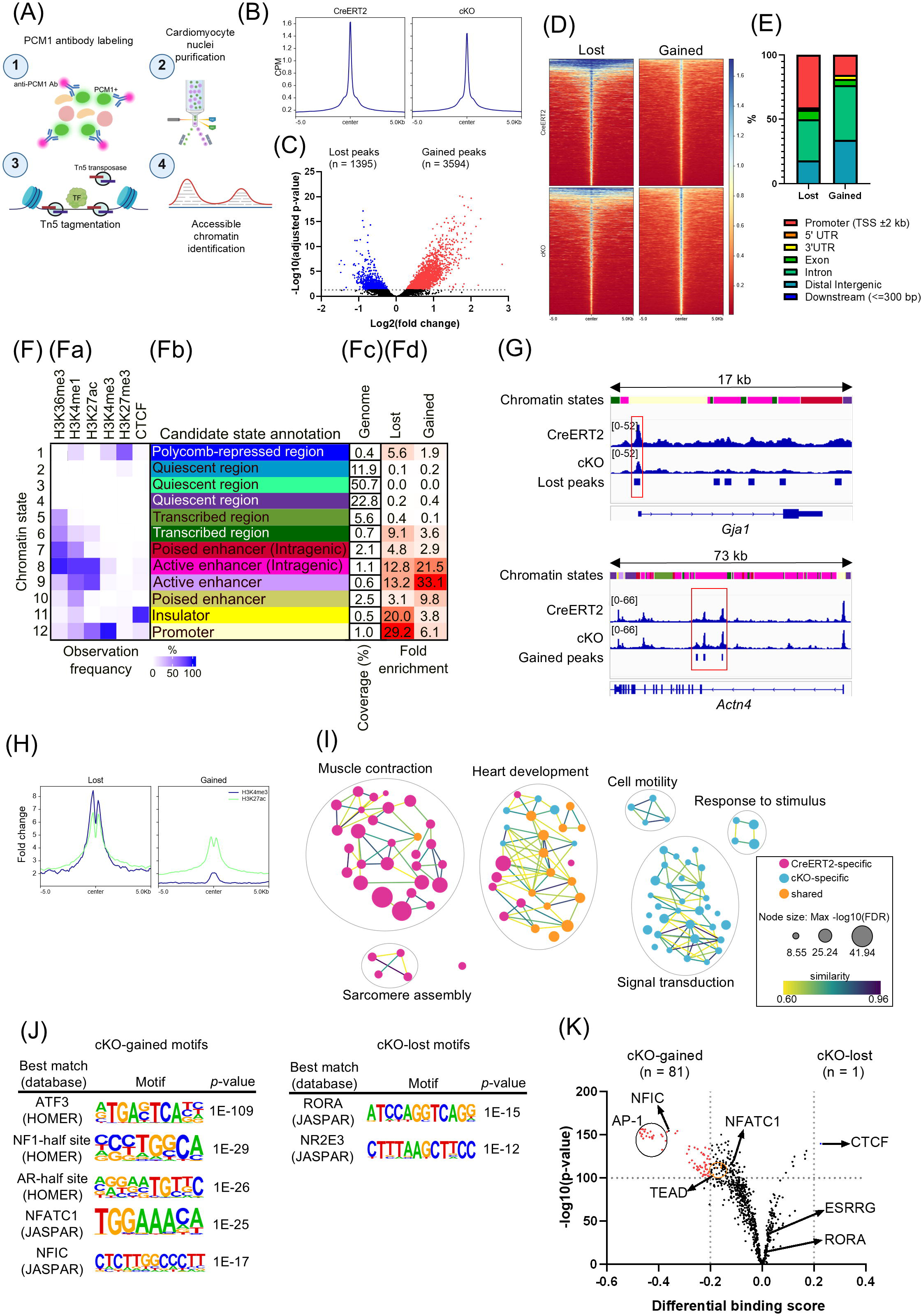
RNF20 maintains the mature chromatin landscape by preserving promoter accessibility and restraining AP-1-associated enhancers. (A) Workflow for ATAC-seq experiment. (B) Metagene profile plots of ATAC-seq signals centered on accessible peaks. CPM, counts per million. (C) Volcano plot showing differentially accessible peaks (adjusted *p*-value < 0.05). (D) Heatmaps showing ATAC-seq signals centered on peaks which lost accessibility (left) and gained accessibility (right) in RNF20-deficient cardiomyocytes. (E) Genomic annotation of differentially accessible peaks. (F) Chromatin state annotation and enrichment analysis. (**Fa**) Observation frequency of chromatin states. Color intensity indicates the frequency of each mark within each chromatin state. (**Fb**) Candidate chromatin state annotation based on combinatorial patterns of epigenetic marks. (**Fc**) Genomic coverage of each chromatin state. (**Fd**) Fold enrichment represents the enrichment of each chromatin state in lost or gained regions relative to genomic background. Color intensity indicates the degree of enrichment. (G) Tracks of ATAC-seq signals in regions showing lost (top) and gained (bottom) chromatin accessibility. Red boxes highlight promoter-associated regions (top) or enhancer-associated regions (bottom). Chromatin state colors correspond to the annotations shown in (Fig. 5Fb). (H) Metagene profile plots of H3K4me3 or H3K27ac ChIP-seq signals centered on differentially accessible peaks. (I) Network graph of enriched GO:BP terms associated with altered chromatin accessibility. Nodes represent terms, colored by enrichment in lost-accessibility regions (red), gained-accessibility regions (cyan), or both (yellow). Node size reflects enrichment significance (FDR). Edges indicate semantic similarity. Functionally related clusters are circled and labeled. (J) De novo motif enrichment analysis. Motifs with *p* ≦ 1 × 10^⁻12^ were considered significant. (K) Footprinting analysis. TF with differential binding activity (*p*-value < 1×10^-100^ and differential binding score > ∣0.2∣) were considered significant and colored with red (upregulated) or blue (downregulated). See also Figure S6.

Global accessibility profiles were broadly comparable between CreERT2 and cKO cardiomyocytes (Fig. 5B); however, differential accessibility analysis identified 1,395 regions losing and 3,594 regions gaining accessibility in RNF20-deficient cardiomyocytes (adjusted *p*-value< 0.05) (Fig. 5C and 5D).

Genomic annotation revealed that promoter-proximal regions were enriched among peaks with decreased accessibility, but not among regions with increased accessibility (40.7% versus 15.4%; Fig. 5E). ChromHMM (*61*) modeling based on multiple epigenetic marks from the ENCODE project (*62–64*) demonstrated that regions with decreased accessibility were predominantly enriched in trimethylation of histone H3 at lysine 4 (H3K4me3)-marked promoter states, whereas regions with increased accessibility were primarily enriched in acetylation of histone H3 at lysine 27 (H3K27ac)-marked active enhancer states (Fig. 5F and 5G). Published histone-mark profiles from isolated cardiomyocyte nuclei (*12*) further supported these annotations: decreased-accessibility regions showed strong signals of H3K4me3 and H3K27ac, while increased-accessibility regions showed higher signal of H3K27ac relative to H3K4me3 (Fig. 5H).

Region-based functional enrichment analysis (*65*), with enriched terms clustered by semantic similarity (*66*), linked decreased-accessibility regions to muscle contraction and sarcomere assembly, whereas increased-accessibility regions to stimulus, signal transduction, and cell motility (Fig. 5I and Data S2). These functions matched the pseudotime-defined dedifferentiation programs (Fig. 3B). Consistently, downregulated Group 1 genes accounted for a greater proportion of genes associated with regions exhibiting reduced chromatin accessibility, whereas upregulated Group 2 and Group 3 genes accounted for a greater proportion of genes associated with regions exhibiting increased accessibility (Fig. S6C).

These findings identify a RNF20-dependent cis-regulatory landscape that stabilize the mature cardiomyocyte state by preserving its identity-associated regulatory architecture while limiting access to alternative stress-responsive programs. Disruption of RNF20 reduces accessibility at promoter regions associated with contractile and sarcomeric programs while increases accessibility at enhancer regions linked to stress-responsive and cytoskeletal remodeling programs.

Given that cardiomyocyte dedifferentiation is accompanied by increased accessibility at enhancer-associated regions (Fig. 5F and Fig. 5H), we next asked which TF networks were linked to this altered-accessibility landscape. Motif enrichment analysis identified AP-1 (ATF3) as the most significantly enriched motif among regions with increased accessibility (*p* = 1 × 10⁻¹⁰⁹), followed by motifs corresponding to members of the NF1, NFAT, and TEAD (AR-half site) (Fig. 5J). In contrast, regions with decreased accessibility showed fewer and weaker motif enrichments, with RORA representing the top motif in this group (*p* = 1 × 10⁻¹⁵) (Fig. 5J). To refine the motif analysis and identify TFs engaged, we performed footprinting analysis (*67*). Eighty-one transcription factors exhibited significantly increased binding in cKO cardiomyocytes, whereas only CTCF showed increased binding in CreERT2 cardiomyocytes (Fig. 5K). Among motifs enriched in regions with increased accessibility (Fig. 5J), AP-1 family members and NFIC displayed increased binding, whereas TEAD and NFAT family members did not show significant footprinting signals (Fig. 5K). Consistently, cKO cardiomyocytes showed increased accessibility at AP-1-related regions that gain accessibility after myocardial infarction, whereas CreERT2 cardiomyocytes retained higher accessibility at injury-lost regions (*68*) (Fig. S6D). Thus, the enhancer-associated accessibility gained after RNF20 deficiency is linked predominantly to AP-1-associated stress-responsive regulatory activity.

The limited enrichment of TEAD motifs (Fig. 5J) suggested that YAP1-associated response following RNF20 depletion is transcriptionally distinct from regenerative YAP1 signaling (*50, 51, 53*). To place this observation into context, we reanalyzed published ATAC-seq data from adult cardiomyocytes expressing constitutively active YAP1 (YAP5SA) (*51*). YAP5SA cardiomyocytes showed robust TEAD motif enrichment (*p* = 1 × 10⁻¹⁴⁴⁰), together with motifs corresponding to additional TFs implicated in cardiomyocyte regeneration, including GATA (*69*) and KLF (*70*) families (Fig. S6E). Among regions with decreased accessibility, YAP5SA cardiomyocytes exhibited broader motif enrichment compared with RNF20-deficient cardiomyocytes, with MEF2 motifs showing the strongest significance (*p* = 1 × 10⁻²⁰⁴) (Fig. S6E). Together, these findings indicate that the regulatory landscape associated with regenerative YAP1 activation was markedly distinct from that of RNF20-deficient cardiomyocytes, the latter being characterized by AP-1-associated stress-responsive enhancer activation with only modest TEAD engagement.

### Stress-responsive cardiomyocyte state preferentially localizes to the subendocardium

The preceding analyses revealed that RNF20 deficiency in cardiomyocytes erodes mature cardiomyocyte state and induces a non-proliferative injury-like state (Fig. 1H, 1I, 3B, and 4). Since snRNA-seq does not preserve information regarding tissue organization, we therefore asked whether destabilization of the mature cardiomyocyte state occurred uniformly across the myocardium or exhibited spatially restricted vulnerability.

We performed high-resolution spatial transcriptomics using the Visium HD platform on formalin-fixed paraffin-embedded hearts collected at 2 wpt (Fig. 6A). Spatial expression of *Dcn* in fibrous connective tissue and *Myh11* in coronary vessels confirms accurate spatial capture of tissue-derived transcripts (Fig. 6B). In addition, the cardiomyocyte marker *Pln* was broadly detected across ventricular myocardium and appeared reduced in the cKO heart (Fig. 6B), consistent with attenuation of mature cardiomyocyte programs.

**Figure 6.**
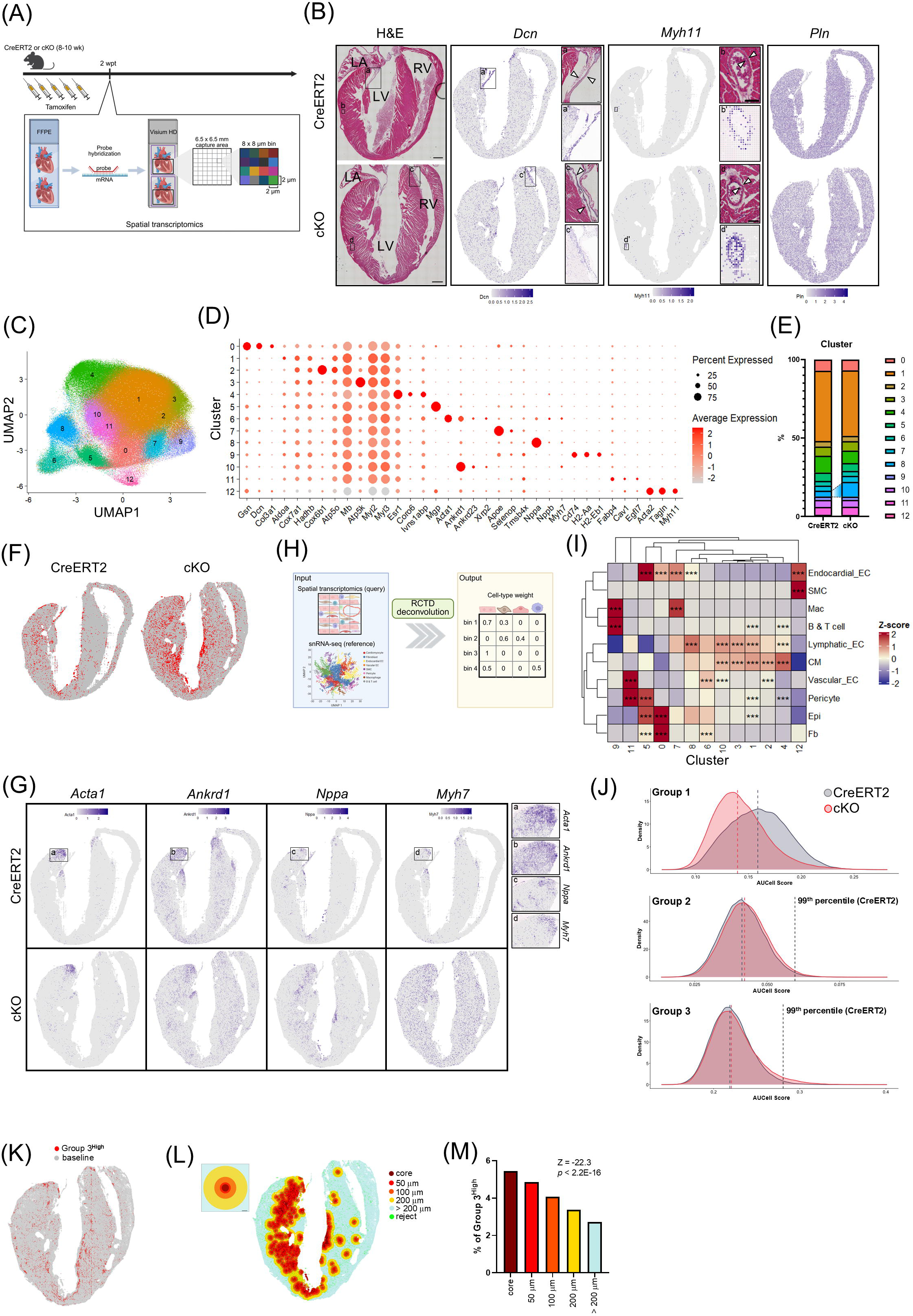
Stress-responsive cardiomyocyte sate preferentially localizes to the subendocardium. (A) Workflow for spatial transcriptomics. (B) Hematoxylin and eosin (H&E) images and spatial expression patterns of marker genes for fibroblasts (*Dcn*), smooth muscle cells (*Myh11*), and cardiomyocytes (*Pln*). Insets show enlarged views of the boxed regions in the corresponding H&E images and spatial maps. LA, left atrium; LV, left ventricle; RV, right ventricle. Arrows show fibrous connective tissue (a and c) or coronary vessels (b and d). Scale bar = 500 μm (50 μm for insets). (C) UMAP visualization of Louvain clusters identified from ventricular 8-µm bins across both genotypes. (D) Dot plot showing differentially expressed genes across clusters. (E) Proportion of clusters. Expansion of cluster 8 is highlighted. (F) Spatial mapping of clusters 8. (G) Spatial feature plots showing regionally enriched expression of mechanical stress-associated genes. Insets show enlarged views of the boxed regions (left atrioventricular plane) in the corresponding spatial maps. (H) Schematic of spatial deconvolution analysis. (I) Heatmap displaying the mean enrichment of cell types within each cluster. EC, endothelial cell; SMC, smooth muscle cell; Mac, macrophage; CM, cardiomyocyte; Epi, epicardial cell; Fb, fibroblast. Asterisks indicate significant enrichment of a cell type within a cluster compared to the remainder of the tissue, determined by one-sided Wilcoxon rank-sum tests (alternative = “greater”) with Bonferroni correction (***, adjusted *p*-value< 0.001). Color scale represents the z-score of cell-type weights. (J) Density plots showing AUCell scores for pseudotime-dependent genes (Fig. 3B) among cardiomyocyte-enriched bins (cardiomyocyte weight > 0.8). Grey, red, and black vertical dashed lines indicate the median value for CreERT2, median value for cKO, and 99^th^ percentile for CreERT2, respectively. The fractions of Group 2 high-scoring bins and Group 3 high-scoring bins (> 99th percentile for CreERT2) in the cKO heart were 1.58% and 3.50%, respectively. (K) Spatial mapping of Group 3^High^ in the cKO heart. (L) Left: Schematic of distance-based spatial stratification. Bins were categorized according to Euclidean distance from the cluster 8 epicenter: core (dark red), 0–50 µm (red), 50–100 µm (orange), 100–200 µm (yellow), and >200 µm (light blue). Scale bar = 50 µm. Right: Spatial mapping of distance-resolved zones in the cKO heart. Green spots indicate sparse cluster 8 bins excluded from the core because of insufficient neighboring cluster 8 bins (n < 25). (M) Percentage of Group 3^High^ across distance-resolved zones in the cKO heart. Cochran-Armitage test for trend. See also Figure S7, S8, S9, and S10.

Unsupervised clustering identified 13 spatial clusters shared between genotypes (Fig. 6C and S7), with several of which exhibited distinct spatial distribution and cell-type–associated marker genes (Fig. 6D and S7). Analysis of cluster composition revealed that most clusters remained relatively stable between genotypes, whereas cluster 8 increased from 3.7% in CreERT2 to 9.4% in cKO (Fig. 6E). Cluster 8 was enriched in the subendocardial region and showed elevated expression of the hypertrophic marker *Nppa* (Fig. 6D and 6F). Spatial mapping of mechanical stress-associated genes further showed that some transcripts, including *Acta1* and *Ankrd1*, remained restricted to the left atrioventricular plane and papillary muscle in both genotypes, whereas others, such as *Nppa*, expanded into subendocardial regions or became broadly distributed throughout the ventricular myocardium (*Myh7*) in the cKO heart (Fig. 6G).

To relate these clusters to cell identity, we performed spatial deconvolution using our snRNA-seq data as reference (Fig. 6H and S8) (*71*). The inferred cell-type composition (Fig. 6I) closely matched marker gene–based cluster annotations (Fig. 6D). Notably, cluster 8 exhibited enrichment of endocardial EC (Fig. 6I), consistent with its subendocardial localization (Fig. 6F).

We next projected the snRNA-seq-derived dedifferentiation programs onto spatial bins (Fig. 3B and S9). Consistent with the dedifferentiation trajectory, enrichment scores for Group 1 mature cardiomyocyte functional programs were globally reduced in cKO, with median scores decreasing from 0.16 in CreERT2 to 0.14 in cKO (Fig. 6J). In contrast, median scores for Group 2 cytoskeletal remodeling and Group 3 stress-responsive programs remained largely unchanged, whereas the high-scoring bins increased in the cKO heart (Fig. 6J). Together, these findings recapitulate the dedifferentiation trajectory inferred from snRNA-seq (Fig. 3B), with widespread reduction of mature cardiomyocyte programs and gradual emergence of stress-responsive and cytoskeletal remodeling programs.

Spatial mapping of Group 3-high-scoring cardiomyocyte-enriched bins in the cKO heart (Group 3^High^) (Fig. 6K) revealed a distribution pattern similar to the expanded subendocardial cluster 8 (Fig. 6F). To determine whether Group 3^High^ bins were preferentially localized within cluster 8 in the cKO heart, we performed distance-based analysis centered on cluster 8 regions (Fig. 6L). The fraction of Group 3^High^ bins was highest within the cluster 8 core and progressively decreased with increasing distance from the core (Cochran–Armitage trend test, *p* < 2.2 × 10^⁻16^) (Fig. 6M). These findings suggest that activation of the stress-responsive transcriptional programs preferentially occur within spatially restricted subendocardial niche.

Although fibroblast expansion and activation were observed in snRNA-seq (Fig. 1F, S10A, S10B, and S10C), spatial transcriptomics did not detect prominent changes in fibroblasts in the analyzed cKO heart section (Fig. S8B). To further assess fibroblast activation in cKO hearts, we examined expression of activated fibroblast marker genes using RT-qPCR. Notably, expression levels of these markers inhomogeneously increased in cKO hearts and positively correlated with the dedifferentiation marker gene *Tnni1* (Fig. S10D and S10E). Together with the selective induction of mechanical stress-associated genes observed in spatial transcriptomics (Fig. 6G), these findings suggest that fibroblast activation is coupled to the extent of mature cardiomyocyte state regression, rather than occurring synchronously across cKO hearts at this stage.

Collectively, these observations indicate that loss of mature-state stability is spatially nonuniform. Instead, mature cardiomyocyte programs are broadly attenuated, whereas the stress-responsive programs, defined by the Group 3 trajectory modules, preferentially localize within the subendocardial niche.

### Mature-state erosion progresses to cardiomyopathy

Given the extensive transcriptional and structural remodeling observed in RNF20-deficient cardiomyocytes, we next examined the functional consequences of RNF20 depletion at the organ-level (Fig. 7A). Longitudinal assessment of cardiac function revealed progressive deterioration in both systolic performance and ventricular conduction in cKO mice, with conduction abnormalities preceding evident systolic dysfunction (Fig. 7B and 7C). QRS interval prolonged as early as 2 wpt and continued to lengthen over time (Fig. 7C). Notably, this early conduction defect coincided with reduced expression of Cx43 (Fig. S4F and S4G), consistent with impaired electrical coupling between cardiomyocytes (*46*). Echocardiographic analysis further demonstrated structural remodeling, including increased LV mass-to-body-weight ratio (Fig. 7D) and LV dilation, reflected by increased LV internal diameter (Fig. 7E and S11A), without appreciable changes in wall thickness (Fig. S11B).

**Figure 7.**
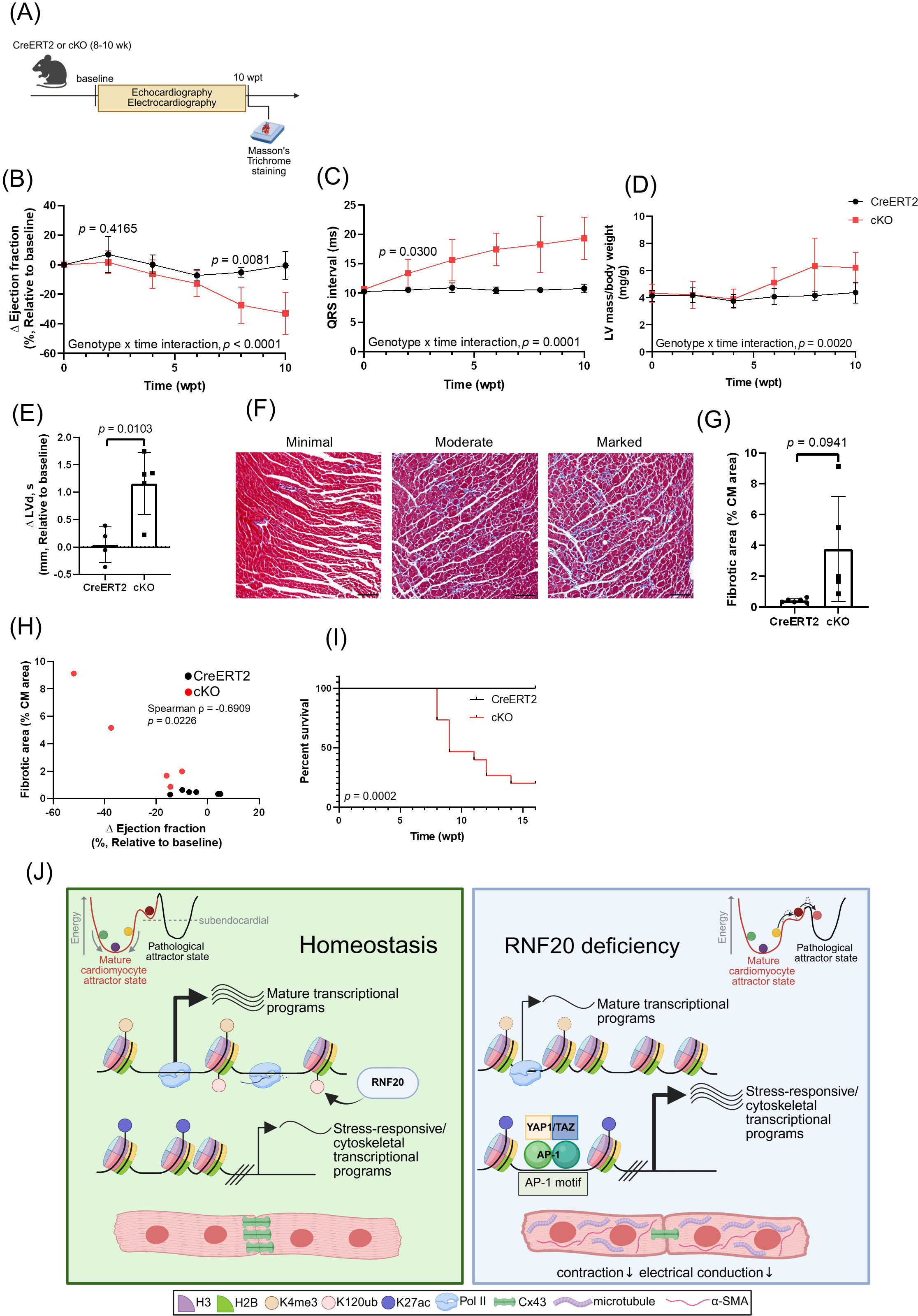
Mature-state erosion progresses to cardiomyopathy. (A) Experimental design for assessment of cardiac function. (B) Change in ejection fraction relative to baseline. Data were generated with n = 4 (CreERT2), 6 (cKO, 0-8 wpt), and 5 (cKO, 10 wpt) biological replicates and are presented as mean ± s.d.. The genotype × time interaction was examined using a mixed-effects model, whereas differences between groups at the indicated time points using unpaired two-tailed t tests. (C) QRS interval measured by electrocardiography. Data were generated with n = 4 (CreERT2), 6 (cKO, 0-8 wpt), and 5 (cKO, 10 wpt) biological replicates and are presented as mean ± s.d.. The genotype × time interaction was examined using a mixed-effects model, whereas difference between groups at the indicated time point using unpaired two-tailed t tests with Welch’s correction. (D) Ratio of LV mass-to-body-weight. LV mass was measured with echocardiography. Data were generated with n = 4 (CreERT2), 6 (cKO, 0-8 wpt), and 5 (cKO, 10 wpt) biological replicates and are presented as mean ± s.d.. The genotype × time interaction was examined using a mixed-effects model. (E) Change in LV end-systolic internal diameter (LVd, s) at 10 wpt relative to baseline. Data were generated with n = 4 (CreERT2) and 5 (cKO) biological replicates and are presented as mean ± s.d.. Unpaired two-tailed t test. (F) Ventricular fibrosis determined with Masson’s Trichrome staining. Images represent minimal (left, 0.4% fibrotic area), moderate (middle, 3.7% fibrotic area), and marked (right, 7.9% fibrotic area) fibrosis. Scale bar = 100 μm. (G) Quantification of fibrotic area at 10 wpt. CM, cardiomyocyte. Data were generated with n = 6 (CreERT2) and 5 (cKO) biological replicates and are presented as mean ± s.d.. Unpaired two-tailed t test with Welch’s correction. (H) Correlation analysis of fibrotic area and change in ejection fraction. Associations were assessed using Spearman’s rank correlation. (I) Survival rate. Data were generated with n = 10 (CreERT2) and 15 (cKO) biological replicates. Log-rank test. (J) Working model: RNF20-dependent chromatin regulation stabilizes the mature cardiomyocyte attractor state while constraining transitions toward pathological cell states, thereby preserving adult cardiac homeostasis. See also Figure S11.

Histological analysis revealed increased interstitial fibrosis in cKO hearts, with a trend toward increased fibrotic area (Fig. 7F and 7G). Fibrosis severity varied among individual mice and was negatively correlated with the change in ejection fraction (Fig. 7G and 7H), indicating heterogeneity in disease progression. This variability was consistent with the heterogeneous induction of fibroblast activation markers across individual cKO hearts (Fig. S10D and S10E), supporting the idea that non-myocyte remodeling scales with the severity of mature cardiomyocyte state regression.

Consistent with progressive cardiac dysfunction, cKO mice showed a tendency toward reduced body-weight gain over time (Fig. S11C). Survival analysis further revealed premature mortality as 53% of cKO mice died within 9 wpt, and 80% died by 16 wpt, whereas all CreERT2 mice survived throughout the observation period (Fig. 7I).

Collectively, these findings show that failure to maintain the mature cardiomyocyte state progresses from early conduction abnormalities to, systolic dysfunction, ventricular dilation, interstitial fibrosis, and premature death, culminating in a dilated cardiomyopathy-like disease course.

## DISCUSSION

Long-lived postmitotic cells must preserve highly specialized functions while retaining sufficient plasticity to respond to physiological demands and pathological stress (*15, 16, 72, 73*). Although maturation is increasingly viewed as a dynamic adaptive state rather than a fixed developmental endpoint (*11*), how an established mature state is constantly maintained remains poorly understood. Here, we show that the mature cardiomyocyte state requires continuous chromatin-based reinforcement for long-term stability. Acute RNF20 depletion after completion of cardiomyocyte maturation destabilized the mature cardiomyocyte transcriptional state, shifting cells toward heterogeneous injury-associated states. These changes were accompanied by altered chromatin accessibility, nuclear YAP1 accumulation without productive cell proliferation, and emergence of a subendocardial stress-responsive niche. The subsequent fibrosis, conduction and contractile dysfunction, and premature mortality show that loss of mature-state stability propagates from transcriptional instability to organ-level disease. Thus, continuous chromatin maintenance preserves adult cardiomyocyte function not by locking cells into a rigid transcriptional state, but by stabilizing the mature cardiomyocyte attractor state while constraining transitions toward pathological cell states (Fig. 7J).

This requirement for continuous chromatin maintenance distinguishes the establishment of cardiomyocyte maturity from its maintenance. Previous studies showed that RNF20 is indispensable during embryonic cardiogenesis and postnatal cardiomyocyte maturation (*33, 34, 74*). However, a developmental role does not establish that the same mechanism remains equally required after the attainment of maturation (*6*). This distinction is evident in postmitotic neurons: de novo histone H3.3 is essential during a restricted window for establishing neuronal chromatin and identity but becomes substantially less important once that state is consolidated (*75*), whereas Pet-1 remains continuously required in adult serotonin neurons to preserve normal anxiety-related behaviors (*76*). Our adult-inducible model suggests that RNF20 belongs to this continuously required class. Despite declining during postnatal maturation, residual RNF20 activity remains necessary to preserve an established mature cardiomyocyte state. Moreover, RNF20 loss did not simply revert cardiomyocytes to a neonatal transcriptional state. Single-nucleus analysis instead resolved a modular transition in which mature contractile and oxidative phosphorylation programs declined while stress-responsive programs—including autophagy, AMPK-signaling pathway, and chromatin-remodeling machinery—together with cytoskeletal remodeling program, progressively activated. The resulting phenotype is therefore best understood as loss of mature-state fidelity and acquisition of a pathological transcriptional state rather than a faithful return to a regenerative neonatal state.

Unlike mechanisms that stabilize cell identity primarily through transcriptional repression (*22, 77*), RNF20/H2Bub operates within actively transcribed chromatin, providing a distinct mechanism for continuously reinforcing the resident transcriptional state. RNF20-dependent genes are enriched for mature cardiomyocyte functions, carry broader promoter-associated H3K4me3 domains (*12*) (Fig. S12A and S12B), exhibit efficient transcriptional elongation (*78*) (Fig. S12C and S12D), and lose promoter accessibility after RNF20 depletion. Broad H3K4me3 domains are associated with transcriptional consistency at cell-identity genes (*79*), and our reanalysis of human heart-failure cardiomyocytes dataset revealed narrowing of H3K4me3 domains (Fig. S12E and S12F) was preferentially associated with downregulated genes (*80*) (Fig. S12G). Integrating findings from murine homeostatic, RNF20-deficient, and human heart-failure datasets, we propose a model in which RNF20-dependent H2Bub across coding regions sustains transcriptional processivity and chromatin integrity, whereas the downstream establishment of promoter-associated H3K4me3 broadening preserves durable transcriptional competence (*81*). This interpretation is consistent with selective retention of H2Bub on long cardiac genes (*74*), its association with full-length transcript production (*27, 28*), and its ability to recruit histone chaperone FACT to coordinate transcription-coupled nucleosome remodeling and restoration (*29*). Such continuous chromatin servicing may be especially important in postmitotic cardiomyocytes, which repeatedly transcribe identity genes despite the absence of replication-dependent chromatin resetting. At the same time, RNF20 may also reconcile state stability with environmental responsiveness. Stress-responsive genes exhibited substantial gene-body H2Bub (Fig. S13) and promoter-proximal Pol II pausing under homeostatic conditions (Fig. S12C and S12D), and RNF20 can restrain pause release at selected inducible loci (*82–84*). RNF20 may therefore perform complementary functions by continuously reinforcing mature-state transcription while maintaining stress-responsive genes in a poised, inducible configuration.

Failure of this chromatin maintenance system generated pathological rather than regenerative reprogramming. RNF20-deficient cardiomyocytes retained lineage identity while acquiring selective immature features together with injury-associated cytoskeletal, survival-oriented metabolic, and chromatin-remodeling programs. This transition was accompanied by loss of accessibility at promoter-associated mature cardiac elements and gain of accessibility at enhancer-associated regions enriched for AP-1 motif. RNF20-deficient cardiomyocytes also accumulated nuclear YAP1 and underwent extensive cytoskeletal remodeling. However, YAP1-associated transcription was biased toward actin-cytoskeletal genes, whereas canonical cell-cycle targets remained largely unchanged and productive cell-cycle re-entry did not occur. Increased ARID1A and other SWI/SNF components may contribute to this restricted output by limiting YAP1–TEAD activity (*85, 86*) (Fig. S14), while the prominent AP-1 signature raises the possibility that YAP1 is preferentially redirected toward injury-associated rather than proliferative programs (*17, 68, 87, 88*). Our findings therefore demonstrate that mature-state regression and YAP1-associated plasticity are not sufficient to confer regenerative competence. Weakening mature-state constraints may permit greater cellular plasticity, but plasticity alone is insufficient for regeneration and can instead compromise cardiac function (*17–20*). These findings have implications beyond RNF20 deficiency, extending to both cardiac regeneration and the generation of stable mature cardiomyocytes in vitro. Successful regeneration requires coordinated control of transient state destabilization, productive cell-cycle progression, and subsequent rematuration (*41, 89*). In a related context, durable maturation of induced pluripotent stem cell-derived cardiomyocytes may depend not only on induction of adult gene-expression programs but also on establishment of the chromatin maintenance systems that continuously reinforce them.

The spatial organization of the response further shows that intrinsic chromatin instability is interpreted through the tissue microenvironment. Group 3^High^ bins were preferentially localized to the expanded subendocardial niche in the cKO heart (cluster 8; Fig. 6K, 6L, and 6M). RNF20 loss therefore creates cell-intrinsic susceptibility to state transition, whereas regional physiological conditions determine where the stress-responsive program is most strongly activated. Compared with the subepicardium, the subendocardium is exposed to distinctive electrophysiological conditions (*90*), greater mechanical stress (*91*), higher metabolic demand (*92, 93*), and increased ischemic vulnerability (*94*), potentially reducing its resilience to disruption of the mature cardiomyocyte state. The accompanying Cx43 loss and fibrosis further indicate that cardiomyocyte maturity is not solely a cell-autonomous transcriptional property. Mature cardiomyocytes help organize electrical continuity, force transmission, and stromal homeostasis; when their state destabilizes, the surrounding ecosystem is remodeled into a pathological rather than pro-renewal niche. Together, our findings support a model in which the mature cardiomyocyte attractor state is maintained through a selective chromatin-based reinforcement. RNF20-dependent chromatin regulation continuously supports core functional transcriptional programs, while restraining excessive activation of stress-responsive regulatory elements under physiological conditions. Loss of this homeostasis instead permits spatially patterned transitions toward pathological cell states.

Several limitations of this study also highlight important directions for future investigation. Although RNF20 is the principal H2B ubiquitin ligase in mammalian cells, it also possesses H2Bub-independent and potentially non-histone functions (*95–97*). The relative contribution of these activities to mature cardiomyocyte state maintenance remains to be determined. H2Bub, FACT, and Pol II occupancy, nascent transcription, and transcript completion were not directly measured in adult RNF20-deficient cardiomyocytes, so the proposed relationships among gene-body H2Bub, promoter H3K4me3 breadth, and transcriptional fidelity remain mechanistic models. Although YAP1 localization, target expression, and motif accessibility implicate YAP1- and AP-1-associated regulation, their causal contributions have not been tested mechanistically. Accessibility losses at CTCF-associated regions (Fig. 5F and 5K) further raise the possibility of altered higher-order genome organization (*98, 99*), though CTCF occupancy and chromatin contacts were not examined. Finally, the signals responsible for the subendocardial niche, the reversibility of the altered states after RNF20 restoration, and the extent to which this maintenance mechanism is disrupted in human heart disease await further discovery.

## MATERIALS and METHODS

### Study design and animal models

All animal experiments were approved by the Institutional Animal Care and Use Committee of Academia Sinica (Protocol #24-06-2184). Mice were maintained under specific pathogen-free conditions with ad libitum access to food and water. Tamoxifen-inducible cardiomyocyte-specific *Rnf20* knockout mice (cKO) were generated by crossing *Rnf20^flox/flox^* mice (*33*) with αMHC-MerCreMer^+/−^ (CreERT2) mice (*35*) on a C57BL/6JNarl background. CreERT2 mice were used as controls. Male mice were used throughout the study.

At 8–10 weeks of age, mice received tamoxifen (20 mg/kg, intraperitoneally) once daily for 5 consecutive days to induce Cre-mediated recombination. Unless otherwise indicated, molecular and cellular analyses were performed at the time points specified in the corresponding figures. For cardiomyocyte proliferation assays, EdU (50 mg/kg) was administered intraperitoneally on 2 consecutive days, and hearts were collected 1 day after the final injection.

### Protein and RNA analyses

LV tissues were used for Western blotting and RT-qPCR. For Western blotting, proteins were extracted in RIPA buffer containing protease and phosphatase inhibitors, separated by SDS-PAGE, transferred to PVDF membranes, and detected by chemiluminescence. RNF20, Cx43, YAP1, phospho-YAP1(S127), α-tubulin, and detyrosinated α-tubulin were analyzed using the antibodies listed in the Supplementary Materials. Ponceau S total-protein staining or GAPDH was used for normalization.

Total RNA was isolated from LV tissue using TRIzol, treated to remove genomic DNA, and reverse-transcribed using the QuantiTect Reverse Transcription Kit. RT-qPCR was performed using KAPA SYBR FAST qPCR Master Mix. *Gapdh* or 18S rRNA served as internal controls, and relative expression was calculated by the 2^−ΔΔCp^ method. Primer sequences are provided in Table S1.

### Histology, cardiomyocyte isolation, and immunofluorescence

For fibrosis analysis, hearts were fixed in 4% paraformaldehyde, paraffin-embedded, sectioned at 5-μm, and subjected to Masson’s Trichrome staining. Fibrotic area was quantified as the collagen-positive area relative to myocardial area in the LV free wall and interventricular septum.

Adult cardiomyocytes were isolated using a Langendorff-free enzymatic perfusion method (*100*). Hearts were digested by perfusion with an enzyme mixture containing collagenase B, collagenase D, and protease XIV. Dissociated cardiomyocytes were separated from nonmyocytes by low-speed centrifugation, fixed in 4% paraformaldehyde, and cytospun onto glass slides.

For tissue immunofluorescence, perfusion-fixed hearts were cryoprotected, embedded in OCT, and sectioned at 10-μm. Tissue sections or isolated cardiomyocytes were permeabilized, blocked, and incubated with primary antibodies against proteins including cTnT, α-actinin, α-SMA, Cx43, N-cadherin, YAP1, dystrophin, Ki67, and PCM1, followed by fluorophore-conjugated secondary antibodies and DAPI. EdU incorporation was detected using the Click-iT Plus EdU Cell Proliferation Kit, and apoptotic nuclei were detected by TUNEL assay. Images were acquired using a Zeiss LSM880 confocal microscope.

Sarcomere organization in cTnT-stained tissue sections and α-actinin-stained isolated cardiomyocytes was quantified using SotaTool (*40, 43*). Cardiomyocyte nucleation, nuclear area, nuclear YAP1, α-SMA-positive area, EdU- and Ki67-positive cardiomyocytes, TUNEL-positive nuclei, and Cx43 localization at N-cadherin-positive intercalated discs were quantified as described in the Supplementary Materials.

### Echocardiography and electrocardiography

Cardiac function was evaluated longitudinally by transthoracic echocardiography and surface electrocardiography under 1–1.5% isoflurane anesthesia. Echocardiography was performed using a Vevo 3100 ultrasound system. LV dimensions and wall thicknesses were measured, from which LV volumes, ejection fraction, and LV mass were calculated.

Surface electrocardiograms were recorded using a PowerLab 8/30 system with an Animal Bio Amp. Standard limb-lead signals were recorded for 10–15 min and analyzed using LabChart and Cardiac Axis software. QRS intervals were determined from the resulting ECG traces.

### Single-nucleus RNA sequencing and analysis

LV nuclei were isolated by mechanical homogenization followed by filtration and fluorescence-activated sorting of 7-AAD-positive singlet nuclei. Single-nucleus RNA-seq libraries (n = 1 per genotype) were generated using the Chromium Next GEM Single Cell 3′ v3.1 platform (10x Genomics), targeting approximately 10,000 nuclei per sample, and sequenced on an Illumina NextSeq 2000.

Reads were aligned to the mm10 genome using STARSolo (*101*). Low-quality droplets and doublets were removed using the Trailmaker pipeline (https://app.trailmaker.parsebiosciences.com/). Subsequent analyses were performed using Seurat (*102*). Expression values were log-normalized, highly variable genes were identified, and dimensionality reduction and clustering were performed by principal component analysis, Harmony integration, Louvain clustering, and UMAP. Cell types were assigned according to established marker genes.

Cardiomyocytes and fibroblasts were independently subclustered. Relative cardiomyocyte differentiation state was assessed using CytoTRACE (*36*), and transcriptional gene-set activity was quantified using AUCell (*38*). Cardiomyocyte state transition was further examined using Monocle3 pseudotime analysis (*48*).

To compare RNF20-deficient cardiomyocytes with developmental cardiomyocyte states, published snRNA-seq dataset from neonatal murine hearts following myocardial infarction (*58*) was reanalyzed and cell-state identities were transferred to the present dataset using Seurat anchor-based mapping.

Overrepresentation analysis of selected gene sets was assessed using DAVID (*103*), with an FDR < 0.05 considered significant.

### Cardiomyocyte ATAC-seq and analysis

Cardiomyocyte nuclei were purified from LV tissue by fluorescence-activated sorting of PCM1- and 7-AAD-positive nuclei. ATAC-seq libraries (n = 3 per genotype) were generated using TDE1 transposase and sequenced as 150-bp paired-end reads on an Illumina NovaSeq X Plus.

Preprocessing was carried out on Galaxy (*104*). Following adapter and quality trimming, reads were aligned to the mm10 genome using Bowtie2 (*105*). Low-quality, mitochondrial, duplicate, and ENCODE-blacklisted reads (*106*) were removed, and Tn5 integration offsets were corrected. Peaks were called using MACS2 (*107*) and combined into a consensus peak set. Differential accessibility was analyzed using DESeq2 (*108*) and peaks with adjusted *p* < 0.05 were considered differentially accessible.

Peaks were annotated relative to genomic features using ChIPseeker (*109*), with promoters defined as ±2 kb from transcription start sites. Chromatin-state annotations were generated with ChromHMM (*61*) using adult mouse-heart chromatin immunoprecipitation sequencing (ChIP-seq) data for H3K4me3, H3K27ac, H3K4me1, H3K27me3, H3K36me3, and CTCF obtained from ENCODE. Functional enrichment of differential accessible regions was analyzed using rGREAT (*65*). Transcription factor motif enrichment was evaluated using HOMER (*110*), and differential transcription-factor footprinting was performed using TOBIAS (*67*).

Published ATAC-seq data from cardiomyocytes expressing constitutively active YAP1 (YAP5SA) (*51*) were processed using the same pipeline for comparative analyses.

### Analysis of published transcriptomic and ChIP-seq datasets

Published murine cardiomyocyte RNA-seq dataset spanning embryonic day 14.5 through postnatal day 56 (*111*) were reanalyzed to define maturation-associated transcriptional programs. Genes exhibiting significant developmental expression dynamics were identified using maSigPro (*112*) and divided into genes increasing during maturation (mature gene set) and genes decreasing during maturation (immature gene set).

Published H2Bub ChIP-seq from P28 murine hearts (*34*), H3K4me3 and H3K27ac ChIP-seq from adult murine cardiomyocyte nuclei (*12*), cardiomyocyte Pol II ChIP-seq (*78*), and RNA-seq and H3K4me3 ChIP-seq from human failing and non-failing cardiomyocyte nuclei (*80*) were reanalyzed using standardized pipelines. Reads were aligned to mm10 or hg38 as appropriate. H3K4me3 domains were identified using broad MACS2 (*107*) peak calling and assigned to associated genes.

Pol II promoter-proximal pausing was quantified using the Pausing Index Calculator (*113*). The pausing index was calculated as the normalized Pol II density within the promoter-proximal region (−50 to +300 bp relative to the transcription start site) divided by Pol II density across the gene body.

### Tracks visualization

Normalized sequencing tracks and genome-wide profiles were generated using deepTools (*114*), and locus-level tracks were visualized using Integrative Genomics Viewer (*115*).

### Spatial transcriptomics and analysis

Spatial transcriptomic profiling was performed using the Visium HD Spatial Gene Expression platform (10x Genomics; n = 1 per genotype). Formalin-fixed paraffin-embedded heart sections were sectioned at 5-μm, stained with hematoxylin and eosin, imaged, and processed following the Visium HD Spatial Gene Expression Reagent Kits User Guide (CG000685). Libraries were sequenced on an Illumina NextSeq 2000.

Reads were processed with Space Ranger against the GRCm39 genome, and downstream analyses were performed using 8-μm bins in Seurat (*102*). Nonventricular and low-quality bins were excluded. Data were normalized using SCTransform, integrated using Harmony, and clustered by the Louvain algorithm. Clusters displaying technical artifacts, very low abundance, or erythrocyte identity were removed before downstream analysis.

Cell-type composition of spatial bins was inferred using Robust Cell Type Decomposition (RCTD) (*116*) with the snRNA-seq dataset generated in this study as the reference. Cardiomyocyte-enriched bins were defined as bins with an RCTD cardiomyocyte weight >0.8. AUCell (*38*) was used to map the snRNA-seq-derived pseudotime gene programs onto cardiomyocyte-enriched spatial bins. Bins in cKO hearts with Group 2 or Group 3 AUCell scores exceeding the 99th percentile of the corresponding CreERT2 distribution were defined as Group 2^High^ or Group 3^High^.

For distance-based analysis of the Group 3 program, spatial cluster 8 bins surrounded by at least 25 neighboring cluster 8 bins were defined as the core, and surrounding bins were stratified according to distances of ≤50, ≤100, ≤200, or >200 μm from this region. The frequency of Group 3^High^ cardiomyocyte-enriched bins was quantified across these zones.

### Statistical analysis

Statistical analyses were performed using R and GraphPad Prism. Two-group comparisons were conducted using unpaired Student’s *t* tests, with Welch’s correction when appropriate, or Mann–Whitney tests for nonparametric data. Comparisons involving more than two groups used one-way ANOVA followed by Tukey’s test or Kruskal–Wallis followed by Dunn’s test. Longitudinal genotype × time effects on cardiac measurements were evaluated using mixed-effects models. Survival was compared by log-rank test, categorical variables by chi-square test, and spatial enrichment of Group 3^High^ bins across distance-resolved zones by a one-sided Cochran–Armitage trend test. Cell-type enrichment among spatial clusters was assessed by one-sided Wilcoxon rank-sum tests with Bonferroni correction. A two-sided *p* < 0.05 was considered statistically significant unless otherwise specified. Sample sizes and error-bar definitions are provided in the corresponding figure legends. No statistical method was used to predetermine sample size. Experiments were not randomized, and investigators were not blinded to group allocation or outcome assessment.

Detailed experimental procedures, reagent information, antibody specifications, computational parameters, and analysis workflows are provided in the Supplementary Materials.

## Acknowledgements

The authors thank Wei-Chen Chu (Imaging Core Facility) and Hsin-Yi Du (Biochemistry Core Facility), Institute of Cellular and Organismic Biology, Academia Sinica, for technical support with immunofluorescence imaging; the Laboratory Animal Core Facility, Institute of Cellular and Organismic Biology, Academia Sinica, for mouse husbandry; the Transgenic Core Facility, Institute of Molecular Biology, Academia Sinica, for technical support in generating transgenic mice; Yi-Hua Chen, High-Throughput Genomics Core, Biodiversity Research Center, Academia Sinica, for technical support with snRNA-seq and spatial transcriptomics; I-Chi Yang, Taiwan Mouse Clinic, Academia Sinica, for technical support with echocardiography and electrocardiography; and the Pathology Core Facility, Institute of Biomedical Sciences, Academia Sinica, for Masson’s Trichrome staining. Schematics were created with BioRender under a publication license.

## Funding

This work was supported by the National Science and Technology Council, Taiwan (NSTC 114-2320-B-001-019-MY3 to C.-F.K.) and Academia Sinica, Taiwan (AS-IV-114-L07 to C.-F.K.).

## Author contributions

H.-H.T. performed most of the experiments and data analyses and contributed to manuscript preparation. H.-Y.T. assisted with the single-nucleus RNA-seq and ATAC-seq experiments. C.-Y.L. assisted with experiments and data analysis. C.-F.K. conceived and supervised the study and contributed to manuscript preparation. H.-H.T. and C.-F.K. wrote the manuscript.

## Declaration of interests

The authors declare that they have no competing interests.

## Declaration of generative AI and AI-assisted technologies

During preparation of this manuscript, the authors used ChatGPT (OpenAI) to assist with bioinformatics code troubleshooting and language editing. All AI-assisted outputs were reviewed, verified, and edited by the authors, who take full responsibility for the final content.

## Data and code availability

All data needed to evaluate the conclusions in the paper are present in the paper and/or the Supplementary Materials.

The raw sequencing data generated in this study have been deposited in the National Center for Biotechnology Information Sequence Read Archive under BioProject accession PRJNA1496066. The data will be released upon publication.

The uncropped blots (https://doi.org/10.5281/zenodo.21700467) and codes generated in this study (https://doi.org/10.5281/zenodo.21700369) have been deposited in Zenodo.

## Notes

### Competing Interest Statement

The authors have declared no competing interest.

## REFERENCES

1. W. A. Whyte, D. A. Orlando, D. Hnisz, B. J. Abraham, C. Y. Lin, M. H. Kagey, P. B. Rahl, T. I. Lee, R. A. Young, Master transcription factors and mediator establish super-enhancers at key cell identity genes. Cell 153, 307–319 (2013).

2. B. K. Lee, A. A. Bhinge, A. Battenhouse, R. M. McDaniell, Z. Liu, L. Song, Y. Ni, E. Birney, J. D. Lieb, T. S. Furey, G. E. Crawford, V. R. Iyer, Cell-type specific and combinatorial usage of diverse transcription factors revealed by genome-wide binding studies in multiple human cells. Genome Res 22, 9–24 (2012).

3. J. Ernst, P. Kheradpour, T. S. Mikkelsen, N. Shoresh, L. D. Ward, C. B. Epstein, X. Zhang, L. Wang, R. Issner, M. Coyne, M. Ku, T. Durham, M. Kellis, B. E. Bernstein, Mapping and analysis of chromatin state dynamics in nine human cell types. Nature 473, 43–49 (2011).

4. Y. Wang, J. Li, A. A. Malcolm, W. Mansfield, S. J. Clark, R. Argelaguet, L. Biggins, R. J. Acton, S. Andrews, W. Reik, G. Kelsey, P. J. Rugg-Gunn, Combinatorial profiling of multiple histone modifications and transcriptome in single cells using scMTR-seq. Sci Adv 11, eadu3308 (2025).

5. S. Liu, C. Y. Wang, P. Zheng, B. B. Jia, N. R. Zemke, P. Ren, H. L. Park, B. Ren, X. Zhuang, Cell type-specific 3D-genome organization and transcription regulation in the brain. Sci Adv 11, eadv2067 (2025).

6. J. Holmberg, T. Perlmann, Maintaining differentiated cellular identity. Nat Rev Genet 13, 429–439 (2012).

7. C. Jopling, S. Boue, J. C. Izpisua Belmonte, Dedifferentiation, transdifferentiation and reprogramming: three routes to regeneration. Nat Rev Mol Cell Biol 12, 79–89 (2011).

8. S. Huang, G. Eichler, Y. Bar-Yam, D. E. Ingber, A. New Collective, Cell fates as high-dimensional attractor states of a complex gene regulatory network. Phys Rev Lett 94, 128701 (2005).

9. T. Enver, M. Pera, C. Peterson, P. W. Andrews, Stem cell states, fates, and the rules of attraction. Cell Stem Cell 4, 387–397 (2009).

10. W. A. Flavahan, E. Gaskell, B. E. Bernstein, Epigenetic plasticity and the hallmarks of cancer. Science 357, (2017).

11. J. R. Alvarez-Dominguez, D. A. Melton, Cell maturation: Hallmarks, triggers, and manipulation. Cell 185, 235–249 (2022).

12. R. Gilsbach, S. Preissl, B. A. Gruning, T. Schnick, L. Burger, V. Benes, A. Wurch, U. Bonisch, S. Gunther, R. Backofen, B. K. Fleischmann, D. Schubeler, L. Hein, Dynamic DNA methylation orchestrates cardiomyocyte development, maturation and disease. Nat Commun 5, 5288 (2014).

13. Y. Guo, W. T. Pu, Cardiomyocyte Maturation: New Phase in Development. Circ Res 126, 1086–1106 (2020).

14. E. R. Porrello, A. I. Mahmoud, E. Simpson, J. A. Hill, J. A. Richardson, E. N. Olson, H. A. Sadek, Transient regenerative potential of the neonatal mouse heart. Science 331, 1078–1080 (2011).

15. Y. Zhu, V. D. Do, A. M. Richards, R. Foo, What we know about cardiomyocyte dedifferentiation. J Mol Cell Cardiol 152, 80–91 (2021).

16. T. Kubin, J. Poling, S. Kostin, P. Gajawada, S. Hein, W. Rees, A. Wietelmann, M. Tanaka, H. Lorchner, S. Schimanski, M. Szibor, H. Warnecke, T. Braun, Oncostatin M is a major mediator of cardiomyocyte dedifferentiation and remodeling. Cell Stem Cell 9, 420–432 (2011).

17. E. Bassat, J. Wang, J. Peña Peña, I. Rivero-García, A. Piszczek, F. Falcon, Y. Taniguchi-Sugiura, J. Yang, P. Fernández-Montes, T. Lendl, L. Domínguez, K. Lust, J. A. Enríquez, F. S. Cabo, M. Torres, E. M. Tanaka, AXL governs axolotl cardiac regeneration and directs mammalian cardiomyocyte dedifferentiation. bioRxiv, 2025.2012.2021.695613 (2025).

18. L. R. J. Bailey, D. Bugg, I. M. Reichardt, C. D. Ortac, A. Nagle, J. Gunaje, A. Martinson, R. Johnson, M. J. MacCoss, T. Sakamoto, D. P. Kelly, M. Regnier, J. Davis, MBNL1 Regulates Programmed Postnatal Switching Between Regenerative and Differentiated Cardiac States. Circulation 149, 1812–1829 (2024).

19. M. R. Pricolo, M. A. Lopez-Unzu, N. Vicente, C. Morales-Lopez, C. Huerta-Lopez, W. Perez-Franco, A. C. Dumitru, J. Pena-Pena, F. M. Espinosa, M. I. Sanchez, R. Garcia, R. Silva-Rojas, M. Torres, E. Herrero-Galan, J. Alegre-Cebollada, Titin cleavage in living cardiomyocytes induces sarcomere disassembly but does not trigger cell proliferation. J Biol Chem 302, 113167 (2026).

20. T. Koopmans, E. van Rooij, Molecular gatekeepers of endogenous adult mammalian cardiomyocyte proliferation. Nat Rev Cardiol 22, 857–882 (2025).

21. Y. Li, S. Ai, X. Yu, C. Li, X. Li, Y. Yue, Y. Wei, C. Y. Li, A. He, Replication-Independent Histone Turnover Underlines the Epigenetic Homeostasis in Adult Heart. Circ Res 125, 198–208 (2019).

22. R. Papait, S. Serio, C. Pagiatakis, F. Rusconi, P. Carullo, M. Mazzola, N. Salvarani, M. Miragoli, G. Condorelli, Histone Methyltransferase G9a Is Required for Cardiomyocyte Homeostasis and Hypertrophy. Circulation 136, 1233–1246 (2017).

23. A. B. Stein, T. A. Jones, T. J. Herron, S. R. Patel, S. M. Day, S. F. Noujaim, M. L. Milstein, M. Klos, P. B. Furspan, J. Jalife, G. R. Dressler, Loss of H3K4 methylation destabilizes gene expression patterns and physiological functions in adult murine cardiomyocytes. J Clin Invest 121, 2641–2650 (2011).

24. S. Venkatesh, J. L. Workman, Histone exchange, chromatin structure and the regulation of transcription. Nat Rev Mol Cell Biol 16, 178–189 (2015).

25. B. Zhu, Y. Zheng, A. D. Pham, S. S. Mandal, H. Erdjument-Bromage, P. Tempst, D. Reinberg, Monoubiquitination of human histone H2B: the factors involved and their roles in HOX gene regulation. Mol Cell 20, 601–611 (2005).

26. N. Minsky, E. Shema, Y. Field, M. Schuster, E. Segal, M. Oren, Monoubiquitinated H2B is associated with the transcribed region of highly expressed genes in human cells. Nat Cell Biol 10, 483–488 (2008).

27. L. Wu, L. Li, B. Zhou, Z. Qin, Y. Dou, H2B ubiquitylation promotes RNA Pol II processivity via PAF1 and pTEFb. Mol Cell 54, 920–931 (2014).

28. R. Pavri, B. Zhu, G. Li, P. Trojer, S. Mandal, A. Shilatifard, D. Reinberg, Histone H2B monoubiquitination functions cooperatively with FACT to regulate elongation by RNA polymerase II. Cell 125, 703–717 (2006).

29. A. Luo, J. Kong, J. Chen, X. Xiao, J. Lan, X. Li, C. Liu, P. Y. Wang, G. Li, W. Li, P. Chen, H2B ubiquitination recruits FACT to maintain a stable altered nucleosome state for transcriptional activation. Nat Commun 14, 741 (2023).

30. G. Fuchs, E. Shema, R. Vesterman, E. Kotler, Z. Wolchinsky, S. Wilder, L. Golomb, A. Pribluda, F. Zhang, M. Haj-Yahya, E. Feldmesser, A. Brik, X. Yu, J. Hanna, D. Aberdam, E. Domany, M. Oren, RNF20 and USP44 regulate stem cell differentiation by modulating H2B monoubiquitylation. Mol Cell 46, 662–673 (2012).

31. O. Karpiuk, Z. Najafova, F. Kramer, M. Hennion, C. Galonska, A. Konig, N. Snaidero, T. Vogel, A. Shchebet, Y. Begus-Nahrmann, M. Kassem, M. Simons, H. Shcherbata, T. Beissbarth, S. A. Johnsen, The histone H2B monoubiquitination regulatory pathway is required for differentiation of multipotent stem cells. Mol Cell 46, 705–713 (2012).

32. L. Wang, Z. Xu, L. Wang, C. Liu, H. Wei, R. Zhang, Y. Chen, L. Wang, W. Liu, S. Xiao, W. Li, W. Li, Histone H2B ubiquitination mediated chromatin relaxation is essential for the induction of somatic cell reprogramming. Cell Prolif 54, e13080 (2021).

33. C.-Y. Lin, Y.-M. Chang, H.-Y. Tseng, Y.-L. Shih, H.-H. Yeh, Y.-R. Liao, H.-H. Tang, C.-L. Hsu, C.-C. Chen, Y.-T. Yan, C.-F. Kao, Epigenetic regulator RNF20 underlies temporal hierarchy of gene expression to regulate postnatal cardiomyocyte polarization. Cell Reports 42, 113416 (2023).

34. N. J. VanDusen, J. Y. Lee, W. Gu, C. E. Butler, I. Sethi, Y. Zheng, J. S. King, P. Zhou, S. Suo, Y. Guo, Q. Ma, G. C. Yuan, W. T. Pu, Massively parallel in vivo CRISPR screening identifies RNF20/40 as epigenetic regulators of cardiomyocyte maturation. Nat Commun 12, 4442 (2021).

35. D. S. Sohal, M. Nghiem, M. A. Crackower, S. A. Witt, T. R. Kimball, K. M. Tymitz, J. M. Penninger, J. D. Molkentin, Temporally regulated and tissue-specific gene manipulations in the adult and embryonic heart using a tamoxifen-inducible Cre protein. Circ Res 89, 20–25 (2001).

36. G. S. Gulati, S. S. Sikandar, D. J. Wesche, A. Manjunath, A. Bharadwaj, M. J. Berger, F. Ilagan, A. H. Kuo, R. W. Hsieh, S. Cai, M. Zabala, F. A. Scheeren, N. A. Lobo, D. Qian, F. B. Yu, F. M. Dirbas, M. F. Clarke, A. M. Newman, Single-cell transcriptional diversity is a hallmark of developmental potential. Science 367, 405–411 (2020).

37. S. Aibar, C. B. Gonzalez-Blas, T. Moerman, V. A. Huynh-Thu, H. Imrichova, G. Hulselmans, F. Rambow, J. C. Marine, P. Geurts, J. Aerts, J. van den Oord, Z. K. Atak, J. Wouters, S. Aerts, SCENIC: single-cell regulatory network inference and clustering. Nat Methods 14, 1083–1086 (2017).

38. S. Aibar, S. Aerts, AUCell: analysis of’gene set’activity in single-cell RNA-seq data. R/Bioconductor package, (2016).

39. F. B. Bedada, S. S. Chan, S. K. Metzger, L. Zhang, J. Zhang, D. J. Garry, T. J. Kamp, M. Kyba, J. M. Metzger, Acquisition of a quantitative, stoichiometrically conserved ratiometric marker of maturation status in stem cell-derived cardiac myocytes. Stem Cell Reports 3, 594–605 (2014).

40. Y. Zhu, M. Ackers-Johnson, M. K. Shanmugam, L. S. Pakkiri, C. L. Drum, Y. Chen, J. Kim, W. G. Paltzer, A. I. Mahmoud, W. L. W. Tan, M. C. J. Lee, J. Jiang, D. A. T. Luu, S. L. Ng, P. Y. Q. Li, A. Wang, R. Qi, G. J. X. Ong, T. Y. Ng, J. J. Haigh, Z. Tiang, A. M. Richards, R. S. Y. Foo, Asparagine Synthetase Marks a Distinct Dependency Threshold for Cardiomyocyte Dedifferentiation. Circulation 149, 1833–1851 (2024).

41. G. D’Uva, A. Aharonov, M. Lauriola, D. Kain, Y. Yahalom-Ronen, S. Carvalho, K. Weisinger, E. Bassat, D. Rajchman, O. Yifa, M. Lysenko, T. Konfino, J. Hegesh, O. Brenner, M. Neeman, Y. Yarden, J. Leor, R. Sarig, R. P. Harvey, E. Tzahor, ERBB2 triggers mammalian heart regeneration by promoting cardiomyocyte dedifferentiation and proliferation. Nat Cell Biol 17, 627–638 (2015).

42. A. Beisaw, C. Kuenne, S. Guenther, J. Dallmann, C. C. Wu, M. Bentsen, M. Looso, D. Y. R. Stainier, AP-1 Contributes to Chromatin Accessibility to Promote Sarcomere Disassembly and Cardiomyocyte Protrusion During Zebrafish Heart Regeneration. Circ Res 126, 1760–1778 (2020).

43. J. M. Stein, U. Arslan, M. Franken, J. C. de Greef, E. H. S, N. Mohammadi, V. V. Orlova, M. Bellin, C. L. Mummery, B. J. van Meer, Software Tool for Automatic Quantification of Sarcomere Length and Organization in Fixed and Live 2D and 3D Muscle Cell Cultures In Vitro. Curr Protoc 2, e462 (2022).

44. D. L. Ruzicka, R. J. Schwartz, Sequential activation of alpha-actin genes during avian cardiogenesis: vascular smooth muscle alpha-actin gene transcripts mark the onset of cardiomyocyte differentiation. J Cell Biol 107, 2575–2586 (1988).

45. Y. Chen, F. F. Luttmann, E. Schoger, H. R. Scholer, L. C. Zelarayan, K. P. Kim, J. J. Haigh, J. Kim, T. Braun, Reversible reprogramming of cardiomyocytes to a fetal state drives heart regeneration in mice. Science 373, 1537–1540 (2021).

46. P. Michela, V. Velia, P. Aldo, P. Ada, Role of connexin 43 in cardiovascular diseases. Eur J Pharmacol 768, 71–76 (2015).

47. A. Aharonov, A. Shakked, K. B. Umansky, A. Savidor, A. Genzelinakh, D. Kain, D. Lendengolts, O. Y. Revach, Y. Morikawa, J. Dong, Y. Levin, B. Geiger, J. F. Martin, E. Tzahor, ERBB2 drives YAP activation and EMT-like processes during cardiac regeneration. Nat Cell Biol 22, 1346–1356 (2020).

48. J. Cao, M. Spielmann, X. Qiu, X. Huang, D. M. Ibrahim, A. J. Hill, F. Zhang, S. Mundlos, L. Christiansen, F. J. Steemers, C. Trapnell, J. Shendure, The single-cell transcriptional landscape of mammalian organogenesis. Nature 566, 496–502 (2019).

49. S. Dupont, L. Morsut, M. Aragona, E. Enzo, S. Giulitti, M. Cordenonsi, F. Zanconato, J. Le Digabel, M. Forcato, S. Bicciato, N. Elvassore, S. Piccolo, Role of YAP/TAZ in mechanotransduction. Nature 474, 179–183 (2011).

50. M. Xin, Y. Kim, L. B. Sutherland, M. Murakami, X. Qi, J. McAnally, E. R. Porrello, A. I. Mahmoud, W. Tan, J. M. Shelton, J. A. Richardson, H. A. Sadek, R. Bassel-Duby, E. N. Olson, Hippo pathway effector Yap promotes cardiac regeneration. Proc Natl Acad Sci U S A 110, 13839–13844 (2013).

51. T. O. Monroe, M. C. Hill, Y. Morikawa, J. P. Leach, T. Heallen, S. Cao, P. H. L. Krijger, W. de Laat, X. H. T. Wehrens, G. G. Rodney, J. F. Martin, YAP Partially Reprograms Chromatin Accessibility to Directly Induce Adult Cardiogenesis In Vivo. Dev Cell 48, 765–779 e767 (2019).

52. Y. Morikawa, J. H. Kim, R. G. Li, L. Liu, S. Liu, V. Deshmukh, M. C. Hill, J. F. Martin, YAP Overcomes Mechanical Barriers to Induce Mitotic Rounding and Adult Cardiomyocyte Division. Circulation 151, 76–93 (2025).

53. Y. Morikawa, M. Zhang, T. Heallen, J. Leach, G. Tao, Y. Xiao, Y. Bai, W. Li, J. T. Willerson, J. F. Martin, Actin cytoskeletal remodeling with protrusion formation is essential for heart regeneration in Hippo-deficient mice. Sci Signal 8, ra41 (2015).

54. J. Dong, G. Feldmann, J. Huang, S. Wu, N. Zhang, S. A. Comerford, M. F. Gayyed, R. A. Anders, A. Maitra, D. Pan, Elucidation of a universal size-control mechanism in Drosophila and mammals. Cell 130, 1120–1133 (2007).

55. A. Elosegui-Artola, I. Andreu, A. E. M. Beedle, A. Lezamiz, M. Uroz, A. J. Kosmalska, R. Oria, J. Z. Kechagia, P. Rico-Lastres, A. L. Le Roux, C. M. Shanahan, X. Trepat, D. Navajas, S. Garcia-Manyes, P. Roca-Cusachs, Force Triggers YAP Nuclear Entry by Regulating Transport across Nuclear Pores. Cell 171, 1397–1410 e1314 (2017).

56. I. Duursma, E. E. Nollet, V. Jansen, J. A. Malone, J. S. Bloem, K. Bedi, K. B. Margulies, S. A. C. Schoonvelde, M. Michels, N. N. van der Wel, J. van der Velden, T. J. Kirby, D. W. D. Kuster, Microtubule detyrosination alters nuclear mechanotransduction and leads to pro-hypertrophic signaling in hypertrophic cardiomyopathy. bioRxiv, (2025).

57. A. Gandin, V. Torresan, L. Ulliana, T. Panciera, P. Contessotto, A. Citron, F. Zanconato, M. Cordenonsi, S. Piccolo, G. Brusatin, Broadly Applicable Hydrogel Fabrication Procedures Guided by YAP/TAZ-Activity Reveal Stiffness, Adhesiveness, and Nuclear Projected Area as Checkpoints for Mechanosensing. Adv Healthc Mater 11, e2102276 (2022).

58. M. Cui, Z. Wang, K. Chen, A. M. Shah, W. Tan, L. Duan, E. Sanchez-Ortiz, H. Li, L. Xu, N. Liu, R. Bassel-Duby, E. N. Olson, Dynamic Transcriptional Responses to Injury of Regenerative and Non-regenerative Cardiomyocytes Revealed by Single-Nucleus RNA Sequencing. Dev Cell 55, 665–667 (2020).

59. M. Patterson, L. Barske, B. Van Handel, C. D. Rau, P. Gan, A. Sharma, S. Parikh, M. Denholtz, Y. Huang, Y. Yamaguchi, H. Shen, H. Allayee, J. G. Crump, T. I. Force, C. L. Lien, T. Makita, A. J. Lusis, S. R. Kumar, H. M. Sucov, Frequency of mononuclear diploid cardiomyocytes underlies natural variation in heart regeneration. Nat Genet 49, 1346–1353 (2017).

60. D. C. Zebrowski, S. Vergarajauregui, C. C. Wu, T. Piatkowski, R. Becker, M. Leone, S. Hirth, F. Ricciardi, N. Falk, A. Giessl, S. Just, T. Braun, G. Weidinger, F. B. Engel, Developmental alterations in centrosome integrity contribute to the post-mitotic state of mammalian cardiomyocytes. Elife 4, (2015).

61. J. Ernst, M. Kellis, Chromatin-state discovery and genome annotation with ChromHMM. Nat Protoc 12, 2478–2492 (2017).

62. E. P. Consortium, An integrated encyclopedia of DNA elements in the human genome. Nature 489, 57–74 (2012).

63. B. C. Hitz, J.-W. Lee, O. Jolanki, M. S. Kagda, K. Graham, P. Sud, I. Gabdank, J. Seth Strattan, C. A. Sloan, T. Dreszer, L. D. Rowe, N. R. Podduturi, V. S. Malladi, E. T. Chan, J. M. Davidson, M. Ho, S. Miyasato, M. Simison, F. Tanaka, Y. Luo, I. Whaling, E. L. Hong, B. T. Lee, R. Sandstrom, E. Rynes, J. Nelson, A. Nishida, A. Ingersoll, M. Buckley, M. Frerker, D. S. Kim, N. Boley, D. Trout, A. Dobin, S. Rahmanian, D. Wyman, G. Balderrama-Gutierrez, F. Reese, N. C. Durand, O. Dudchenko, D. Weisz, S. S. P. Rao, A. Blackburn, D. Gkountaroulis, M. Sadr, M. Olshansky, Y. Eliaz, D. Nguyen, I. Bochkov, M. S. Shamim, R. Mahajan, E. Aiden, T. Gingeras, S. Heath, M. Hirst, W. James Kent, A. Kundaje, A. Mortazavi, B. Wold, J. M. Cherry, The ENCODE Uniform Analysis Pipelines. bioRxiv, 2023.2004.2004.535623 (2023).

64. Y. Luo, B. C. Hitz, I. Gabdank, J. A. Hilton, M. S. Kagda, B. Lam, Z. Myers, P. Sud, J. Jou, K. Lin, U. K. Baymuradov, K. Graham, C. Litton, S. R. Miyasato, J. S. Strattan, O. Jolanki, J. W. Lee, F. Y. Tanaka, P. Adenekan, E. O’Neill, J. M. Cherry, New developments on the Encyclopedia of DNA Elements (ENCODE) data portal. Nucleic Acids Res 48, D882–D889 (2020).

65. Z. Gu, D. Hubschmann, rGREAT: an R/bioconductor package for functional enrichment on genomic regions. Bioinformatics 39, (2023).

66. G. Yu, F. Li, Y. Qin, X. Bo, Y. Wu, S. Wang, GOSemSim: an R package for measuring semantic similarity among GO terms and gene products. Bioinformatics 26, 976–978 (2010).

67. A. M. Bentsen, P. Goymann, H. Schultheis, K. Klee, A. Petrova, R. Wiegandt, Fust, J. Preussner, C. Kuenne, T. Braun, J. Kim, M. Looso, ATAC-seq footprinting unravels kinetics of transcription factor binding during zygotic genome activation. Nat Commun 11, 4267 (2020).

68. K. van Duijvenboden, D. E. M. de Bakker, J. C. K. Man, R. Janssen, M. Gunthel, M. C. Hill, I. B. Hooijkaas, I. van der Made, P. H. van der Kraak, A. Vink, E. E. Creemers, J. F. Martin, P. Barnett, J. Bakkers, V. M. Christoffels, Conserved NPPB+ Border Zone Switches From MEF2- to AP-1-Driven Gene Program. Circulation 140, 864–879 (2019).

69. A. M. Malek Mohammadi, B. Kattih, A. Grund, N. Froese, M. Korf-Klingebiel, Gigina, U. Schrameck, C. Rudat, Q. Liang, A. Kispert, K. C. Wollert, J. Bauersachs, J. Heineke, The transcription factor GATA4 promotes myocardial regeneration in neonatal mice. EMBO Mol Med 9, 265–279 (2017).

70. M. Ogawa, F. S. Geng, D. T. Humphreys, E. Kristianto, D. Z. Sheng, S. P. Hui, Y. Zhang, K. Sugimoto, M. Nakayama, D. Zheng, D. Hesselson, M. P. Hodson, O. Bogdanovic, K. Kikuchi, Kruppel-like factor 1 is a core cardiomyogenic trigger in zebrafish. Science 372, 201–205 (2021).

71. D. M. Cable, E. Murray, L. L. S. Zou, A. Goeva, E. Z. Macosko, F. Chen, R. A. Irizarry, Robust decomposition of cell type mixtures in spatial transcriptomics. Nat Biotechnol 40, 517–+ (2022).

72. E. S. Deneris, O. Hobert, Maintenance of postmitotic neuronal cell identity. Nat Neurosci 17, 899–907 (2014).

73. D. A. Gallegos, U. Chan, L. F. Chen, A. E. West, Chromatin Regulation of Neuronal Maturation and Plasticity. Trends Neurosci 41, 311–324 (2018).

74. S. Barish, K. Berg, J. Drozd, I. Berglund-Brown, L. Khizir, L. K. Wasson, C. E. Seidman, J. G. Seidman, S. Chen, M. Brueckner, The H2Bub1-deposition complex is required for human and mouse cardiogenesis. Development 150, (2023).

75. O. H. Funk, Y. Qalieh, D. Z. Doyle, M. M. Lam, K. Y. Kwan, Postmitotic accumulation of histone variant H3.3 in new cortical neurons establishes neuronal chromatin, transcriptome, and identity. Proc Natl Acad Sci U S A 119, e2116956119 (2022).

76. C. Liu, T. Maejima, S. C. Wyler, G. Casadesus, S. Herlitze, E. S. Deneris, Pet-1 is required across different stages of life to regulate serotonergic function. Nat Neurosci 13, 1190–1198 (2010).

77. S. Ai, X. Yu, Y. Li, Y. Peng, C. Li, Y. Yue, G. Tao, C. Li, W. T. Pu, A. He, Divergent Requirements for EZH1 in Heart Development Versus Regeneration. Circ Res 121, 106–112 (2017).

78. Z. Li, F. Yao, P. Yu, D. Li, M. Zhang, L. Mao, X. Shen, Z. Ren, L. Wang, B. Zhou, Postnatal state transition of cardiomyocyte as a primary step in heart maturation. Protein Cell 13, 842–862 (2022).

79. B. A. Benayoun, E. A. Pollina, D. Ucar, S. Mahmoudi, K. Karra, E. D. Wong, K. Devarajan, A. C. Daugherty, A. B. Kundaje, E. Mancini, B. C. Hitz, R. Gupta, T. A. Rando, J. C. Baker, M. P. Snyder, J. M. Cherry, A. Brunet, H3K4me3 breadth is linked to cell identity and transcriptional consistency. Cell 158, 673–688 (2014).

80. R. Gilsbach, M. Schwaderer, S. Preissl, B. A. Gruning, D. Kranzhofer, P. Schneider, T. G. Nuhrenberg, S. Mulero-Navarro, D. Weichenhan, C. Braun, M. Dressen, A. R. Jacobs, H. Lahm, T. Doenst, R. Backofen, M. Krane, B. D. Gelb, L. Hein, Distinct epigenetic programs regulate cardiac myocyte development and disease in the human heart in vivo. Nat Commun 9, 391 (2018).

81. W. Xie, S. Nagarajan, S. J. Baumgart, R. L. Kosinsky, Z. Najafova, V. Kari, M. Hennion, D. Indenbirken, S. Bonn, A. Grundhoff, F. Wegwitz, A. Mansouri, S. A. Johnsen, RNF40 regulates gene expression in an epigenetic context-dependent manner. Genome Biol 18, 32 (2017).

82. E. Shema, J. Kim, R. G. Roeder, M. Oren, RNF20 inhibits TFIIS-facilitated transcriptional elongation to suppress pro-oncogenic gene expression. Mol Cell 42, 477–488 (2011).

83. Y. Dou, N. Tetik-Elsherbiny, R. Gao, Y. Ren, Y. W. Chen, M. Merbecks, A. Setya, O. Lityagina, Y. Wang, E. Chichelnitskiy, A. Abouissa, C. C. Wu, G. Barreto, M. Potente, T. Wieland, R. Ola, P. Grieshaber, T. Loukanov, M. Gorenflo, J. Heineke, J. Cordero, G. Dobreva, Endothelial RNF20 suppresses endothelial-to-mesenchymal transition and safeguards physiological angiocrine signaling to prevent congenital heart disease. Nat Commun 16, 9480 (2025).

84. N. Tetik-Elsherbiny, A. Elsherbiny, A. Setya, J. Gahn, Y. Tang, P. Gupta, Y. Dou, H. Serke, T. Wieland, A. Dubrac, J. Heineke, M. Potente, J. Cordero, R. Ola, G. Dobreva, RNF20-mediated transcriptional pausing and VEGFA splicing orchestrate vessel growth. Nat Cardiovasc Res 3, 1199–1216 (2024).

85. C. J. Boogerd, I. Perini, E. Kyriakopoulou, S. J. Han, P. La, B. van der Swaan, J. B. Berkhout, D. Versteeg, J. Monshouwer-Kloots, E. van Rooij, Cardiomyocyte proliferation is suppressed by ARID1A-mediated YAP inhibition during cardiac maturation. Nat Commun 14, 4716 (2023).

86. L. Chang, L. Azzolin, D. Di Biagio, F. Zanconato, G. Battilana, R. Lucon Xiccato, M. Aragona, S. Giulitti, T. Panciera, A. Gandin, G. Sigismondo, J. Krijgsveld, M. Fassan, G. Brusatin, M. Cordenonsi, S. Piccolo, The SWI/SNF complex is a mechanoregulated inhibitor of YAP and TAZ. Nature 563, 265–269 (2018).

87. L. He, H. Pratt, M. Gao, F. Wei, Z. Weng, K. Struhl, YAP and TAZ are transcriptional co-activators of AP-1 proteins and STAT3 during breast cellular transformation. Elife 10, (2021).

88. W. Wang, C. K. Hu, A. Zeng, D. Alegre, D. Hu, K. Gotting, A. Ortega Granillo, Y. Wang, S. Robb, R. Schnittker, S. Zhang, D. Alegre, H. Li, E. Ross, N. Zhang, A. Brunet, A. Sanchez Alvarado, Changes in regeneration-responsive enhancers shape regenerative capacities in vertebrates. Science 369, (2020).

89. W. E. Wang, L. Li, X. Xia, W. Fu, Q. Liao, C. Lan, D. Yang, H. Chen, R. Yue, C. Zeng, L. Zhou, B. Zhou, D. D. Duan, X. Chen, S. R. Houser, C. Zeng, Dedifferentiation, Proliferation, and Redifferentiation of Adult Mammalian Cardiomyocytes After Ischemic Injury. Circulation 136, 834–848 (2017).

90. Q. Lou, V. V. Fedorov, A. V. Glukhov, N. Moazami, V. G. Fast, I. R. Efimov, Transmural heterogeneity and remodeling of ventricular excitation-contraction coupling in human heart failure. Circulation 123, 1881–1890 (2011).

91. G. Ciucci, D. Lorizio, N. Bartoloni, M. Budini, A. Colliva, S. Vodret, A. V. Nguyen, L. Ciacci, B. Texler, B. Cardini, R. Oberhuber, S. Bindelli, I. L. C. Del Giudice, R. Vuerich, F. Riccitelli, E. Zago, H. N. Finsberg, M. Chiesa, G. L. Perrucci, R. Bussani, F. Silvestri, M. Maglione, G. I. Dellino, G. Sinagra, M. Giacca, T. Eschenhagen, P. Golino, G. Pompilio, P. G. Pelicci, L. Andolfi, M. Pinamonti, M. Dal Ferro, S. Wall, F. S. Loffredo, S. Zacchigna, Mechanical load inhibits cancer growth in mouse and human hearts. Science 392, eads9412 (2026).

92. J. Vinten-Johansen, H. R. Weiss, Oxygen consumption in subepicardial and subendocardial regions of the canine left ventricle. The effect of experimental acute valvular aortic stenosis. Circ Res 46, 139–145 (1980).

93. D. J. Duncker, A. Koller, D. Merkus, J. M. Canty, Jr., Regulation of coronary blood flow in health and ischemic heart disease. Prog Cardiovasc Dis 57, 409–422 (2015).

94. D. Algranati, G. S. Kassab, Y. Lanir, Why is the subendocardium more vulnerable to ischemia? A new paradigm. Am J Physiol Heart Circ Physiol 300, H1090–1100 (2011).

95. J. H. Lee, G. Y. Lee, H. Jang, S. S. Choe, S. H. Koo, J. B. Kim, Ring finger protein20 regulates hepatic lipid metabolism through protein kinase A-dependent sterol regulatory element binding protein1c degradation. Hepatology 60, 844–857 (2014).

96. S. In, Y. I. Kim, J. E. Lee, J. Kim, RNF20/40-mediated eEF1BdeltaL monoubiquitylation stimulates transcription of heat shock-responsive genes. Nucleic Acids Res 47, 2840–2855 (2019).

97. Y. G. Jeon, J. H. Lee, Y. Ji, J. H. Sohn, D. Lee, D. W. Kim, S. G. Yoon, K. C. Shin, J. Park, J. K. Seong, J. Y. Cho, S. S. Choe, J. B. Kim, RNF20 Functions as a Transcriptional Coactivator for PPARgamma by Promoting NCoR1 Degradation in Adipocytes. Diabetes 69, 20–34 (2020).

98. J. M. Dowen, Z. P. Fan, D. Hnisz, G. Ren, B. J. Abraham, L. N. Zhang, A. S. Weintraub, J. Schujiers, T. I. Lee, K. Zhao, R. A. Young, Control of cell identity genes occurs in insulated neighborhoods in mammalian chromosomes. Cell 159, 374–387 (2014).

99. D. P. Lee, W. L. W. Tan, C. G. Anene-Nzelu, C. J. M. Lee, P. Y. Li, T. D. A. Luu, C. X. Chan, Z. Tiang, S. L. Ng, X. Huang, M. Efthymios, M. I. Autio, J. Jiang, M. J. Fullwood, S. Prabhakar, E. Lieberman Aiden, R. S. Foo, Robust CTCF-Based Chromatin Architecture Underpins Epigenetic Changes in the Heart Failure Stress-Gene Response. Circulation 139, 1937–1956 (2019).

100. M. Ackers-Johnson, P. Y. Li, A. P. Holmes, S. M. O’Brien, D. Pavlovic, R. S. Foo, A Simplified, Langendorff-Free Method for Concomitant Isolation of Viable Cardiac Myocytes and Nonmyocytes From the Adult Mouse Heart. Circ Res 119, 909–920 (2016).

101. B. Kaminow, D. Yunusov, A. Dobin, STARsolo: accurate, fast and versatile mapping/quantification of single-cell and single-nucleus RNA-seq data. bioRxiv, (2021).

102. A. Y. Hao, T. Stuart, M. H. Kowalski, S. Choudhary, P. Hoffman, A. Hartman, Srivastava, G. Molla, S. Madad, C. Fernandez-Granda, R. Satija, Dictionary learning for integrative, multimodal and scalable single-cell analysis. Nat Biotechnol 42, 293–304 (2024).

103. B. T. Sherman, M. Hao, J. Qiu, X. Jiao, M. W. Baseler, H. C. Lane, T. Imamichi, W. Chang, DAVID: a web server for functional enrichment analysis and functional annotation of gene lists (2021 update). Nucleic Acids Res 50, W216–W221 (2022).

104. C. Galaxy, Galaxy for accessible, reproducible, and collaborative data analyses: 2026 update. Nucleic Acids Res 54, W105–W116 (2026).

105. B. Langmead, S. L. Salzberg, Fast gapped-read alignment with Bowtie 2. Nat Methods 9, 357–359 (2012).

106. H. M. Amemiya, A. Kundaje, A. P. Boyle, The ENCODE Blacklist: Identification of Problematic Regions of the Genome. Sci Rep 9, 9354 (2019).

107. J. Feng, T. Liu, B. Qin, Y. Zhang, X. S. Liu, Identifying ChIP-seq enrichment using MACS. Nat Protoc 7, 1728–1740 (2012).

108. M. I. Love, W. Huber, S. Anders, Moderated estimation of fold change and dispersion for RNA-seq data with DESeq2. Genome Biol 15, 550 (2014).

109. G. Yu, L. G. Wang, Q. Y. He, ChIPseeker: an R/Bioconductor package for ChIP peak annotation, comparison and visualization. Bioinformatics 31, 2382–2383 (2015).

110. S. Heinz, C. Benner, N. Spann, E. Bertolino, Y. C. Lin, P. Laslo, J. X. Cheng, C. Murre, H. Singh, C. K. Glass, Simple combinations of lineage-determining transcription factors prime cis-regulatory elements required for macrophage and B cell identities. Mol Cell 38, 576–589 (2010).

111. J. Feng, Y. Li, Y. Nie, Methods of mouse cardiomyocyte isolation from postnatal heart. J Mol Cell Cardiol 168, 35–43 (2022).

112. A. Conesa, M. J. Nueda, A. Ferrer, M. Talon, maSigPro: a method to identify significantly differential expression profiles in time-course microarray experiments. Bioinformatics 22, 1096–1102 (2006).

113. D. S. Day, B. Zhang, S. M. Stevens, F. Ferrari, E. N. Larschan, P. J. Park, W. T. Pu, Comprehensive analysis of promoter-proximal RNA polymerase II pausing across mammalian cell types. Genome Biol 17, 120 (2016).

114. F. Ramirez, D. P. Ryan, B. Gruning, V. Bhardwaj, F. Kilpert, A. S. Richter, S. Heyne, F. Dundar, T. Manke, deepTools2: a next generation web server for deep-sequencing data analysis. Nucleic Acids Res 44, W160–165 (2016).

115. J. T. Robinson, H. Thorvaldsdottir, W. Winckler, M. Guttman, E. S. Lander, G. Getz, J. P. Mesirov, Integrative genomics viewer. Nat Biotechnol 29, 24–26 (2011).

116. D. M. Cable, E. Murray, L. S. Zou, A. Goeva, E. Z. Macosko, F. Chen, R. A. Irizarry, Robust decomposition of cell type mixtures in spatial transcriptomics. Nat Biotechnol 40, 517–526 (2022).

117. C. Oliveros J, Venny. An interactive tool for comparing lists with Venn diagrams. http://bioinfogp.cnb.csic.es/tools/venny/index.html, (2007).

