## Supplementary Materials for "Transcription-Coupled Chromatin Reinforcement Maintains the Mature Cardiomyocyte State"

Han-Hsuan Tang *et al.*

*Cheng-Fu Kao.

**This PDF file includes:**

Supplementary Text
Figs. S1 to S14
Tables S1 to S2

**Other Supplementary Materials for this manuscript include the following:**

Data S1 to S2

**Supplementary Text**

Full details of methods

#### Experimental Model and Study Participant Details

All animal experiments were conducted in accordance with the protocol approved by the Institutional Animal Care and Use Committee of Academia Sinica (Protocol #24-06-2184). Animals were maintained under specific pathogen-free conditions at 20 ± 2 °C with 50 ± 20 % humidity under a 14 h light/ 10 h dark cycle (lights on 07:00), with ad libitum access to chow diet (LabDiet) and reverse-osmosis water. To generate tamoxifen-inducible cardiomyocyte-specific *Rnf20* knockout mice (cKO), mice harboring two forward loxP sites flanking exon 3, 4, and 5 (*Rnf20*$^{flox/flox}$) (*1*) were crossed with *αMHC-MerCreMer*$^{+/-}$ (CreERT2) (*2*) transgenic mice. All mice were maintained on C57BL/6JNarl background. CreERT2 mice were used as controls. Only male mice were used to avoid potential confounding effects of female hormonal cycles.

#### Method Details

##### Genotyping

Genotyping was performed using genomic DNA extracted from toe tissues with the hot alkaline method (*3*). Briefly, the tissue was incubated in alkaline lysis buffer (25 mM NaOH and 0.2 mM disodium EDTA) for 10 min at 95°C, cooled on ice, and neutralized with an equal volume of 40 mM Tris-HCl. Genomic DNA was amplified by standard polymerase chain reaction (PCR) using Ex Taq DNA polymerase (TaKaRa) on a Veriti™ Thermal Cycler (Thermo Fisher). PCR products were resolved by agarose gel electrophoresis containing HealthView™ Nucleic acid stain (Genomics), and gel images were acquired using a GelDoc Go Gel Imaging System (Bio-Rad). Primers are available in the Table S1.

##### Drug treatment

Tamoxifen (Sigma-Aldrich) (20 mg/kg/day) was intraperitonially injected for 5 consecutive days to 8-10-week male mice to induce Cre-mediated recombination. To measure cell proliferation, 5-ethynyl-2'-deoxyuridine (EdU, Sigma-Aldrich) (50 mg/kg/day) was intraperitonially injected for 2 consecutive days and the hearts were collected 1 day after the final injection.

##### Western blotting

Protein was extracted from left ventricular (LV) tissues using radioimmunoprecipitation assay lysis buffer (150 mM NaCl, 1% NP-40, 0.5% sodium deoxycholate, 0.1% sodium dodecyl-sulfate, and 50 mM Tris-HCl, pH 8) supplemented with protease (Roche) and phosphatase (Sigma-Aldrich) inhibitor cocktails. Protein concentrations were determined using the detergent compatible protein assay (Bio-Rad). Equal amounts of protein were separated by sodium dodecyl-sulfate polyacrylamide gel electrophoresis and transferred onto polyvinylidene difluoride membranes. Membranes were blocked with 5% non-fat milk for 1h at room temperature and incubated with primary antibodies overnight at 4 °C. Following incubation with horseradish peroxidase-conjugated secondary antibodies for 1h at room temperature, immunoreactive bands were visualized by Immobilon ECL Ultra Western HRP Substrate (Millipore) and imaged using UVP ChemoStudio Imaging Systems (Analytik Jena). Total protein detected by 0.1% Ponceau S staining (Fluka) was generally used as the loading control; in specific experiments, GAPDH was used as an internal loading control.

Primary antibodies used were: anti-RNF20 (RRID: AB_10734436, Proteintech, 1:1000), anti-Cx43 (RRID: AB_297976, Abcam, 1:25,000), anti-YAP1 (RRID: AB_2650491, Cell Signaling, 1:1000), anti-p-YAP (S127) (RRID: AB_2650553, Cell Signaling, 1:1000), anti-GAPDH (RRID: AB_1080976, GeneTex, 1:1000), anti-α-tubulin (RRID: AB_1904178, Cell Signaling, 1:1000), and anti-detyrosinated α-tubulin (RRID: AB_869990, Abcam, 1:1000).

**Real-time quantitative PCR (RT-qPCR)**

RNA from LV tissues was extracted using TRIzol™ reagent (Thermo Fisher) according to the manufacturer's instructions. Briefly, the frozen tissue was pulverized into a fine powder using a mortar and pestle pre-chilled with liquid nitrogen before being transferred into TRIzol™ reagent for lysis. Genomic DNA was removed, and RNA was reverse-transcribed into complementary DNA using the QuantiTect Reverse Transcription Kit (QIAGEN). RT-qPCR was performed using the KAPA SYBR® FAST qPCR Master Mix (Kapa Biosystems) on a LighCycler480 instrument (Roche). *Gapdh* or 18S rRNA was used as the internal normalization control. Primers are available in the Table S1.

**Masson's Trichrome staining**

Adult hearts were briefly washed in pre-chilled phosphate-buffered saline (PBS), and fixed overnight in 4% paraformaldehyde at 4 °C. Fixed hearts were embedded in paraffin and sectioned at 5-µm to obtain four-chamber views. Masson's Trichrome staining was performed following standard protocols. Briefly, paraffin sections were deparaffinized in xylene, rehydrated through a graded ethanol series, and rinsed in deionized water. To enhance staining intensity, the sections were incubated in Bouin's solution. Nuclei were stained with Weigert's Iron Hematoxylin Solution, muscle fibers with Biebrich Scarlet-Acid Fuchsin solution, and collagen with Aniline Blue Solution. Finally, the sections were dehydrated through absolute ethanol and xylene before mounting. Images were captured with the Tissue Slide Scanner (TissueGnostics) equipped with a 20× objective.

**Adult murine cardiomyocyte isolation**

Adult cardiomyocytes were isolated using a Langendorff-free method (*4*). Mice were anesthetized with a mixture of isoflurane and oxygen. The descending aorta and vena cava were severed, and pre-chilled EDTA buffer (130 mM NaCl, 5 mM KCl, 0.5 mM $NaH_2PO4$, 10 mM HEPES, 10 mM glucose, 10 mM 2,3-Butanedione 2-monoxime, 10 mM taurine, and 5 mM EDTA, pH 7.8) was perfused through the right ventricle to arrest cardiac contractions. After clamping the ascending aorta, the heart was excised and further perfused through the left ventricle with EDTA buffer to remove residual blood, followed by perfusion with perfusion buffer (130 mM NaCl, 5 mM KCl, 0.5 mM $NaH_2PO4$, 10 mM HEPES, 10 mM glucose, 10 mM 2,3-Butanedione 2-monoxime, 10 mM taurine, and 1 mM $MgCl_2$, pH 7.8). The cleared heart was then digested by perfusion with pre-warmed (37 °C) perfusion buffer supplemented with collagenase B (0.5 mg/mL, Roche), collagenase D (0.5 mg/mL, Roche) and protease XIV (0.05 mg/mL, Sigma-Aldrich). Digested ventricular tissue was gently teased apart and triturated with wide-bore P1000 pipette tips, and enzymatic activity was quenched with 5% fetal bovine serum (Gibco). The cell suspensions were filtered through 200-µm cell strainers to remove undigested tissue and cardiomyocytes were separated from non-myocytes by centrifugation (20 x g for 3 minutes). Isolated cardiomyocytes were fixed in 4% paraformaldehyde for 15 minutes at room temperature with gentle mix on the nutator and washed with PBS. Fixed cardiomyocytes were adhered to Superfrost Plus slides using the Shandon Cytospin®4 centrifuge (Thermo Fisher) at 500 rpm for 5 min and subsequently processed for immunofluorescence staining.

**Cardiac tissue cryosectioning**

Adult hearts were perfused with pre-chilled PBS followed by 4% paraformaldehyde. The fixed hearts were then excised and cryoprotected by sequential incubation in 15% and 30% sucrose buffer. After cryoprotection, the hearts were embedded in Tissue-Tek® optimal cutting temperature compound (SAKURA), snap-frozen, and transversely sectioned at 10-µm. Tissue sections were mounted onto Superfrost Plus slides and subsequently processed for immunofluorescence staining or stored at -80 °C.

**Immunofluorescence staining, EdU incorporation assay, and TUNEL assay**

Frozen tissue sections or fixed isolated cardiomyocytes were permeabilized with 0.2% Triton X-100 for 45 min, blocked with 2% BSA for 1h at room temperature, and incubated with primary antibodies overnight at 4°C. Following PBS washes, samples were incubated with fluorophore-conjugated secondary antibodies for 1h at room temperature, counterstained with 4',6-diamidino-2-phenylindole (DAPI), mounted with Fluoromount-G® (SouthernBiotech), and sealed with coverslips.

EdU incorporation was detected using the Click-iT Plus EdU Cell Proliferation Kit (Thermo Fisher) according to the manufacturer's instructions. Briefly, following permeabilization and blocking, tissue sections were incubated with the Click-iT reaction cocktail, allowing fluorophore-conjugated picolyl azide to label incorporated EdU via click chemistry.

Apoptotic cells were detected using the TUNEL assay Kit (Elabscience) according to the manufacturer's instructions. Briefly, tissue sections were treated with proteinase K for 10 min, and fragmented DNA was labeled with fluorophore-conjugated dUTP using terminal deoxynucleotidyl transferase. Tissue sections treated with DNase I (200 U/mL) served as positive controls.

Primary antibodies used were: anti-cTnT (RRID: AB_11000742, Thermo Fisher, 1:500), anti-Cx43 (RRID: AB_297976, Abcam, 1:400), anti-YAP1 (RRID: AB_2650491, Cell Signaling, 1:100), anti-α-actinin (RRID: AB_11157538, Abcam, 1:800), anti-α-SMA-Cy3 (RRID: AB_476856, Sigma-Aldrich, 1:500), anti-dystrophin (RRID: AB_301831, Abcam, 1:100), anti-N-cadherin (RRID: AB_2313779, Thermo Fisher, 1:500), anti-Ki67 (RRID: AB_443209, Abcam, 1:1000), and anti-PCM1- Alexa Fluor® 647 (RRID: AB_2827155, Santa Cruz, 1:100)

All fluorescence images were acquired using a LSM880 confocal microscope (Zeiss) equipped with a 20× objective. Images of isolated cardiomyocytes used for sarcomere organization analysis were acquired using a 40× oil-immersion objective.

**Echocardiography and electrocardiography**

Cardiac function was evaluated by transthoracic echocardiography and surface electrocardiography (ECG). Mice were anesthetized with inhaled isoflurane in oxygen (1–1.5% at a flow rate of 1 L/min) and placed in the supine position on a heated platform to maintain body temperature throughout the procedures.

Transthoracic echocardiography was performed using a Vevo 3100 high-frequency ultrasound imaging system (FUJIFILM VisualSonics). LV systolic function was assessed using the PSLAX M-mode protocol, and measurements were analyzed with the Vevo Lab desktop software (FUJIFILM VisualSonics). LV internal diameter, anterior and posterior wall thicknesses at systole and diastole were measured. LV end-diastolic and end-systolic volumes, ejection fraction, and LV mass were subsequently calculated by the software.

Surface ECG recordings were acquired using a PowerLab 8/30 system equipped with an Animal Bio Amp (ADInstruments). Needle electrodes were inserted subcutaneously into the limbs according to the manufacturer's protocol to obtain standard limb ECG recordings. ECG signals were recorded for 10–15 min and analyzed using LabChart and Cardiac Axis software (ADInstruments). QRS interval was determined from the analyzed ECG traces.

**Cardiac nuclear isolation**

*Nuclear isolation for single-nucleus RNA-sequencing (snRNA-seq)*

Nuclear isolation for snRNA-seq was adapted from the previously published method with minor modifications (*5*). LV tissues were minced with fine-tip scissors and homogenized on ice using a Polytron™ handheld homogenizer (Kinematica) in lysis buffer (10 mM Tris-HCl, 10 mM NaCl, 3 mM $MgCl_2$, and 10% NP-40, pH 7.4). The reaction was stopped by adding wash buffer (2% BSA in PBS). The homogenate was filtered sequentially through 100-μm and 40-μm cell strainers and centrifuged at 1000 × *g* for 5 minutes at 4 °C to pellet the nuclei. The nuclei were resuspended in wash buffer and centrifuged again under the same conditions. The resulting pellet was resuspended in wash buffer containing 7-Aminoactinomycin D (7-AAD) (2 μg/mL) and incubated on ice for 25 minutes. 7-AAD-positive singlet nuclei were then sorted using a FACSAria III cell sorter (BD Bioscience). The quality of sorted nuclei was assessed using a EVOS M7000 microscope (Thermo Fisher).

*Nuclear isolation for assay for transposase-accessible chromatin using sequencing (ATAC-seq)*

Cardiomyocyte nuclei were purified for bulk ATAC-seq with modifications to a previously published protocol (*6*). LV tissues were finely minced with fine-tip scissors and homogenized on ice using a Polytron™ handheld homogenizer in lysis buffer A (5 mM $CaCl_2$, 3 mM $Mg(CH_3COO)_2$, 2 mM EDTA, 0.5 mM EGTA, and 10 mM Tris-HCl, pH 8.0). The homogenate was permeabilized in lysis buffer B (lysis buffer A supplemented with 0.1% Triton X-100), filtered through a 40-μm cell strainer, and centrifuged at 1,000 × g for 5 min at 4°C to pellet nuclei. Nuclei were resuspended in sucrose buffer (1 M sucrose, 3 mM $Mg(CH_3COO)_2$, and 10 mM Tris-HCl, pH 8.0) and centrifuged again under the same conditions. The resulting nuclei were incubated with anti-PCM1- Alexa Fluor® 647 antibody (RRID: AB_2827155, Santa Cruz, 1:250) for 30 min at room temperature to label cardiomyocyte nuclei, followed by staining with 7-AAD (2 µg/mL) in staining buffer (1% BSA in PBS) for 25 min on ice. Finally, 7-AAD-positive and PCM1-positive nuclei were purified using a FACSAria III cell sorter.

**SnRNA-seq library preparation**

Libraries for snRNA-seq (n =1 per genotype) were prepared using the Chromium Next GEM Single Cell 3′ Kit v3.1 Chemistry (10x Genomics) according to the manufacturer's protocol (CG000204), targeting approximately 10,000 nuclei per sample. Briefly, nuclei were partitioned into gel beads-in-emulsion (GEMs) using the Chromium Controller, where nucleus lysis, reverse transcription, and barcode incorporation were performed. Barcoded full-length cDNA was recovered, purified using Dynabeads MyOne SILANE beads (Thermo Fisher), amplified by PCR, and purified with 0.6× SPRIselect beads (Beckman Coulter). Sequencing libraries were generated through cDNA fragmentation, end repair, A-tailing, adaptor ligation, and sample indexing according to the manufacturer's protocol. Double-sided size selection was performed using SPRIselect beads with a 0.6×/0.8× bead ratio. Libraries were assessed for quality using a Bioanalyzer (Agilent Technologies) and sequenced on an Illumina NextSeq 2000 platform using paired-end sequencing.

**ATAC-seq library preparation**

ATAC-seq libraries (n = 3 per genotype) were prepared as previously described with minor modifications (*7, 8*). Briefly, PCM1-positive cardiomyocyte nuclei were incubated with Tagment DNA TDE1 enzyme (Illumina) at 37°C for 1 h in a thermomixer at 1,000 rpm. Tagmented DNA was purified using a MinElute PCR Purification Kit (QIAGEN) and initially amplified for five PCR cycles, followed by RT-qPCR to determine the additional number of amplification cycles required. Libraries were subsequently purified and assessed for quality using a Bioanalyzer. Paired-end (2 × 150 bp) sequencing was performed on an Illumina NovaSeq X Plus platform.

**Spatial transcriptomics library preparation**

Sample preparation and library construction for Visium HD spatial transcriptomics (10x Genomics; n = 1 per genotype) were performed according to the Visium HD Spatial Gene Expression Reagent Kits User Guide (CG000685). Formalin-fixed paraffin-embedded hearts were sectioned at 5-µm, stained with hematoxylin and eosin, and imaged using a Tissue Slide Scanner equipped with a 20× objective. Following tissue destaining and decrosslinking, mRNA transcripts were detected using the Visium Mouse Transcriptome Probe Set v2.0 (10x Genomics). Hybridized and ligated probes were transferred in situ from the tissue sections onto the Visium HD slide using the CytAssist instrument (10x Genomics). Captured probes were subsequently barcoded and released for library preparation. Libraries were initially amplified for 10 PCR cycles, purified using 1.2× SPRIselect beads, and subjected to RT-qPCR to determine the additional amplification cycles. Following PCR amplification, libraries were indexed for multiplexing, purified using 0.85× SPRIselect beads, assessed for quality using a Bioanalyzer, and sequenced on an Illumina NextSeq 2000 platform using paired-end sequencing.

**SnRNA-seq analysis**

*Preprocessing*

FASTQ files were aligned to the mm10 reference genome using STARSolo (v2.7.11a) (*9, 10*) with the parameter --soloFeatures Full on Galaxy (*11*). Data preprocessing was performed using Trailmaker™ (https://app.trailmaker.parsebiosciences.com/; Parse Biosciences, analysis completed on May 17, 2024), which included multiple steps to remove low-quality nuclei: (1) Classifier filter—empty droplets were removed using *emptyDrops* (*12*) with a false discovery rate < 0.01; (2) Number of genes vs. transcripts filter—nuclei exhibiting under- or over-amplification were excluded based on predictions from the R function *predict*; and (3) Doublet filter—potential doublets were identified and removed using *scDblFinder* with default settings (*13*). Subsequent analyses were performed using Seurat following the standard workflow (*14*). Gene expression values were log-normalized, and the top 2,000 highly variable features were selected for downstream analyses. Linear dimensionality reduction was conducted using principal component analysis (PCA) with the first 23 PC. Harmony method was adopted for data integration. Clustering was performed using the Louvain algorithm with a resolution of 0.8. Uniform Manifold Approximation and Projection with a minimum cosine distance of 0.3 was used for two-dimensional embedding. A second round of *scDblFinder* (v1.22.0) (*13*) analysis was applied to the processed Seurat object, which identified a cluster co-expressing macrophage and endothelial cell markers as a putative doublet population. This cluster was removed prior to downstream analyses.

*Subclustering of cardiomyocytes and fibroblasts*

Cardiomyocytes and fibroblasts were subset, re-clustered, and integrated using procedures analogous to those applied during the initial Seurat preprocessing, with the following modifications. Cardiomyocytes were analyzed using 30 PCs for dimensionality reduction and a Louvain clustering resolution of 0.8, after which two clusters containing fewer than 50 cells were excluded from downstream analyses. Fibroblasts were analyzed using 20 PCs and a Louvain clustering resolution of 0.3, after which one fibroblast cluster containing fewer than 50 cells was merged with cluster 0.

*Potency scoring*

CytoTRACE (v0.3.3) (*15*) was adopted to infer the relative differentiation state of cardiomyocytes based on the number of detectably expressed genes per cell. The analysis was performed using the raw count matrix extracted from the Seurat object, following default settings.

*Single-cell gene signature scoring*

AUCell (v1.30.1) is a method used to quantify gene set activity at the single-cell level (*16*). Briefly, genes in each cell were ranked according to their expression levels. The top 5% of ranked genes was then used for the recovery analysis, in which the x-axis represents the gene rank and the y-axis represents the cumulative number of genes recovered from the specified gene set. The area under this recovery curve was calculated and used as the AUCell score.

*Pseudotime analysis*

Monocle3 (v1.4.26) (*17*) was used to construct single-cell trajectory of cardiomyocyte dedifferentiation and to identify pseudotime-dependent genes following the standard workflow. A cell from CM2 was selected as the trajectory root and differential expression analysis along the trajectory was performed using graph-autocorrelation framework, which applies Moran's I. Genes with a $q$-value < 0.05 were considered significant. Pseudotime-dependent genes were subsequently grouped into modules using the Louvain algorithm. For simplicity, these modules were further clustered into three groups using hierarchical clustering implemented in the ComplexHeatmap (*18, 19*).

*Cross-referencing with the published snRNA-seq dataset*

Cardiomyocyte populations were cross-referenced with a published snRNA-seq dataset from neonatal murine hearts following myocardial infarction (n = 1 per treatment group per developmental stage) (*20*) to assess whether the dedifferentiated cardiomyocytes identified in the present study resembled injury-related or regeneration-associated cell states. The published dataset was preprocessed using the Trailmaker$^{TM}$ pipeline as described above. Because substantial mitochondrial contamination was observed, nuclei with mitochondrial transcript content exceeding three median absolute deviations above the median were removed. Cardiomyocytes were then subclustered following procedures analogous to those described in the subclustering section, using 30 PC for dimensionality reduction and a Louvain clustering resolution of 0.4. Cardiomyocyte states were annotated based on marker genes identified in the original study (*20*). To compare datasets, the *FindTransferAnchors* function in Seurat was used to transfer cell-state labels from the reference dataset (neonatal cardiomyocytes) to the query dataset (cardiomyocytes from the present study).

*Overrepresentation analysis*

To annotate the biological functions of gene sets—including pseudotime-dependent genes and activated YAP1 target genes (*21*)—overrepresentation analysis was performed using the DAVID (v2023q4) (*22, 23*). Annotation terms with a false discovery rate < 0.05 were considered significantly enriched.

#### ATAC-seq analysis

*Preprocessing*

Preprocessing was carried out on Galaxy following their analysis protocol (*11, 24, 25*). Quality control was performed with FastQC (v0.12.1) (*26*). Low-quality reads and Nextera adapter sequences were removed using Cutadapt (v5.2) (*27*) with the parameters: -a CTGTCTCTTATACACATCT --nextseq-trim 20 --minimum-length 20. Reads were aligned to the mm10 reference genome using Bowtie2 (v2.5.5) (*28, 29*) with the parameters: -I 0 -X 1000 --very-sensitive. Low-quality (MAPQ < 30), unpaired, and mitochondrial reads were removed using BAMTools (v2.5.3) (*30*). Duplicate reads were identified and removed with Picard (v3.1.1) (*31*). Tn5 offset was adjusted and reads in blacklisted regions (*32*) were removed with *alignmentSieve* --ATACshift from deepTools (v3.5.4) (*33*). Published ATAC-seq data from adult cardiomyocytes expressing constitutively active YAP1 (YAP5SA, n = 2 per genotype) (*21*) were processed using the same pipeline described above.

*Peak calling and merging for consensus peak set*

Shifted BAM files were converted to single-end BED format using BEDTools (v2.31.1) (*34*). Peaks were called using *callpeak* function from MACS2 (v2.2.91)(*35, 36*) with the following parameters: --format BED --gsize 1.87e9 --nomodel --extsize 150 --shift -75 --qvalue 0.05 --call-summits --nolamda --keep-dup all. An iterative overlap peak-merging strategy (https://github.com/corceslab/ATAC_IterativeOverlapPeakMerging) was used to generate a merged peak set using default settings (*37*).

*Differentially accessible peaks analysis*

The BEDTools (*34*) *multicov* function was used to quantify the number of transposition events per peak for each sample. Peaks with at least 50 counts in at least one sample were retained for downstream analyses. Batch effects were corrected using the RUVseq (v1.42.0) (*38*). Differential accessibility analysis was performed using DESeq2 (v1.48.2) with default settings (*39*). Peaks with adjusted $p$-value < 0.05 were considered significant.

*Genomic annotation*

Genomic regions corresponding to differentially accessible peaks were annotated using the *annotatePeak* function from the ChIPSeeker package (v1.44.0) (*40*). The UCSC mm10 transcript annotation file was used as the reference. Promoter regions were defined as ±2 kb from the transcription start site.

*Chromatin state modeling*

Chromatin states were inferred using ChromHMM (v1.27) following the standard workflow (*41*). Processed chromatin immunoprecipitation sequencing (ChIP-seq) BAM files for multiple histone modifications and CTCF, along with corresponding input controls from adult murine heart tissues (n = 2), were obtained from ENCODE (*42-44*) and used as input for model training. Briefly, BAM files were binarized using the *BinarizeBam*, and a 12-state chromatin model was learned using the *LearnModel*. This approach enabled the identification of established regulatory elements, including Polycomb-repressed regions (H3K27me3-positive) (*45*), transcribed regions (H3K36me3-positive) (*46*), poised enhancers (H3K4me1-positive, H3K27ac-/H3K4me3-negative) (*47*), active enhancers (H3K27ac-positive, H3K4me3-negative) (*47*), promoters (H3K4me3-positive) (*48*), and insulators (CTCF-positive) (*49*). The *OverlapEnrichment* function was subsequently used to compute fold enrichment of chromatin states across genomic annotations as well as differentially accessible peaks.

*Region-based enrichment analysis*

Region-centric gene set enrichment analysis was performed using the rGREAT (v2.10.0) (*50*), which implements the GREAT algorithm (*51*) to associate regulatory regions with Gene Ontology Biological Process terms. Analyses were conducted using default settings, and the merged peak set served as the background. Terms with adjusted $p$-values < 0.1 in at least one genotype were preserved. The top 100 differential terms, ranked by the absolute difference in log10–adjusted $p$-values, were selected for network analysis. Graph-based semantic similarity among enriched terms was computed using the GOSemSim (v2.34.0) to construct pathway similarity networks (*52*). Network visualization and refinement were performed in Cytoscape (v3.10.4) (*53*). The complete lists of enriched and the top 100 differential terms are available in Data S2.

*De novo motif enrichment analysis*

Enriched motifs within differentially accessible peaks were identified using the HOMER (v5.1) with the *findMotifsGenome.pl* function and default settings (*54*). The merged peak set was used as the background for motif enrichment analysis.

*Footprinting analysis*

Footprinting analysis was performed with TOBIAS (v0.14.0) following the standard workflow (*55*). Unshifted ATAC-seq BAM files from biological replicates were first merged for each genotype. Tn5 insertion bias was corrected using the *ATACorrect* function. Footprint scores were then computed using the *FootprintScores* function. Finally, transcription factor binding prediction and differential binding analysis were conducted with *BINDetect*. Nonredundant vertebrate core transcription factor binding motifs were obtained from the JASPAR 2024 database (*56*).

**Bulk RNA-seq analysis**

*Preprocessing*

RNA-seq FASTQ files for murine cardiomyocytes collected at embryonic day 14.5 and postnatal days 1, 4, 7, 14, and 56 (*57*) (n = 3) and human heart-failure cardiomyocyte nuclei (*58*) (n = 3 for non-failing and 4 for failing group) were accessed from previously published datasets. Preprocessing was performed on Galaxy following standard procedures (*11, 25, 59*). Quality control was performed with FastQC (*26*). For the mouse datasets, low-quality reads were removed using Cutadapt (*27*) with the parameters --quality-cutoff 20 --minimum-length 20. Reads were aligned to the mm10 or hg38 reference genome, as appropriate, using STAR (v2.7.11a) (*10*) with the parameters: --sjdbOverhang set to 149 (mouse) or 49 (human) --quantMode GeneCounts. Gene-level read counts were obtained using *featureCounts* from subread (v2.0.3) with default parameters (*60, 61*). For the human dataset, both exonic and intronic reads were quantified because nuclear RNA predominantly comprises nascent transcripts.

*Construction of maturation-associated gene sets*

For murine cardiomyocytes RNA-seq, genes with counts per million ≥ 1 in at least three samples were retained for downstream analyses. Count normalization was performed using the Trimmed Mean of M values method implemented in the NOISeq (v2.48.0) (*62, 63*). Normalized counts were analyzed using maSigPro (v1.76.0)(*64*) to identify temporally dynamic expression profiles across cardiomyocyte maturation. Significant genes ($R^2 > 0.6$) were partitioned into two clusters: one containing genes whose expression increased during maturation (mature gene set) and the other containing genes whose expression decreased over time (immature gene set). Lists of the maturation-associated genes are available in Data S1.

*Differentially expressed genes analysis*

For human cardiomyocyte nuclei RNA-seq, differentially expressed genes were identified with DESeq2 (*39*) with default settings and genes with adjusted *p*-values < 0.05 were considered significant.

**ChIP-seq analysis**

*Preprocessing*

ChIP-seq FASTQ files for H2Bub from murine hearts at postnatal day 28 (n = 2) (*65*), H3K4me3 and H3K27ac from adult murine cardiomyocytes nuclei (n = 2) (*66*), RNA polymerase II (Pol II) from adult murine cardiomyocytes (n = 1) (*67*), and H3K4me3 from human heart failure cardiomyocyte nuclei (n = 2 for non-failing and 3 for failing group) (*58*) were accessed from previously published datasets. Preprocessing was carried out on Galaxy (*11*). Quality control of the reads was performed with FastQC (*26*). Adapter sequences were removed from the human dataset using Cutadapt (*27*), whereas low-quality bases were trimmed from the Pol II dataset using the parameters --quality-cutoff 20 --minimum-length 20. No additional trimming was required for the remaining datasets. Reads were aligned to the mm10 or hg38 reference genome, as appropriate, using Bowtie2 (*28, 29*) with the parameters -I 0 -X 1000 --very-sensitive. Low-quality (MAPQ < 30), unpaired, and mitochondrial reads were removed using BAMTools.(*30*) Duplicate reads were identified and removed with Picard (*31*).

*H3K4me3 peak identification and annotation*

Peaks were called using *callpeak* function from MACS2 (*35, 36*) with the following parameters: --format BAMPE --gsize 1.87e9 (or 2.7e9 for human) --broad. Blacklisted regions (*32*) were removed and common peaks between replicates were identified using the *intersect* function from BEDTools (*34*). ChIPSeeker (*40*) was used to annotate the common peaks and the maximal peak breadth was used for each annotated gene.

**Tracks visualization**

Biological replicates were merged at the BAM level prior to visualization. For ATAC-seq, shifted BAM files were converted to reads-per-million-normalized bigWig files using the *bamCoverage* function in deepTools (*33*). For ChIP-seq, merged BAM files were normalized to their corresponding input controls using the *bamCompare* function in deepTools (*33*) with reads per kilobase per million normalization to generate ratio bigWig files. Blacklisted genomic regions (*32*) were excluded from all analyses. These bigWig files were subsequently used to generate heatmaps and metagene profile plots of signal distributions across genomic regions using the *computeMatrix* and *plotHeatmap* functions in deepTools (*33*). Injury-associated accessible peaks for metagene profile plots were retrieved from a previously published dataset (*68*) and converted from the mm9 to the mm10 genome assembly using the liftOver tool (UCSC Genome Browser) (*69*). Integrative Genomics Viewer (v2.19.7) (*70*) was used to visualize normalized sequencing tracks, differentially accessible peaks, and chromatin states at specific loci.

**Spatial transcriptomics analysis**

*Preprocessing*

Spatial transcriptomic data were processed using the Space Ranger count pipeline (v4.0.1) implemented in the 10x Genomics Cloud Analysis platform to generate feature-bin matrices. Reads were aligned to the GRCm39 reference genome using default parameters. Downstream analyses were performed in Seurat using the 8-μm binning resolution following the standard workflow (*14*). Non-ventricular bins, identified in Loupe Browser (v9), were excluded. Low-quality bins, including most residual erythrocyte-containing bins, were removed based on the following criteria: median unique molecular identifier counts <100, median detected genes <60 for CreERT2 or <50 for cKO, and mitochondrial transcript percentage >40%. After quality control, 202,660 and 223,303 bins were retained from the CreERT2 and cKO heart, respectively. Data were normalized using *SCTransform* with regression of mitochondrial transcript percentage. Mitochondrial genes were excluded from downstream clustering analyses to minimize clustering driven by regional differences in mitochondrial gene expression. *SketchData* function with the LeverageScore sampling method was used to subsample 50,000 bins from each dataset. PCA was performed, datasets were integrated using Harmony, and the resolution was set at 0.8 for Louvain clustering. Cluster identities and dimensional reductions learned from the sketched dataset were subsequently projected to the full dataset using the *ProjectIntegration* and *ProjectData* functions.

A total of 21 clusters were initially identified. Four clusters displaying stripe-like spatial patterns suggestive of technical artifacts, three rare populations (<50 bins), and one erythrocyte cluster identified by canonical marker gene expression were excluded from downstream analyses, resulting in 13 high-quality clusters.

*Reference-based spatial deconvolution analysis*

Because each 8-µm bin may contain transcripts derived from more than one cell, spatial deconvolution was performed using the Robust Cell Type Decomposition (RCTD) algorithm from the spacexr package (v1.4.0) (*71*) in doublet mode with our snRNA-seq dataset as the reference. RCTD assigns up to two cell types to each bin together with their corresponding weights, representing the estimated contribution of each cell type.

*Spatial gene signature scoring and distance-based analysis*

To examine the spatial distribution of dedifferentiation-associated transcriptional programs, AUCell scoring (*16*) was performed as described above using snRNA-seq-derived pseudotime-dependent gene sets. Cardiomyocyte-enriched bins were first defined as those with an RCTD cardiomyocyte weight >0.8. Among these bins, those from cKO with AUCell scores for trajectory Group 2 or Group 3 exceeding the 99th percentile of the corresponding scores in CreERT2 were designated as Group $2^{High}$ and Group $3^{High}$, respectively. To determine whether Group $3^{High}$ bins preferentially localized to spatial cluster 8, distance-based analysis was performed modified from a previously published method (*72*) (https://github.com/10XGenomics/HumanColonCancer_VisiumHD/blob/main/Methods/AuxFunctions.R). Briefly, bins with at least 25 neighboring bins belonging to spatial cluster 8 were defined as the core region, and the surrounding bins were stratified into concentric zones extending 50 µm, 100 µm, 200 µm, or >200 µm from the core (https://doi.org/10.5281/zenodo.21700369).

#### Quantification and Statistical Analysis

##### Quantification of protein level for Western blotting

Immunoreactive band intensities were quantified using Fiji (National Health Institute, v1.54), whereas Ponceau S-stained band intensities were quantified using Image Lab software (Bio-Rad, v6.1). Target protein band intensities were normalized to the corresponding loading controls. Fold changes were calculated by dividing the normalized band intensities by the mean normalized intensity of the CreERT2 group.

##### Quantification of gene expression for RT-qPCR

Cp values of the target genes were normalized to those of the corresponding internal controls to obtain ΔCp values. The ΔCp value for each sample was then normalized to the mean ΔCp value of the CreERT2 group to obtain ΔΔCp values. Fold changes in gene expression were calculated using the $2^{-\Delta\Delta Cp}$ method.

#### Quantification of fibrotic area

Fibrosis was measured using the images from Masson's Trichrome-stained tissue sections. Fibrotic area was quantified by measuring the ratio of collagen-positive area (blue staining) to muscle area (red staining) in both the LV free wall and interventricular septum using Fiji.

#### Quantification for fluorescence images

*Cardiomyocyte nucleation and nuclear area*

Because cardiomyocyte proliferative capacity is associated with mononucleation (*73*), and mature murine cardiomyocytes are predominantly bi- or multinucleated (*74*), we further examined cardiomyocyte nucleation status using isolated cardiomyocytes. Nuclei were counterstained with DAPI. Overlapping cells were excluded from the analysis, and at least 200 cardiomyocytes were evaluated per sample. Maximum-intensity z-projection images were generated for each field of view (FOV) with Fiji. Cells containing one, two, or three or more nuclei were classified as mononucleated, binucleated, or multinucleated, respectively. For nuclear area measurements, DAPI signals were thresholded to remove background prior to quantification using Fiji.

*Quantification of nuclear YAP1, proliferative cardiomyocytes, and apoptotic nuclei*

To quantify nuclear YAP1 localization, cardiomyocyte proliferation, and apoptosis, at least four FOV from the LV myocardium were acquired for each sample. Maximum-intensity z-projection images were generated for each FOV and cardiomyocyte nuclei were identified as nuclei enclosed by cTnT-positive sarcomeres, dystrophin-labeled sarcolemma, or positive for PCM1 using Fiji. YAP1 was classified as nuclear when the fluorescence intensity within the nucleus exceeded that of the surrounding cytoplasm. EdU-positive and Ki67-positive cardiomyocyte nuclei and TUNEL-positive nuclei were manually counted and expressed as percentages of the corresponding total nuclear populations.

*Sarcomere organization analysis*

Sarcomeres were immunolabeled using anti-cTnT antibody in tissue sections or anti-α-actinin antibody in isolated cardiomyocytes as described above. Maximum-intensity z-projection images were generated for tissue sections and isolated cardiomyocytes using Fiji. For tissue sections, longitudinal regions of the left ventricle were analyzed using SotaTool (*75, 76*). SotaTool parameters were set as follows: image resolution = 2.4089 pixels/µm for tissue sections or 4.8177 pixels/µm for isolated cardiomyocytes; background subtraction = enabled; segmentation = enabled (4 × 4) for tissue sections; and "disregard lowest GLCM-bin" = 3. For tissue sections, images of yielding numerical sarcomere organization scores were classified as having detectable sarcomere periodicity, whereas images returning NA values were classified as having undetectable sarcomere periodicity. For isolated cardiomyocytes, sarcomere length and sarcomere organization scores were quantified using SotaTool.

*Quantification of a-smooth muscle actin (α-SMA)*

At least four FOV from the LV myocardium were acquired for each sample. Maximum-intensity z-projection images were generated for each FOV and analyzed using Fiji. The α-SMA-positive area within α-actinin-positive cardiomyocytes was divided by the total α-actinin-positive area and multiplied by 100 to derive the percentage of α-SMA-positive area.

*Quantification of Cx43 at intercalated discs (IDs)*

Quantification of Cx43 localization at IDs was adapted from a previously published method (*77, 78*) with minor modifications using Fiji. N-cadherin–positive regions were defined as IDs. Briefly, N-cadherin fluorescence images were stacked, background-subtracted using a rolling-ball radius of 10 pixels, converted to 8-bit depth, and uniformly thresholded using the default method to generate a binary ID mask. The Fiji “Image Calculator” function with the “AND” operation was then used to extract Cx43 signals within the N-cadherin mask, yielding an ID-restricted Cx43 image. Mean fluorescence intensity was measured for both the ID-restricted Cx43 image and the corresponding original Cx43 image. The Cx43 ID co-localization index was calculated as the ratio of the mean fluorescence intensity of ID-restricted Cx43 to that of total Cx43. All values were normalized to the mean value of the CreERT2 group.

**Quantification for spatial transcriptomics**

*Heatmap visualization of cell-type enrichment*

Heatmap was generated with R for visualization of cell-type distribution among spatial clusters. The mean RCTD weight of each cell type was calculated across all bins within each spatial cluster and standardized by z-score across clusters to facilitate comparison of relative enrichment.

*AUCell score density plot*

The distribution of AUCell scores among cardiomyocyte-enriched bins was visualized using kernel density plots generated with the *geom_density* function in ggplot2 (*79*). Median AUCell scores for each genotype and the 99th percentile of the scores in CreERT2 were indicated by dashed vertical lines.

*Quantification of Group $3^{High}$ bins*

The percentage of Group $3^{High}$ bins in the cKO heart was calculated by dividing the number of Group $3^{High}$ bins by the total number of cardiomyocyte-enriched bins and multiplying by 100 for both the cluster 8 core and each surrounding concentric zone.

**Pausing index (PI) calculation**

PI was calculated from Pol II ChIP-seq data following the published Pausing Index Calculator pipeline (https://github.com/MiMiroot/PIC) (*80*). Briefly, Pol II read counts were normalized to the total number of mapped reads, and the normalized input signal was subtracted from the normalized Pol II signal. PI was then calculated by dividing the length-normalized read counts within the promoter region (50 bp upstream to 300 bp downstream of the transcription start site) by the length-normalized read counts within the gene body (300 bp downstream of the transcription start site to 3 kb downstream of the transcription end site).

**Statistical analysis**

Statistical analyses were performed using R (v4.5.0) and GraphPad Prism (v8). Comparisons between two groups were performed using an unpaired Student's t-test, with Welch's correction applied when variances were unequal, or the Mann–Whitney U test for nonparametric data. Comparisons among more than two groups were performed using one-way ANOVA followed by Tukey's multiple-comparisons test for parametric data or the Kruskal–Wallis test followed by Dunn's multiple-comparisons test for nonparametric data. Cell-type enrichment among spatial clusters was assessed by testing whether the RCTD weights of a given cell type were significantly higher in one cluster than in all remaining clusters using a one-sided Wilcoxon rank-sum test (alternative = "greater"), followed by Bonferroni correction for multiple testing. Enrichment of Group $3^{High}$ bins across distance-resolved zones in spatial transcriptomics was evaluated using the one-sided Cochran–Armitage trend test. The genotype × time interaction was assessed using a mixed-effects model for longitudinal changes in ejection fraction, QRS interval, LV mass-to-body-weight ratio, and body weight between groups. Survival curves were compared using the log-rank (Mantel–Cox) test. Categorical variables were compared using the chi-square test. A two-sided $p$ value < 0.05 was considered statistically significant. Sample sizes, statistical analyses, and definitions of error bars are specified in the corresponding figure legends or Method Details. No statistical methods were used to predetermine sample size. No data were excluded from the analyses, except for the published human heart failure H3K4me3 ChIP-seq dataset as one of the replicates from the non-failing group exhibited substantially reduced promoter-associated enrichment and poor reproducibility relative to the remaining biological replicates. The experiments were not randomized, and investigators were not blinded to group allocation or outcome assessment.

Fig. S1

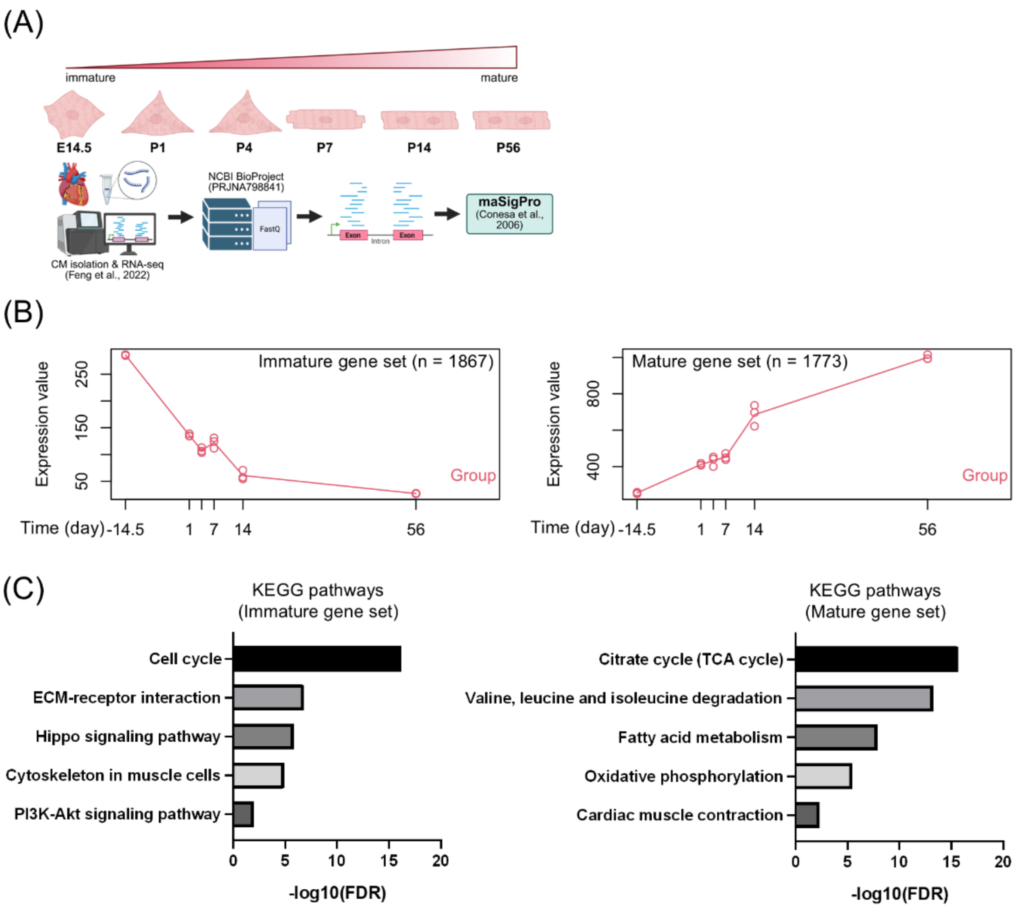


**Fig. S1.**

Construction of maturation-associated gene sets. Related to Figure 1.

(**A**) Schematic for analysis of maturation-associated genes.

(**B**) Time-dependent gene expression profiles for immature (left) and mature (right) gene set. The experiments were performed with n = 3 biological replicates.

(**C**) Overrepresentation analysis against KEGG pathways for immature (left) and mature (right) gene set. FDR, false discovery rate.

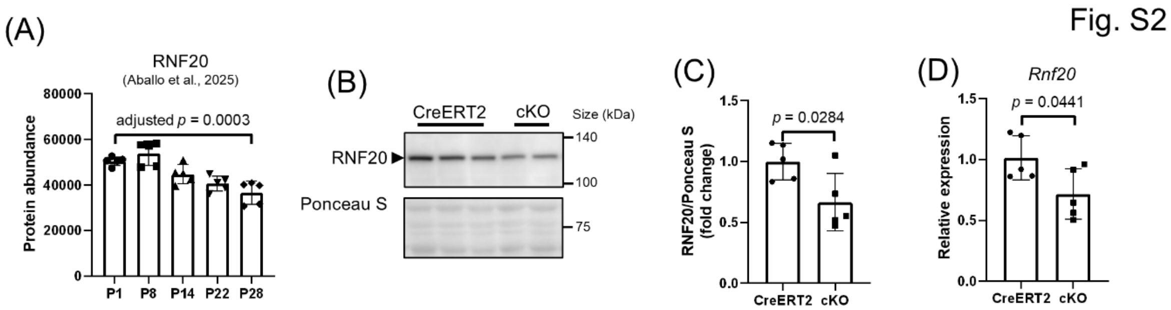


**Fig. S2.**

RNF20 depletion in adult hearts. Related to Figure 1.

(**A**) Protein abundance of RNF20 from murine ventricular tissues. Data were obtained from Aballo et al. (*81*). The data were generated with n = 5 biological replicates and are presented as mean ± s.d.. One-way ANOVA with Tukey's test.

(**B**) Western blot analysis of RNF20 from left ventricular (LV) tissues at 2 weeks post-tamoxifen (wpt). Total protein stained with Ponceau S was used as normalization control.

(**C**) Quantification of (B). The data were generated with n = 5 biological replicates and are presented as mean ± s.d.. Unpaired two-tailed t tests.

(**D**) Real-time quantitative polymerase chain reaction (RT-qPCR) analysis of *Rnf20* from LV tissues at 2 wpt. *Gapdh* was used as normalization control. The data were generated with n = 5 biological replicates and are presented as mean ± s.d.. Unpaired two-tailed t tests.

Fig. S3

(A)

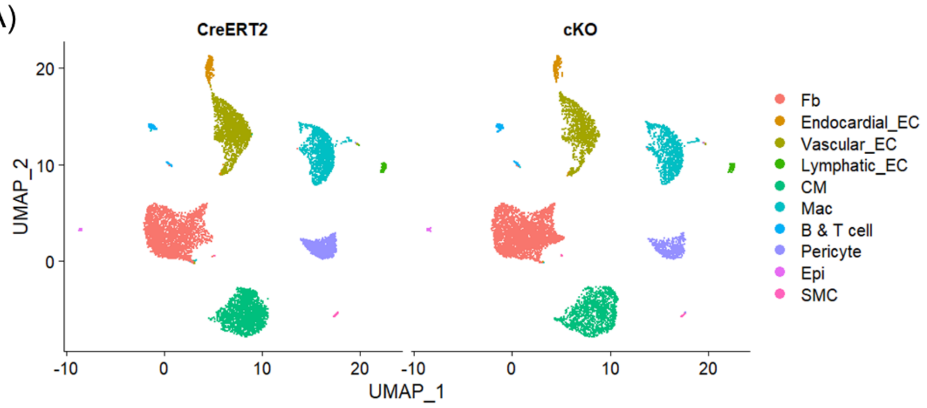


(B)

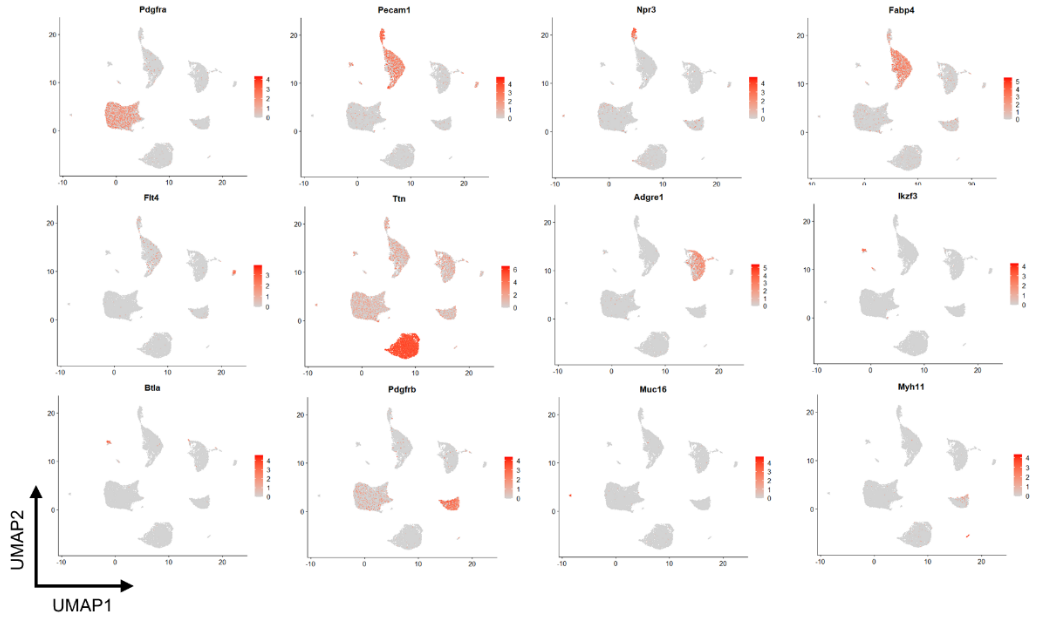


(C)

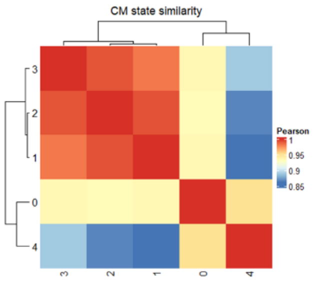


(D)

| Cell state | GO:BP | FDR | Associated genes |
|---|---|---|---|
| CM1 | fatty acid beta-oxidation | 2.00E-06 | *Decr1*, *Abcd3*, *Acaa2*, *Acadm*, *Cpt1a*, *Cpt2*, *Ech1*, *Eci1*, *Hadha* |
| CM2 | regulation of transcription by RNA polymerase II | 4.18E-02 | *Ecsit*, *Klf13*, *Phf8*, *2610008E11Rik*, *Brwd3*, *Ezh2*, *Foxk1*, *Nucks1*, *Pcgf3*, *Ppm1d*, *Satb1*, *Sp1*, *Zfp160*, *Zfp260*, *Zfp369*, *Zfp397*, *Zfp672* |
| CM3 | cardiac myofibril assembly | 2.48E-02 | *Adprhl1*, *Actc1*, *Cavin4*, *Csrp3* |

**Fig. S3.**

Homeostatic cardiomyocyte states exhibit distinct transcriptional and functional identities. Related to Figure 1.

(**A**) UMAP visualization of cardiac clusters in CreERT2 (left) and cKO (right) hearts at 2 wpt. Fb, fibroblast; CM, cardiomyocyte; EC, endothelial cell; Mac, macrophage; Epi, epicardial cell; SMC, smooth muscle cell.
(**B**) UMAP visualization of known marker genes among cardiac cell types.
(**C**) Cardiomyocyte state similarity determined by Pearson correlation coefficient with the top 2000 highly variable genes.
(**D**) Overrepresented Gene Ontology terms and their associated genes in CM1, CM2, and CM3. GO:BP, Gene Ontology: Biological Processes.

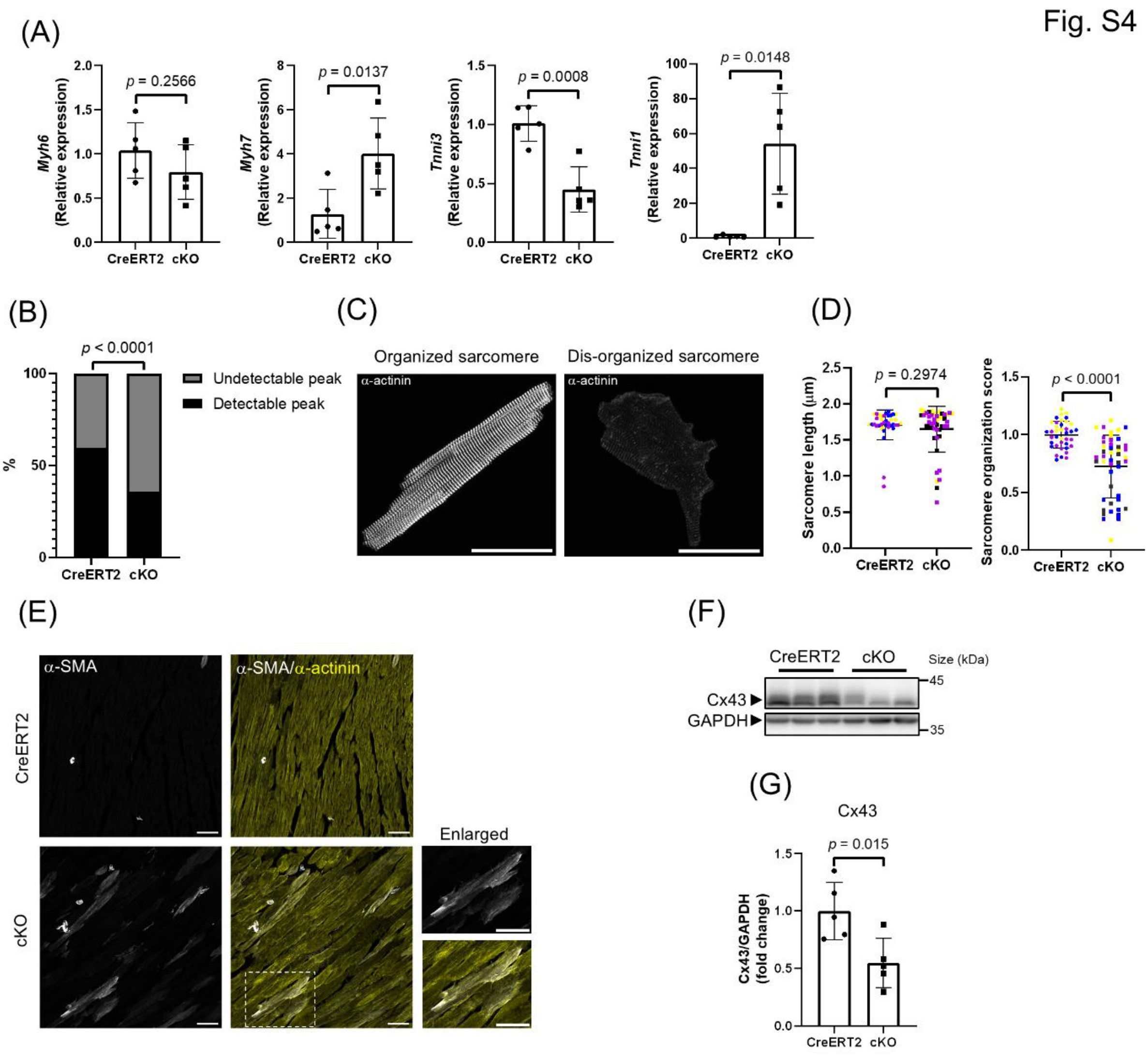


**Figure S4.**

Extended analyses supporting RNF20-dependent maintenance of molecular and structural features in mature cardiomyocytes. Related to Figure 2.

(**A**) RT-qPCR analysis of maturation markers from LV tissues at 2 wpt. *Gapdh* was used as normalization control. The data were generated with n = 5 biological replicates and are presented as mean ± s.d.. Unpaired two-tailed t test. Welch's correction was performed for *Tnni1*.

(**B**) Quantification of sarcomere organization by immunofluorescence analysis of cardiac troponin T (cTnT) at 2 wpt. The data were generated with n = 3 (CreERT2 with 153 sub-images) and 4 (cKO with 221 sub-images) biological replicates. Chi-square test.

(**C**) Sarcomere organization evaluation by immunofluorescence analysis of α-actinin with isolated cardiomyocytes at 4 wpt. Cardiomyocytes with organized (left) and dis-organized (right) sarcomere are shown. Scale bar = 50 μm.

(**D**) Quantification of (C). Data were generated with n = 3 (CreERT2 with 38 cells) and 4 (cKO with 46 cells) biological replicates and are presented as mean ± s.d.. Data points with the same color represent measurements from the same heart. Unpaired two-tailed t test with Welch's correction.

(**E**) Immunofluorescence analysis of α-SMA at 6 wpt. Scale bar = 50 μm.

(**F**) Western blot analysis of Cx43 from LV tissues at 2 wpt. GAPDH was used as normalization control.

(**G**) Quantification of (F). The data were generated with n = 5 biological replicates and are presented as mean ± s.d.. Unpaired two-tailed t test.

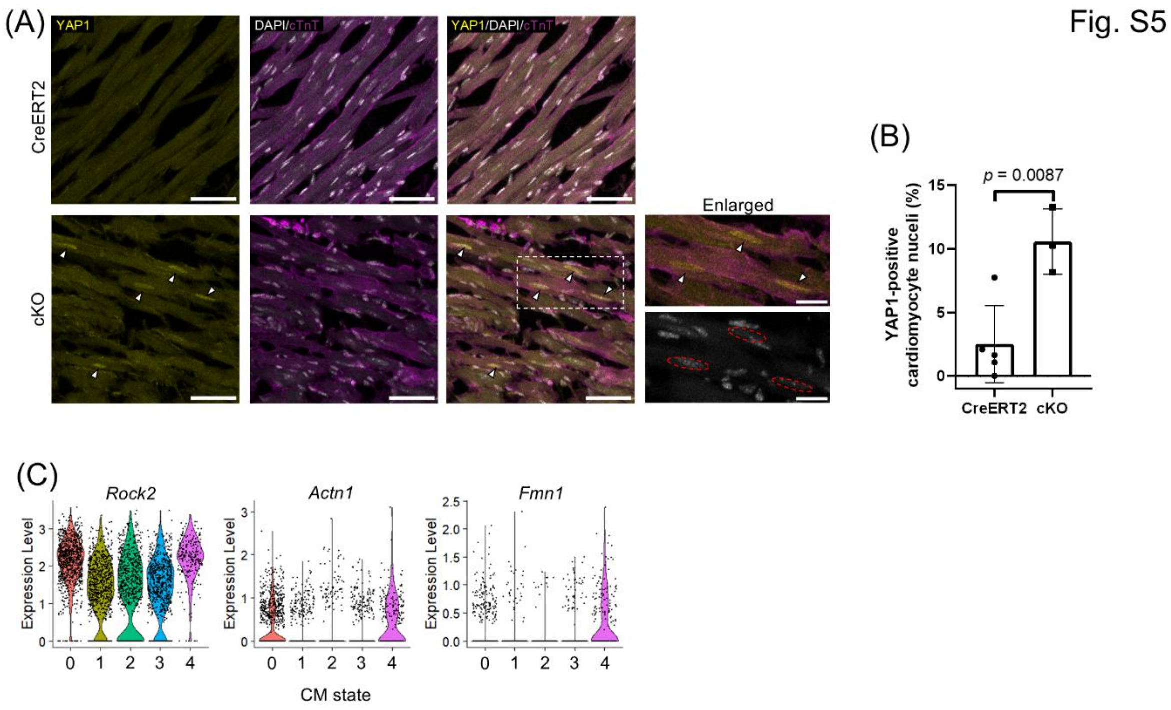


**Figure S5.**

Extended analyses supporting YAP1 activation in RNF20-deficient cardiomyocytes. Related to Figure 3.

(**A**) Immunofluorescence analysis of YAP1 at 2 wpt. Arrows show nuclear YAP1. YAP1 was regarded nuclear if the intensity of YAP1 in the nucleus was stronger than the cytoplasm. Cardiomyocyte nuclei are encircled in the insets for better identification. Scale bar = 50 µm (20 µm for insets).

(**B**) Quantification of nuclear YAP1 from (A). Data were generated with n = 5 (CreERT2) and 3 (cKO) and are presented as mean ± s.d.. Unpaired two-tailed t test.

(**C**) Violin plots showing expression of representative upregulated YAP1 targets involving in actin cytoskeleton organization across individual nuclei in cardiomyocyte states.

Fig. S6

(A)

| Genotype | Replicate # | Unique concordant alignment rate (%) | Read pair after deduplication | Peak number | Merged peak number | |
|---|---|---|---|---|---|---|
| CreERT2 | 1 | 78.53 | 60,735,081 | 450,202 | 44,117 | 48,796 |
| | 2 | 78.71 | 50,172,353 | 374,496 | | |
| | 3 | 78.13 | 45,046,778 | 363,385 | | |
| cKO | 1 | 80.61 | 76,285,715 | 658,780 | 43,650 | |
| | 2 | 82.23 | 50,367,196 | 471,766 | | |
| | 3 | 78.98 | 45,806,634 | 404,110 | | |

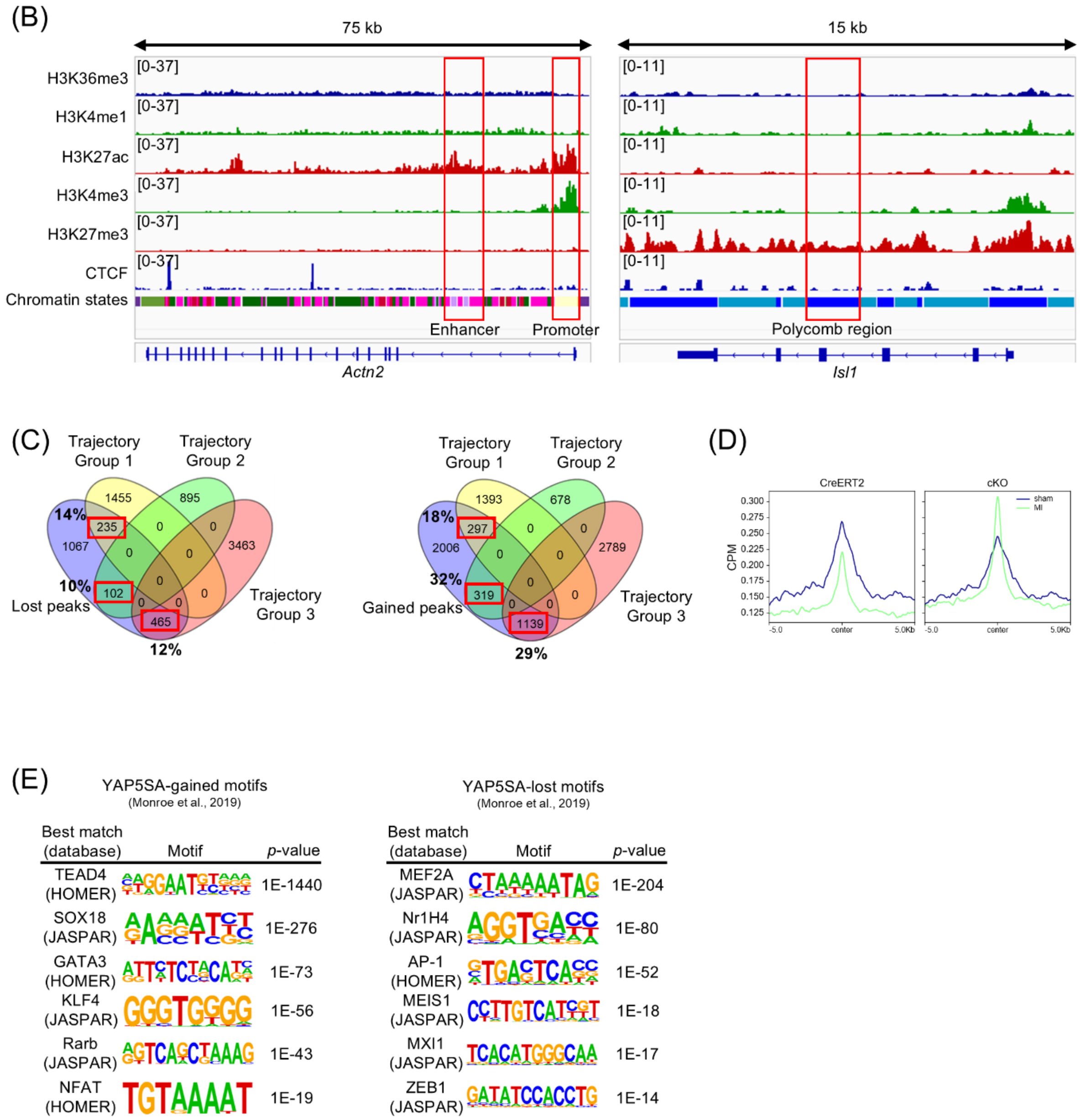

**Figure S6.**

Extended analyses supporting injury-associated chromatin accessibility remodeling in RNF20-deficient cardiomyocytes. Related to Figure 5.

(**A**) Quality of ATAC-seq.

(**B**) Representative genome tracks illustrating distributions of epigenetic marks and the corresponding chromatin state annotations. Red boxes highlight regulatory regions.

(**C**) Venn diagrams (*82*) showing overlap between pseudotime-dependent genes (Fig. 3B) and genes linked to differentially accessible peaks. Red boxes denote overlapping genes. Percentages indicate the proportion of pseudotime-dependent genes overlapping with differentially accessible regions.

(**D**) Metagene profile plots showing ATAC-seq signals centered on injury-associated peaks. Peak sets (4-day post-injury) were generated by van Duijvenboden et al. (*68*). CPM, counts per million; MI, myocardial infarction.

(**E**) De novo motif enrichment analysis performed with ATAC-seq from YAP5SA adult murine cardiomyocyte nuclei (*21*).

758

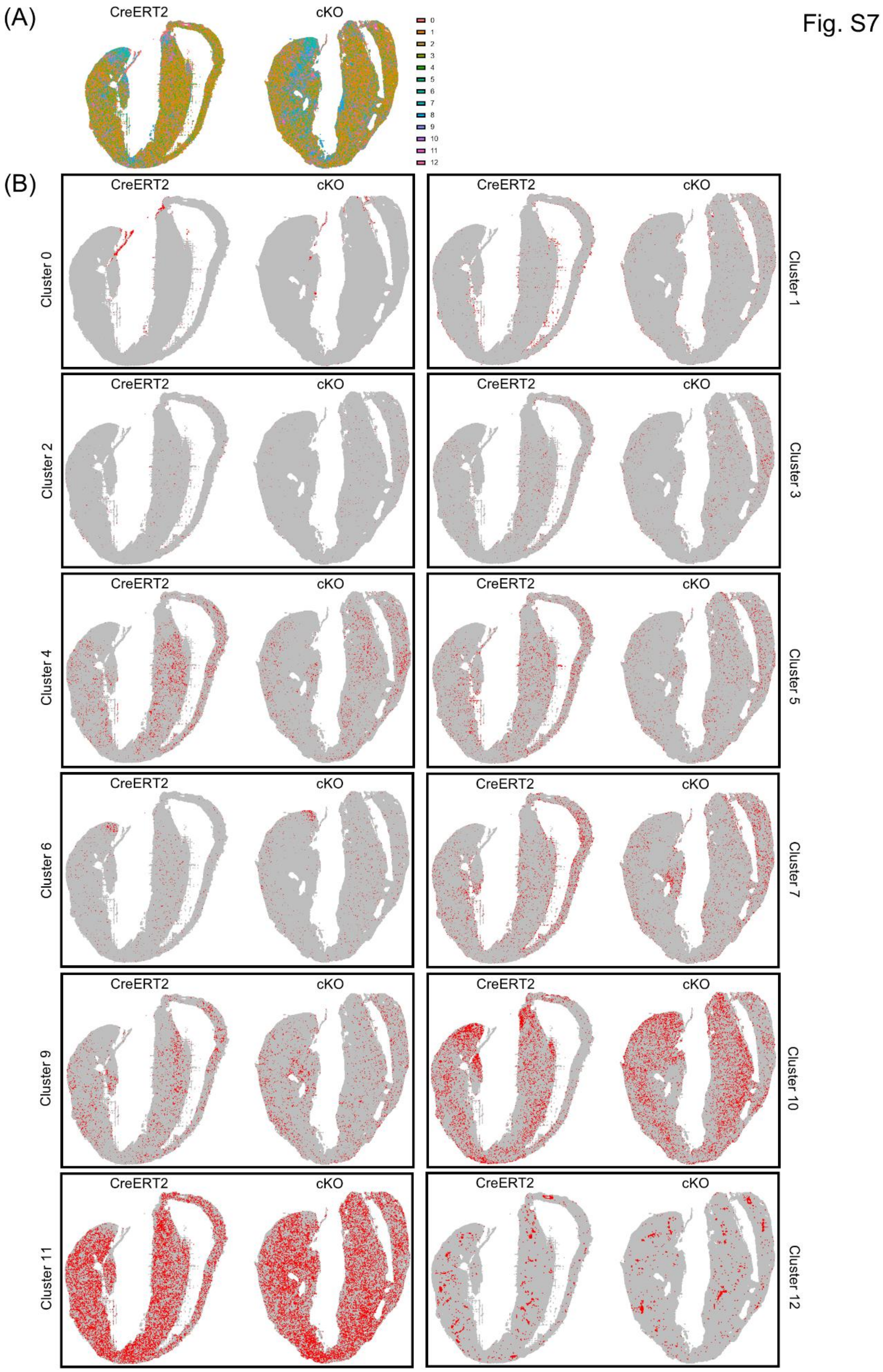

Fig. S7
(A)
CreERT2
cKO
0
1
2
3
4
5
6
7
8
9
10
11
12
(B)
Cluster 0
Cluster 1
Cluster 2
Cluster 3
Cluster 4
Cluster 5
Cluster 6
Cluster 7
Cluster 9
Cluster 10
Cluster 11
Cluster 12

**Figure S7.**

Spatial mapping of clusters across the myocardium. Related to Figure 6.

(**A**) Combined cluster maps.

(**B**) Cluster-specific maps. Positive bins are shown in red and negative bins in grey.

Fig. S8

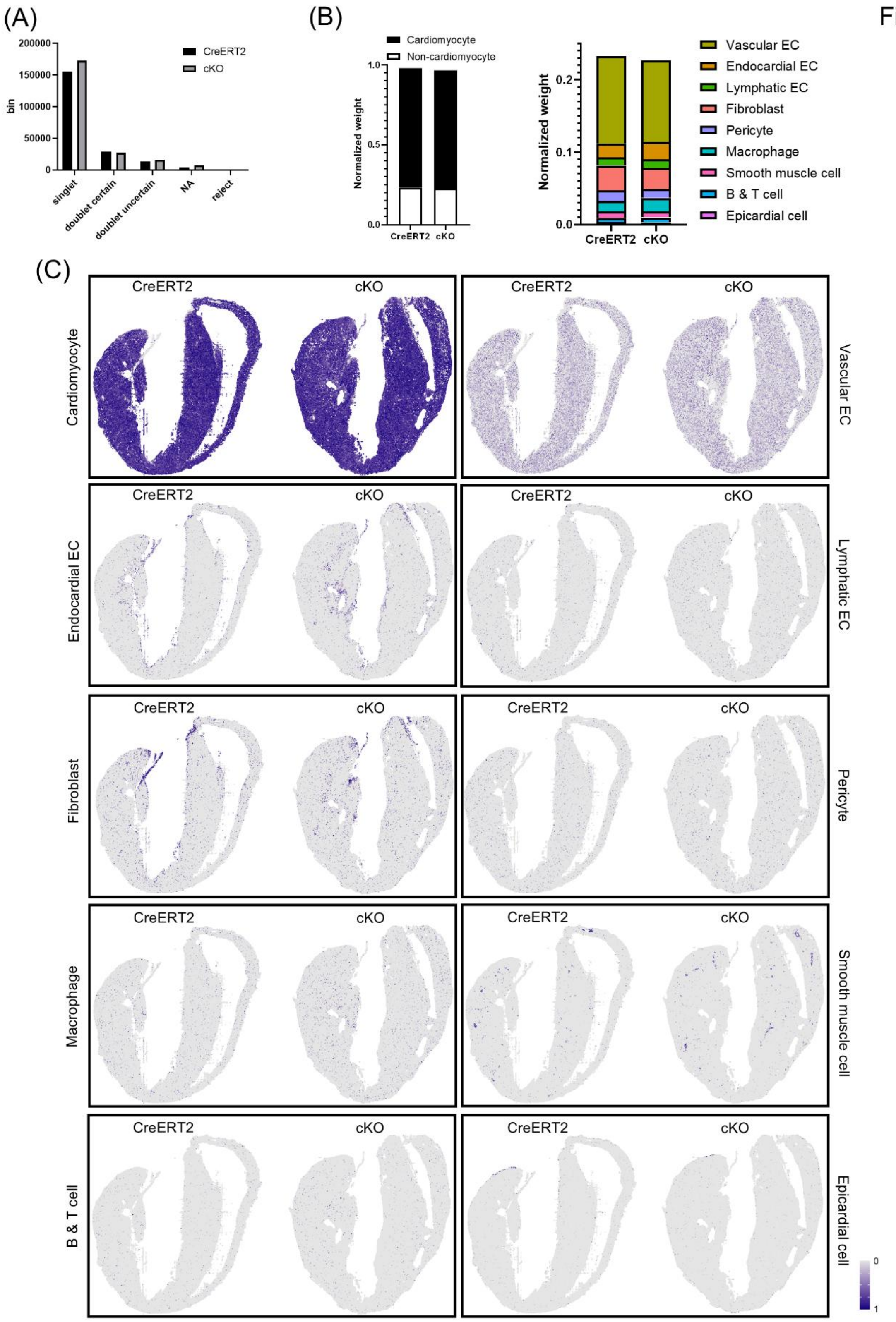
(A)
(B)
(C)
CreERT2
cKO
bin
200000
150000
100000
50000
0
singlet
doublet certain
doublet uncertain
NA
reject
Cardiomyocyte
Non-cardiomyocyte
Normalized weight
1.0
0.5
0.0
0.2
0.1
Vascular EC
Endocardial EC
Lymphatic EC
Fibroblast
Pericyte
Macrophage
Smooth muscle cell
B & T cell
Epicardial cell

764

**Figure S8.**

Results from spatial deconvolution. Related to Figure 6.

(**A**) Spot class assignments from RCTD deconvolution. NA, not available.

(**B**) Left: proportion of cell-type weights across genotypes. Right: proportion of non-cardiomyocyte cell-type weights across genotypes. EC, endothelial cell.

(**C**) Spatial feature plots showing weights of indicated cell types.

Fig. S9

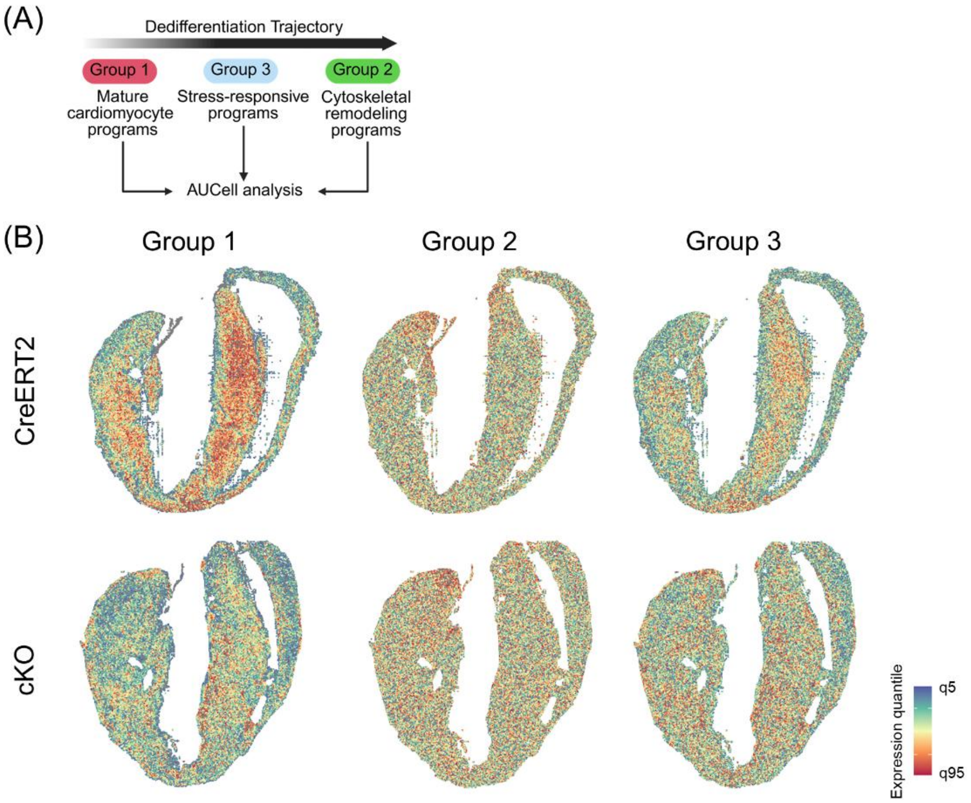


**Figure S9.**

Spatial gene signature scoring. Related to Figure 6.

(**A**) Schematic for AUCell analysis using pseudotime-dependent genes derived from snRNA-seq (Fig. 3B).

(**B**) Spatial feature plots showing AUCell scores for pseudotime-dependent genes.

Fig. S10

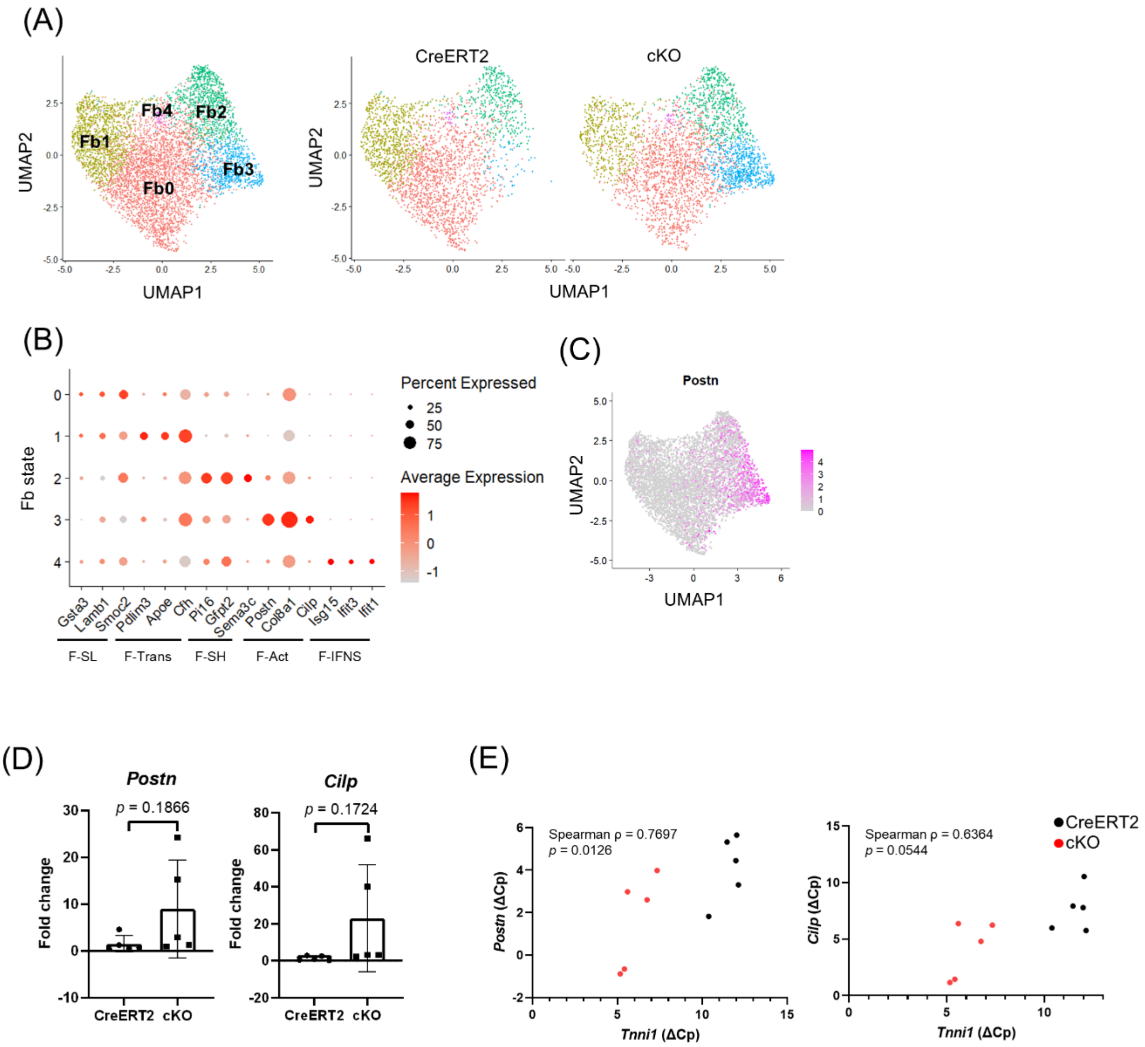


**Figure S10.**

Validation of fibroblast activation. Related to Figure 6.

(**A**) UMAP visualization of subclustered fibroblasts colored by cell states (left) and separated by genotype (right).

(**B**) Dot plot of expression of known marker genes among fibroblast states.(*83*) F-SL, *Sca1*$^{low}$ fibroblasts; F-Trans, transitory fibroblast; F-SH, *Sca1*$^{high}$ fibroblasts; F-Act, activated fibroblast; F-IFNS, interferon-stimulated fibroblast.

(**C**) *Postn* expression in fibroblasts.

(**D**) RT-qPCR analysis of *Postn* (left) and *Cilp* (right) from LV tissues at 2 wpt. 18S rRNA was used as normalization control. Data were generated with n = 5 biological replicates and are presented as mean ± s.d.. Unpaired two-tailed t test with Welch's correction.

(**E**) Correlation analysis between *Tnni1* and *Postn* (left) or *Tnni1* and *Cilp* (right) expression levels. Δ Cp values were determined by RT-qPCR analysis. Associations were assessed using Spearman's rank correlation.

Fig. S11

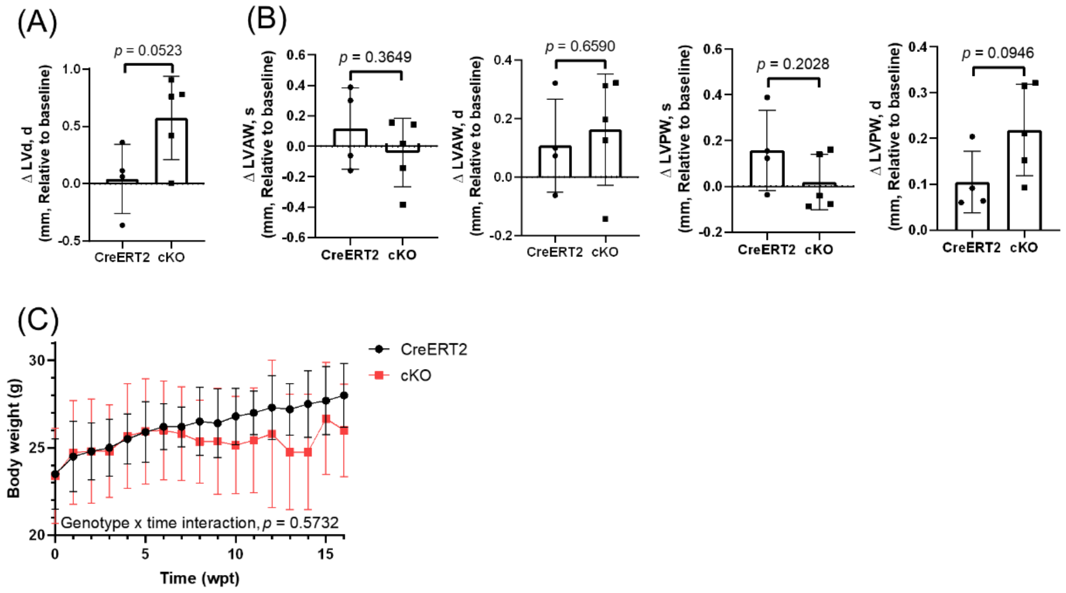


**Figure S11.**

Limited changes in LV wall thickness in the hearts from cKO mice. Related to Figure 7.

(**A**) Change in LV end-diastolic internal diameter (LVd, d) at 10 wpt relative to baseline. Data were generated with n = 4 (CreERT2) and 5 (cKO) biological replicates and are presented as mean ± s.d.. Unpaired two-tailed t test.

(**B**) Change in LV wall thickness at 10 wpt relative to baseline. LVAW, LV anterior wall; LVPW, LV posterior wall. Data were generated with n = 4 (CreERT2) and 5 (cKO) biological replicates and are presented as mean ± s.d.. Unpaired two-tailed t test.

(**C**) Body weight. Data were generated with n = 10 (CreERT2), 15 (cKO, 0-7 wpt), 14 (cKO, 8 wpt), 11 (cKO, 9 wpt), 7 (cKO, 10-11 wpt), 5 (cKO, 12 wpt), 4 (cKO, 13-14 wpt), and 3 (cKO, 15-16 wpt) biological replicates and are presented as mean ± s.d.. The genotype × time interaction was assessed using a mixed-effects model.

Fig. S12

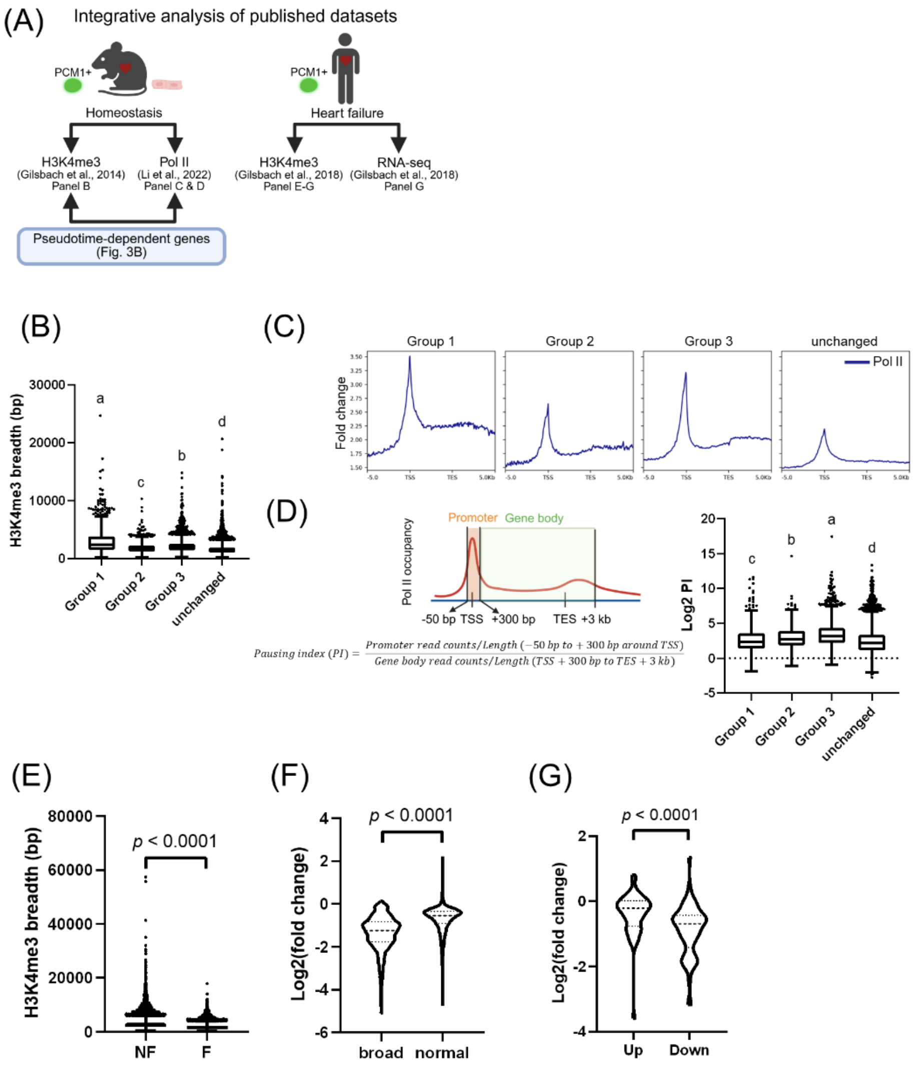


**Figure S12.**

RNF20-dependent mature cardiomyocyte transcriptional programs exhibit broader H3K4me3 domains and efficient transcriptional elongation.

(**A**) Schematic illustrating the integration of reanalyzed published epigenomic datasets with pseudotime-dependent genes identified in this study (Fig. 3B). Published H3K4me3 ChIP-seq (*66*) and RNA polymerase II (Pol II) ChIP-seq (*67*) from homeostatic adult murine cardiomyocytes were reanalyzed to evaluate H3K4me3 domain breadth and promoter-proximal pausing of pseudotime-dependent genes. Published human heart-failure cardiomyocytes H3K4me3 ChIP-seq and bulk RNA-seq were further analyzed to assess association between H3K4me3 breadth and gene expression under stress (*58*).

(**B**) H3K4me3 peak breadth among pseudotime-dependent genes. Data are presented as box-and-whisker plots. Center line indicates the median; box indicates the interquartile range (IQR); whiskers extend to the most extreme values within 1.5 × IQR; points represent outliers. Kruskal–Wallis test followed by Dunn's test. Groups with different lowercase letters are significantly different (adjusted *p*-value< 0.05).

(**C**) Metagene profile plots showing occupancy of Pol II across pseudotime-dependent genes in adult murine cardiomyocytes, spanning gene bodies with 5 kb upstream and downstream flanking regions. The y-axis represents fold change of signal relative to input. TSS, transcription start site; TES, transcription end site.

(**D**) Left: Schematic of pausing index (PI) calculation adapted from Day et al. (*80*) Right: PI among pseudotime-dependent genes in adult murine cardiomyocytes. Data are presented as box-and-whisker plots. Center line indicates the median; box indicates the IQR; whiskers extend to the most extreme values within 1.5 × IQR; points represent outliers. Kruskal–Wallis test followed by Dunn's test. Groups with different lowercase letters are significantly different (adjusted *p*-value< 0.05).

(**E**) H3K4me3 peak breadth was globally reduced in adult human failing cardiomyocytes (F) compared with non-failing cardiomyocytes (NF). Data are presented as box-and-whisker plots. Center line indicates the median; box indicates the IQR; whiskers extend to the most extreme values within 1.5 × IQR; points represent outliers. Mann-Whitney test.

(**F**) Narrowing of H3K4me3 peak breadth was more pronounced in broad domains (breadth> $95^{\text{th}}$ percentile) compared with normal domains (breadth≦ $95^{\text{th}}$ percentile). Data are presented as violin plots. Mann-Whitney test.

(**G**) Downregulated genes (Down) exhibited reduced H3K4me3 peak breadth, whereas upregulated genes (Up) were relatively stable in peak breadth. Data are presented as violin plots. Mann-Whitney test.

Fig. S13

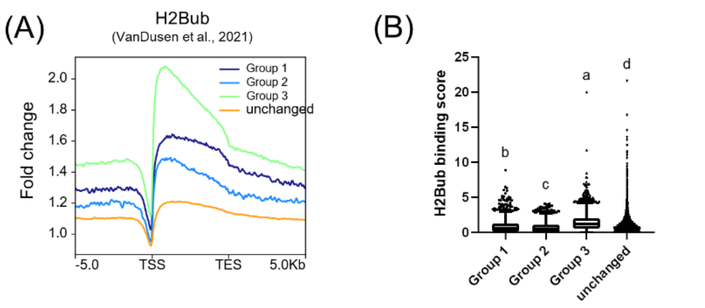


**Figure S13.**

Stress-responsive genes show higher gene-body H2Bub enrichment.

(**A**) Metagene profile plot showing occupancy of H2Bub across pseudotime-dependent genes (Fig. 3B) in postnatal day 28 murine hearts, spanning gene bodies with 5 kb upstream and downstream flanking regions. The y-axis represents fold change of signal relative to input.

(**B**) Gene length-normalized H2Bub binding scores among pseudotime-dependent genes. Normalized H2Bub binding scores were obtained from VanDusen et al. (*65*). Data were generated with n = 2 biological replicates and are presented as box-and-whisker plot. Center line indicates the median; box indicates the IQR range; whiskers extend to the most extreme values within 1.5 × IQR; points represent outliers. Kruskal–Wallis test followed by Dunn’s test. Groups with different lowercase letters are significantly different (adjusted $p$-value $< 0.05$).

Fig. S14

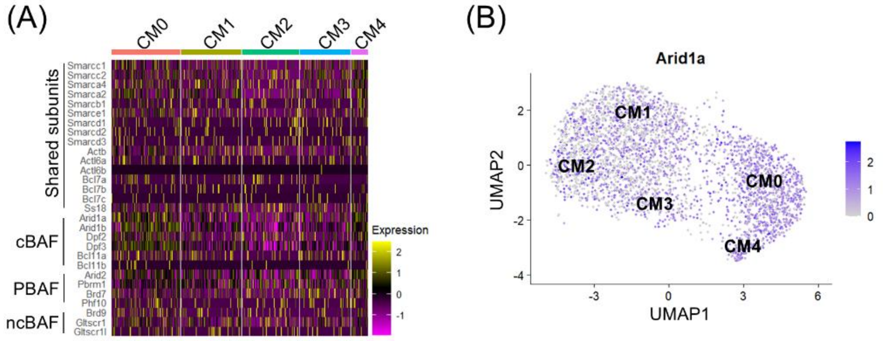


**Fig S14.**

Increased expression of SWI/SNF complex genes in cKO cardiomyocytes.

(**A**) Heatmap showing expression of SWI/SNF complex subunits among cardiomyocyte states. cBAF, canonical BAF; PBAF, polybromo-associated BAF; ncBAF, non-canonical BAF (*84*).

(**B**) *Arid1a* expression in cardiomyocytes.

| Application | Target | Forward (5’→3’) | Reverse (5’→3’) |
|---|---|---|---|
| Genotyping | *Rnf20*$^{flox/flox}$ | CTTGAGAGTGCTATACTTGCTG | GATTCTTGGATTGCAGTGTTCG |
| | *Cre* | CCACGACCAAGTGACAGCAATG | CAGAGACGGAAATCCATCGCTC |
| RT-qPCR | *Myh6* | GAGTGCTTCGTGCCTGATGAC | TCCTTTATGGTCACCGTCTTTCC |
| | *Myh7* | GTGCCAAGGGCCTGAATGAG | GCAAAGGCTCCAGGTCTGA |
| | *Tnni1* | GGCTCTAAGCACAAGGTGTCCA | GCCAGACATAGCCTCCACATTC |
| | *Tnni3* | CAACTACCGAGCCTATGCCAC | GCAATCTGCAGCATCAGAGTCTTC |
| | *Gapdh* | AGCTCACTGGCATGGCCTTC | ATGCCTGCTTCACCACCTTC |
| | *Rnf20* | CGAGTGTCCGTCTTGGAGTC | GCCTTTTGAATTCACCCGCTC |
| | 18S rRNA | TCGTCTTCGAAACTCCGACT | CGCGGTTCTATTTGTTGGT |
| | *Postn* | ATCAAGGTGCTATCTGCGGG | GTCAATAGGCATCACTGCGG |
| | *Cilp* | TCACGATGCCCAAGACTAGC | ACAATGTATGGGGTCTCTGCC |

**Table S1.**

Primers used in genotyping and RT-qPCR.

| REAGENT or RESOURCE | SOURCE | IDENTIFIER |
|---|---|---|
| Antibodies | | |
| Rabbit polyclonal anti-RNF20 antibody | Proteintech | Cat#21625-1-AP; RRID: AB_10734436 |
| Mouse monoclonal anti-cTnT antibody | Thermo Fisher | Cat#MA5-12960; RRID: AB_11000742 |
| Rabbit polyclonal anti-Cx43 antibody | Abcam | Cat#ab11370; RRID: AB_297976 |
| Rabbit recombinant monoclonal anti-YAP1 antibody (D8H1X) | Cell Signaling | Cat#14074; RRID: AB_2650491 |
| Rabbit recombinant monoclonal anti-p-YAP1 (S127) antibody (D9W2I) | Cell Signaling | Cat#13008; RRID: AB_2650553 |
| Rabbit polyclonal anti-GAPDH antibody | GeneTex | Cat#GTX100118; RRID: AB_1080976 |
| Rabbit monoclonal anti-a-actinin antibody | Abcam | Cat#ab68167; RRID: AB_11157538 |
| Mouse monoclonal-a-SMA antibody | Sigma-Aldrich | Cat#C6198; RRID: AB_476856 |
| Rabbit polyclonal anti-dystrophin antibody | Abcam | Cat#ab15277; RRID: AB_301813 |
| Mouse monoclonal anti-N-cadherin antibody | Thermo Fisher | Cat#33-3900; RRID: AB_2313779 |
| Rabbit polyclonal anti-Ki67 antibody | Abcam | Cat#ab15580; RRID: AB_443209 |
| Mouse monoclonal anti-PCM1-Alexa Fluor® 647 antibody | Santa Cruz | Cat#SC-398365; RRID: AB_2827155 |
| Mouse monoclonal anti-a-tubulin antibody | Cell Signaling | Cat#3873; RRID: AB_1904178 |
| Rabbit polyclonal anti-detyrosinated a-tubulin antibody | Abcam | Cat#ab48389; RRID: AB_869990 |
| Chemicals, peptides, and recombinant proteins | | |
| Ex Taq DNA polymerase | TaKaRa | Cat#RR001 |
| Tamoxifen | Sigma-Aldrich | Cat#T5648 |
| EdU | Sigma-Aldrich | Cat#900584 |

| | | |
|---|---|---|
| Protease inhibitor cocktail | Roche | Cat#4693132001 |
| Phosphatase inhibitor cocktail | Sigma-Aldrich | Cat#P0044 |
| TRIzol™ reagent | Thermo Fisher | Cat#15596026 |
| Collagenase B | Roche | Cat#111088807001 |
| Collagenase D | Roche | Cat#11088858001 |
| Protease XIV | Sigma-Aldrich | Cat#P5147 |
| Critical commercial assays | | |
| QuantiTect Reverse Transcription Kit | QIAGEN | Cat#205311 |
| KAPA SYBR® FAST qPCR Master Mix | Kapa Biosystems | Cat#KR0389_S |
| Click-iT Plus EdU Cell Proliferation Kit | Thermo Fisher | Cat#C10640 |
| TUNEL assay Kit | Elabscience | Cat#E-CK-A324 |
| Chromium Next GEM Single Cell 3′ Kit v3.1 Chemistry | 10x Genomics | Cat#PN-1000269 |
| Illumina Tagment DNA TDE1 Enzyme and Buffer Kits | Illumina | Cat#20034210 |
| MinElute PCR Purification Kit | QIAGEN | Cat#28004 |
| Visium HD Spatial Gene Expression Reagent Kit | 10x Genomics | Cat#PN-1000668 |
| Visium Mouse Transcriptome Probe Set v2.0 | 10x Genomics | Cat#PN-1000667 |
| Deposited data | | |
| SnRNA-seq | This study | BioProject: PRJNA1496066 |
| Spatial transcriptomics | This study | BioProject: PRJNA1496066 |
| ATAC-seq | This study | BioProject: PRJNA1496066 |
| Uncropped Western blots | This study | https://doi.org/10.5281/zenodo.21700467 |
| Murine cardiomyocyte bulk RNA-seq | Feng and Nie (*57*) | BioProject: PRJNA798841 |
| Murine heart H2Bub ChIP-seq | VanDusen et al. (*65*) | GEO: GSE139975 |
| Murine myocardial infarction snRNA-seq | Cui et al. (*20*) | GEO: GSE130699 |
| Murine cardiomyocyte nuclei H3K4me3 and H3K27ac ChIP-seq | Gilsbach et al. (*66*) | BioProject: PRJNA229480 |
| Murine YAP5SA cardiomyocyte nuclei ATAC-seq | Monroe et al. (*21*) | GEO: GSE123457 |
| Murine cardiomyocyte Pol II ChIP-seq | Li et al. (*67*) | BioProject: PRJNA655138 |
| Human heart failure cardiomyocyte nuclei RNA-seq and H3K4me3 ChIP-seq | Gilsbach et al. (*58*) | BioProject: PRJNA353755 |

| | | |
|---|---|---|
| Murine heart H3K4me3 ChIP-seq | ENCODE | ENCODE: ENCSR000CAM |
| Murine heart H3K27me3 ChIP-seq | ENCODE | ENCODE: ENCSR000CEJ |
| Murine heart H3K27ac ChIP-seq | ENCODE | ENCODE: ENCSR000CDF |
| Murine heart H3K4me1 ChIP-seq | ENCODE | ENCODE: ENCSR000CAE |
| Murine heart H3K36me3 ChIP-seq | ENCODE | ENCODE: ENCSR000CEK |
| Murine heart CTCF ChIP-seq | ENCODE | ENCODE: ENCSR000CBI |
| Murine heart input | ENCODE | ENCODE: ENCSR000CAV |
| Experimental models: Organisms/strains | | |
| Mouse: C57BL/6JNarl-*Rnf20*$^{flox/flox}$ | Lin et al. (*1*) | N/A |
| Mouse: B6.FVB(129)-A1cfTg(Myh6-cre/Esr1*)1Jmk/J | The Jackson Laboratory | RRID:IMSR_JAX:005657 |
| Mouse: C57BL/6JNarl; *Rnf20*$^{flox/flox}$; *aMHC-MerCreMer*$^{+/-}$ | This study | N/A |
| Oligonucleotides | | |
| See Table S1 | | |
| Software and algorithms | | |
| Fiji | National Institute of Health | v1.54 |
| Image Lab software | Bio-Rad | v6.1 |
| SOTAtool | Stein et al. (*75*) | https://github.com/steinjm/SotaTool |
| Galaxy-Main | The Galaxy Community (*11*) | https://usegalaxy.org/ |
| Galaxy-Europe | The Galaxy Community (*11*) | https://usegalaxy.eu/ |
| STAR | Kaminow et al. (*9*); Dobin et al. (*10*) | V2.7.11a |

| | | |
|---|---|---|
| Trailmaker™ | Parse Biosciences | https://app.trailmaker.parsebiosciences.com/ |
| Seurat | Hao et al. (*14*) | v4; v5.5.1 |
| scDblFinder | Germain et al. (*13*) | v1.22.0 |
| CytoTRACE | Gulati et al. (*15*) | v0.3.3 |
| AUCell | Aibar et al. (*16*) | v1.30.1 |
| Monocle3 | Cao et al. (*17*) | v1.4.26 |
| NOIseq | Tarazona et al. (*62*) | v2.48.0 |
| masigPro | Conesa at al. (*64*) | v1.76.0 |
| RUVseq | Risso et al. (*38*) | v1.42.0 |
| DESeq2 | Love et al. (*39*) | v1.48.2 |
| ChIPSeeker | Yu et al. (*40*) | v1.44.0 |
| ChromHMM | Ernst et al. (*41*) | v1.27 |
| rGREAT | Gu et al. (*50*) | v2.10.0 |
| GOSemSim | Yu et al. (*52*) | v2.34.0 |
| Cytoscape | Shannon et al. (*53*) | v3.10.4 |
| HOMER | Heinz et al. (*54*) | v5.1 |
| TOBIAS | Bentsen et al. (*55*) | v0.14.0 |
| FastQC | Andrews et al. (*26*) | v0.12.1 |
| Cutadapt | Martin et al. (*27*) | v5.2 |
| Bowtie2 | Langmead et al. (*29*) | v2.5.5 |
| BAMTools | Barnett et al. (*30*) | v2.5.3 |
| Picard | Broad Institute (*31*) | v3.1.1 |
| deepTools | Ramirez et al. (*33*) | v3.5.4 |
| BEDTools | Quinlan et al. (*34*) | v2.31.1 |
| MACS2 | Zhang et al. (*35*) | v2.2.91 |
| UCSC Genome Browser | Hinrichs et al. (*69*) | RRID:SCR_005780 |
| Integrative Genomics Viewers | Robinson et al. (*70*) | v2.19.7 |
| subread | Liao et al. (*61*); Liao et al. (*60*) | v2.0.3 |
| Space Ranger count | 10x Genomics | v4.0.1 |
| Loupe browser | 10x Genomics | v9 |
| spacexr | Cable et al. (*71*) | v1.4.0 |

| | | |
|---|---|---|
| Pausing Index Calculator | Day et al. (*80*) | https://github.com/MiMiroot/PIC |
| DAVID | Huang da et al. (*22*) | v2023q4 |
| Iterative Overlap Peak Merging | Corces et al. (*37*) | https://github.com/corceslab/ATAC_IterativeOverlapPeakMerging |
| Customized codes for distance-based analysis | This study | https://doi.org/10.5281/zenodo.21700369 |
| GraphPad PRISM | GraphPad | RRID:SCR_002798; v8 |
| R Project for Statistical Computing | N/A | RRID:SCR_001905; v4.5.0 |

**Table S2.**

Key resources table.

**Data S1. (separate file)**

Maturation-associated gene sets.

**Data S2. (separate file)**

The complete lists of enriched and the top 100 differential terms from the rGREAT analysis.

### REFERENCES

1. C.-Y. Lin, Y.-M. Chang, H.-Y. Tseng, Y.-L. Shih, H.-H. Yeh, Y.-R. Liao, H.-H. Tang, C.-L. Hsu, C.-C. Chen, Y.-T. Yan, C.-F. Kao, Epigenetic regulator RNF20 underlies temporal hierarchy of gene expression to regulate postnatal cardiomyocyte polarization. *Cell Reports* **42**, 113416 (2023).
2. D. S. Sohal, M. Nghiem, M. A. Crackower, S. A. Witt, T. R. Kimball, K. M. Tymitz, J. M. Penninger, J. D. Molkentin, Temporally regulated and tissue-specific gene manipulations in the adult and embryonic heart using a tamoxifen-inducible Cre protein. *Circ Res* **89**, 20–25 (2001).
3. G. E. Truett, P. Heeger, R. L. Mynatt, A. A. Truett, J. A. Walker, M. L. Warman, Preparation of PCR-quality mouse genomic DNA with hot sodium hydroxide and tris (HotSHOT). *Biotechniques* **29**, 52, 54 (2000).
4. M. Ackers-Johnson, P. Y. Li, A. P. Holmes, S. M. O'Brien, D. Pavlovic, R. S. Foo, A Simplified, Langendorff-Free Method for Concomitant Isolation of Viable Cardiac Myocytes and Nonmyocytes From the Adult Mouse Heart. *Circ Res* **119**, 909–920 (2016).
5. M. D. Santos, S. Gioftsidi, S. Backer, L. Machado, F. Relaix, P. Maire, P. Mourikis, Extraction and sequencing of single nuclei from murine skeletal muscles. *STAR Protoc* **2**, 100694 (2021).
6. M. Yekelchyk, X. Li, S. Guenther, T. Braun, Single-Nucleus ATAC-seq for Mapping Chromatin Accessibility in Individual Cells of Murine Hearts. *Methods Mol Biol* **2752**, 245–257 (2024).
7. J. D. Buenrostro, B. Wu, H. Y. Chang, W. J. Greenleaf, ATAC-seq: A Method for Assaying Chromatin Accessibility Genome-Wide. *Curr Protoc Mol Biol* **109**, 21 29 21–21 29 29 (2015).
8. M. R. Corces, A. E. Trevino, E. G. Hamilton, P. G. Greenside, N. A. Sinnott-Armstrong, S. Vesuna, A. T. Satpathy, A. J. Rubin, K. S. Montine, B. Wu, A. Kathiria, S. W. Cho, M. R. Mumbach, A. C. Carter, M. Kasowski, L. A. Orloff, V. I. Risca, A. Kundaje, P. A. Khavari, T. J. Montine, W. J. Greenleaf, H. Y. Chang, An improved ATAC-seq protocol reduces background and enables interrogation of frozen tissues. *Nat Methods* **14**, 959–962 (2017).
9. B. Kaminow, D. Yunusov, A. Dobin, STARsolo: accurate, fast and versatile mapping/quantification of single-cell and single-nucleus RNA-seq data. *bioRxiv*, (2021).
10. A. Dobin, C. A. Davis, F. Schlesinger, J. Drenkow, C. Zaleski, S. Jha, P. Batut, M. Chaisson, T. R. Gingeras, STAR: ultrafast universal RNA-seq

aligner. *Bioinformatics* **29**, 15–21 (2013).
11. C. Galaxy, Galaxy for accessible, reproducible, and collaborative data analyses: 2026 update. *Nucleic Acids Res* **54**, W105–W116 (2026).
12. A. T. L. Lun, S. Riesenfeld, T. Andrews, T. P. Dao, T. Gomes, J. participants in the 1st Human Cell Atlas, J. C. Marioni, EmptyDrops: distinguishing cells from empty droplets in droplet-based single-cell RNA sequencing data. *Genome Biol* **20**, 63 (2019).
13. P. L. Germain, A. Lun, C. Garcia Meixide, W. Macnair, M. D. Robinson, Doublet identification in single-cell sequencing data using scDblFinder. *F1000Res* **10**, 979 (2021).
14. Y. Hao, T. Stuart, M. H. Kowalski, S. Choudhary, P. Hoffman, A. Hartman, A. Srivastava, G. Molla, S. Madad, C. Fernandez-Granda, R. Satija, Dictionary learning for integrative, multimodal and scalable single-cell analysis. *Nat Biotechnol* **42**, 293–304 (2024).
15. G. S. Gulati, S. S. Sikandar, D. J. Wesche, A. Manjunath, A. Bharadwaj, M. J. Berger, F. Ilagan, A. H. Kuo, R. W. Hsieh, S. Cai, M. Zabala, F. A. Scheeren, N. A. Lobo, D. Qian, F. B. Yu, F. M. Dirbas, M. F. Clarke, A. M. Newman, Single-cell transcriptional diversity is a hallmark of developmental potential. *Science* **367**, 405–411 (2020).
16. S. Aibar, S. Aerts, AUCell: analysis of'gene set'activity in single-cell RNA-seq data. *R/Bioconductor package*,  (2016).
17. J. Cao, M. Spielmann, X. Qiu, X. Huang, D. M. Ibrahim, A. J. Hill, F. Zhang, S. Mundlos, L. Christiansen, F. J. Steemers, C. Trapnell, J. Shendure, The single-cell transcriptional landscape of mammalian organogenesis. *Nature* **566**, 496–502 (2019).
18. Z. Gu, R. Eils, M. Schlesner, Complex heatmaps reveal patterns and correlations in multidimensional genomic data. *Bioinformatics* **32**, 2847–2849 (2016).
19. Z. Gu, Complex heatmap visualization. *Imeta* **1**, e43 (2022).
20. M. Cui, Z. Wang, K. Chen, A. M. Shah, W. Tan, L. Duan, E. Sanchez-Ortiz, H. Li, L. Xu, N. Liu, R. Bassel-Duby, E. N. Olson, Dynamic Transcriptional Responses to Injury of Regenerative and Non-regenerative Cardiomyocytes Revealed by Single-Nucleus RNA Sequencing. *Dev Cell* **55**, 665–667 (2020).
21. T. O. Monroe, M. C. Hill, Y. Morikawa, J. P. Leach, T. Heallen, S. Cao, P. H. L. Krijger, W. de Laat, X. H. T. Wehrens, G. G. Rodney, J. F. Martin, YAP Partially Reprograms Chromatin Accessibility to Directly Induce Adult

Cardiogenesis In Vivo. *Dev Cell* **48**, 765–779 e767 (2019).

22. W. Huang da, B. T. Sherman, R. A. Lempicki, Systematic and integrative analysis of large gene lists using DAVID bioinformatics resources. *Nat Protoc* **4**, 44–57 (2009).
23. B. T. Sherman, M. Hao, J. Qiu, X. Jiao, M. W. Baseler, H. C. Lane, T. Imamichi, W. Chang, DAVID: a web server for functional enrichment analysis and functional annotation of gene lists (2021 update). *Nucleic Acids Res* **50**, W216–W221 (2022).
24. L. Delisle, M. Doyle, F. Heyl.
25. S. Hiltemann, H. Rasche, S. Gladman, H.-R. Hotz, D. Larivière, D. Blankenberg, P. D. Jagtap, T. Wollmann, A. Bretaudeau, N. Goué, T. J. Griffin, C. Royaux, Y. L. Bras, S. Mehta, A. Syme, F. Coppens, B. Droesbeke, N. Soranzo, W. Bacon, F. Psomopoulos, C. Gallardo-Alba, J. Davis, M. C. Föll, M. Fahrner, M. A. Doyle, B. Serrano-Solano, A. C. Fouilloux, P. Heusden, W. Maier, D. Clements, F. Heyl, B. Grüning, B. B. and, Galaxy Training: A powerful framework for teaching! *PLoS Comput Biol* **19**, e1010752 (2023).
26. S. Andrews. (Babraham Institute, Babraham, UK, 2010).
27. M. Martin, Cutadapt removes adapter sequences from high-throughput sequencing reads. *EMBnet. journal* **17**, 10–12 (2011).
28. B. Langmead, C. Trapnell, M. Pop, S. L. Salzberg, Ultrafast and memory-efficient alignment of short DNA sequences to the human genome. *Genome Biol* **10**, R25 (2009).
29. B. Langmead, S. L. Salzberg, Fast gapped-read alignment with Bowtie 2. *Nat Methods* **9**, 357–359 (2012).
30. D. W. Barnett, E. K. Garrison, A. R. Quinlan, M. P. Stromberg, G. T. Marth, BamTools: a C++ API and toolkit for analyzing and managing BAM files. *Bioinformatics* **27**, 1691–1692 (2011).
31. in *Broad Institute, GitHub repository*. (Broad Institute, 2019).
32. H. M. Amemiya, A. Kundaje, A. P. Boyle, The ENCODE Blacklist: Identification of Problematic Regions of the Genome. *Sci Rep* **9**, 9354 (2019).
33. F. Ramirez, D. P. Ryan, B. Gruning, V. Bhardwaj, F. Kilpert, A. S. Richter, S. Heyne, F. Dundar, T. Manke, deepTools2: a next generation web server for deep-sequencing data analysis. *Nucleic Acids Res* **44**, W160–165 (2016).
34. A. R. Quinlan, I. M. Hall, BEDTools: a flexible suite of utilities for comparing genomic features. *Bioinformatics* **26**, 841–842 (2010).

35. Y. Zhang, T. Liu, C. A. Meyer, J. Eeckhoute, D. S. Johnson, B. E. Bernstein, C. Nusbaum, R. M. Myers, M. Brown, W. Li, X. S. Liu, Model-based analysis of ChIP-Seq (MACS). *Genome Biol* **9**, R137 (2008).

36. J. Feng, T. Liu, B. Qin, Y. Zhang, X. S. Liu, Identifying ChIP-seq enrichment using MACS. *Nat Protoc* **7**, 1728–1740 (2012).

37. M. R. Corces, J. M. Granja, S. Shams, B. H. Louie, J. A. Seoane, W. Zhou, T. C. Silva, C. Groeneveld, C. K. Wong, S. W. Cho, A. T. Satpathy, M. R. Mumbach, K. A. Hoadley, A. G. Robertson, N. C. Sheffield, I. Felau, M. A. A. Castro, B. P. Berman, L. M. Staudt, J. C. Zenklusen, P. W. Laird, C. Curtis, N. Cancer Genome Atlas Analysis, W. J. Greenleaf, H. Y. Chang, The chromatin accessibility landscape of primary human cancers. *Science* **362**, (2018).

38. D. Risso, J. Ngai, T. P. Speed, S. Dudoit, Normalization of RNA-seq data using factor analysis of control genes or samples. *Nat Biotechnol* **32**, 896–902 (2014).

39. M. I. Love, W. Huber, S. Anders, Moderated estimation of fold change and dispersion for RNA-seq data with DESeq2. *Genome Biol* **15**, 550 (2014).

40. G. Yu, L. G. Wang, Q. Y. He, ChIPseeker: an R/Bioconductor package for ChIP peak annotation, comparison and visualization. *Bioinformatics* **31**, 2382–2383 (2015).

41. J. Ernst, M. Kellis, Chromatin-state discovery and genome annotation with ChromHMM. *Nat Protoc* **12**, 2478–2492 (2017).

42. E. P. Consortium, An integrated encyclopedia of DNA elements in the human genome. *Nature* **489**, 57–74 (2012).

43. B. C. Hitz, J.-W. Lee, O. Jolanki, M. S. Kagda, K. Graham, P. Sud, I. Gabdank, J. Seth Strattan, C. A. Sloan, T. Dreszer, L. D. Rowe, N. R. Podduturi, V. S. Malladi, E. T. Chan, J. M. Davidson, M. Ho, S. Miyasato, M. Simison, F. Tanaka, Y. Luo, I. Whaling, E. L. Hong, B. T. Lee, R. Sandstrom, E. Rynes, J. Nelson, A. Nishida, A. Ingersoll, M. Buckley, M. Frerker, D. S. Kim, N. Boley, D. Trout, A. Dobin, S. Rahmanian, D. Wyman, G. Balderrama-Gutierrez, F. Reese, N. C. Durand, O. Dudchenko, D. Weisz, S. S. P. Rao, A. Blackburn, D. Gkountaroulis, M. Sadr, M. Olshansky, Y. Eliaz, D. Nguyen, I. Bochkov, M. S. Shamim, R. Mahajan, E. Aiden, T. Gingeras, S. Heath, M. Hirst, W. James Kent, A. Kundaje, A. Mortazavi, B. Wold, J. M. Cherry, The ENCODE Uniform Analysis Pipelines. *bioRxiv*, 2023.2004.2004.535623 (2023).

44. Y. Luo, B. C. Hitz, I. Gabdank, J. A. Hilton, M. S. Kagda, B. Lam, Z. Myers, P.

Sud, J. Jou, K. Lin, U. K. Baymuradov, K. Graham, C. Litton, S. R. Miyasato, J. S. Strattan, O. Jolanki, J. W. Lee, F. Y. Tanaka, P. Adenekan, E. O'Neill, J. M. Cherry, New developments on the Encyclopedia of DNA Elements (ENCODE) data portal. *Nucleic Acids Res* **48**, D882–D889 (2020).

45. J. A. Simon, R. E. Kingston, Occupying chromatin: Polycomb mechanisms for getting to genomic targets, stopping transcriptional traffic, and staying put. *Mol Cell* **49**, 808–824 (2013).

46. A. Shilatifard, Chromatin modifications by methylation and ubiquitination: implications in the regulation of gene expression. *Annu Rev Biochem* **75**, 243–269 (2006).

47. M. P. Creyghton, A. W. Cheng, G. G. Welstead, T. Kooistra, B. W. Carey, E. J. Steine, J. Hanna, M. A. Lodato, G. M. Frampton, P. A. Sharp, L. A. Boyer, R. A. Young, R. Jaenisch, Histone H3K27ac separates active from poised enhancers and predicts developmental state. *Proc Natl Acad Sci U S A* **107**, 21931–21936 (2010).

48. M. G. Guenther, S. S. Levine, L. A. Boyer, R. Jaenisch, R. A. Young, A chromatin landmark and transcription initiation at most promoters in human cells. *Cell* **130**, 77–88 (2007).

49. S. Cuddapah, R. Jothi, D. E. Schones, T. Y. Roh, K. Cui, K. Zhao, Global analysis of the insulator binding protein CTCF in chromatin barrier regions reveals demarcation of active and repressive domains. *Genome Res* **19**, 24–32 (2009).

50. Z. Gu, D. Hubschmann, rGREAT: an R/bioconductor package for functional enrichment on genomic regions. *Bioinformatics* **39**, (2023).

51. C. Y. McLean, D. Bristor, M. Hiller, S. L. Clarke, B. T. Schaar, C. B. Lowe, A. M. Wenger, G. Bejerano, GREAT improves functional interpretation of cis-regulatory regions. *Nat Biotechnol* **28**, 495–501 (2010).

52. G. Yu, F. Li, Y. Qin, X. Bo, Y. Wu, S. Wang, GOSemSim: an R package for measuring semantic similarity among GO terms and gene products. *Bioinformatics* **26**, 976–978 (2010).

53. P. Shannon, A. Markiel, O. Ozier, N. S. Baliga, J. T. Wang, D. Ramage, N. Amin, B. Schwikowski, T. Ideker, Cytoscape: a software environment for integrated models of biomolecular interaction networks. *Genome Res* **13**, 2498–2504 (2003).

54. S. Heinz, C. Benner, N. Spann, E. Bertolino, Y. C. Lin, P. Laslo, J. X. Cheng, C. Murre, H. Singh, C. K. Glass, Simple combinations of lineage-determining transcription factors prime cis-regulatory elements required

for macrophage and B cell identities. *Mol Cell* **38**, 576–589 (2010).

55. M. Bentsen, P. Goymann, H. Schultheis, K. Klee, A. Petrova, R. Wiegandt, A. Fust, J. Preussner, C. Kuenne, T. Braun, J. Kim, M. Looso, ATAC-seq footprinting unravels kinetics of transcription factor binding during zygotic genome activation. *Nat Commun* **11**, 4267 (2020).

56. I. Rauluseviciute, R. Riudavets-Puig, R. Blanc-Mathieu, J. A. Castro-Mondragon, K. Ferenc, V. Kumar, R. B. Lemma, J. Lucas, J. Cheneby, D. Baranasic, A. Khan, O. Fornes, S. Gundersen, M. Johansen, E. Hovig, B. Lenhard, A. Sandelin, W. W. Wasserman, F. Parcy, A. Mathelier, JASPAR 2024: 20th anniversary of the open-access database of transcription factor binding profiles. *Nucleic Acids Res* **52**, D174–D182 (2024).

57. J. Feng, Y. Li, Y. Nie, Methods of mouse cardiomyocyte isolation from postnatal heart. *J Mol Cell Cardiol* **168**, 35–43 (2022).

58. R. Gilsbach, M. Schwaderer, S. Preissl, B. A. Gruning, D. Kranzhofer, P. Schneider, T. G. Nuhrenberg, S. Mulero-Navarro, D. Weichenhan, C. Braun, M. Dressen, A. R. Jacobs, H. Lahm, T. Doenst, R. Backofen, M. Krane, B. D. Gelb, L. Hein, Distinct epigenetic programs regulate cardiac myocyte development and disease in the human heart in vivo. *Nat Commun* **9**, 391 (2018).

59. B. Batut, M. Freeberg, M. Heydarian, A. Erxleben, P. Videm, C. Blank, M. Doyle, N. Soranzo, P. Heusden, L. Delisle.

60. Y. Liao, G. K. Smyth, W. Shi, featureCounts: an efficient general purpose program for assigning sequence reads to genomic features. *Bioinformatics* **30**, 923–930 (2014).

61. Y. Liao, G. K. Smyth, W. Shi, The Subread aligner: fast, accurate and scalable read mapping by seed-and-vote. *Nucleic Acids Res* **41**, e108 (2013).

62. S. Tarazona, F. Garcia-Alcalde, J. Dopazo, A. Ferrer, A. Conesa, Differential expression in RNA-seq: a matter of depth. *Genome Res* **21**, 2213–2223 (2011).

63. S. Tarazona, P. Furio-Tari, D. Turra, A. D. Pietro, M. J. Nueda, A. Ferrer, A. Conesa, Data quality aware analysis of differential expression in RNA-seq with NOISeq R/Bioc package. *Nucleic Acids Res* **43**, e140 (2015).

64. A. Conesa, M. J. Nueda, A. Ferrer, M. Talon, maSigPro: a method to identify significantly differential expression profiles in time-course microarray experiments. *Bioinformatics* **22**, 1096–1102 (2006).

65. N. J. VanDusen, J. Y. Lee, W. Gu, C. E. Butler, I. Sethi, Y. Zheng, J. S. King, P.

Zhou, S. Suo, Y. Guo, Q. Ma, G. C. Yuan, W. T. Pu, Massively parallel in vivo CRISPR screening identifies RNF20/40 as epigenetic regulators of cardiomyocyte maturation. *Nat Commun* **12**, 4442 (2021).

66. R. Gilsbach, S. Preissl, B. A. Gruning, T. Schnick, L. Burger, V. Benes, A. Wurch, U. Bonisch, S. Gunther, R. Backofen, B. K. Fleischmann, D. Schubeler, L. Hein, Dynamic DNA methylation orchestrates cardiomyocyte development, maturation and disease. *Nat Commun* **5**, 5288 (2014).

67. Z. Li, F. Yao, P. Yu, D. Li, M. Zhang, L. Mao, X. Shen, Z. Ren, L. Wang, B. Zhou, Postnatal state transition of cardiomyocyte as a primary step in heart maturation. *Protein Cell* **13**, 842–862 (2022).

68. K. van Duijvenboden, D. E. M. de Bakker, J. C. K. Man, R. Janssen, M. Gunthel, M. C. Hill, I. B. Hooijkaas, I. van der Made, P. H. van der Kraak, A. Vink, E. E. Creemers, J. F. Martin, P. Barnett, J. Bakkers, V. M. Christoffels, Conserved NPPB+ Border Zone Switches From MEF2- to AP-1-Driven Gene Program. *Circulation* **140**, 864–879 (2019).

69. A. S. Hinrichs, D. Karolchik, R. Baertsch, G. P. Barber, G. Bejerano, H. Clawson, M. Diekhans, T. S. Furey, R. A. Harte, F. Hsu, J. Hillman-Jackson, R. M. Kuhn, J. S. Pedersen, A. Pohl, B. J. Raney, K. R. Rosenbloom, A. Siepel, K. E. Smith, C. W. Sugnet, A. Sultan-Qurraie, D. J. Thomas, H. Trumbower, R. J. Weber, M. Weirauch, A. S. Zweig, D. Haussler, W. J. Kent, The UCSC Genome Browser Database: update 2006. *Nucleic Acids Res* **34**, D590–598 (2006).

70. J. T. Robinson, H. Thorvaldsdottir, W. Winckler, M. Guttman, E. S. Lander, G. Getz, J. P. Mesirov, Integrative genomics viewer. *Nat Biotechnol* **29**, 24–26 (2011).

71. D. M. Cable, E. Murray, L. S. Zou, A. Goeva, E. Z. Macosko, F. Chen, R. A. Irizarry, Robust decomposition of cell type mixtures in spatial transcriptomics. *Nat Biotechnol* **40**, 517–526 (2022).

72. M. F. Oliveira, J. P. Romero, M. Chung, S. R. Williams, A. D. Gottscho, A. Gupta, S. E. Pilipauskas, S. Mohabbat, N. Raman, D. J. Sukovich, D. M. Patterson, H. D. D. T. Visium, S. E. B. Taylor, High-definition spatial transcriptomic profiling of immune cell populations in colorectal cancer. *Nat Genet* **57**, 1512–1523 (2025).

73. M. Patterson, L. Barske, B. Van Handel, C. D. Rau, P. Gan, A. Sharma, S. Parikh, M. Denholtz, Y. Huang, Y. Yamaguchi, H. Shen, H. Allayee, J. G. Crump, T. I. Force, C. L. Lien, T. Makita, A. J. Lusis, S. R. Kumar, H. M.

Sucov, Frequency of mononuclear diploid cardiomyocytes underlies natural variation in heart regeneration. *Nat Genet* **49**, 1346–1353 (2017).

74. K. Alkass, J. Panula, M. Westman, T. D. Wu, J. L. Guerquin-Kern, O. Bergmann, No Evidence for Cardiomyocyte Number Expansion in Preadolescent Mice. *Cell* **163**, 1026–1036 (2015).

75. J. M. Stein, U. Arslan, M. Franken, J. C. de Greef, E. H. S, N. Mohammadi, V. V. Orlova, M. Bellin, C. L. Mummery, B. J. van Meer, Software Tool for Automatic Quantification of Sarcomere Length and Organization in Fixed and Live 2D and 3D Muscle Cell Cultures In Vitro. *Curr Protoc* **2**, e462 (2022).

76. Y. Zhu, M. Ackers-Johnson, M. K. Shanmugam, L. S. Pakkiri, C. L. Drum, Y. Chen, J. Kim, W. G. Paltzer, A. I. Mahmoud, W. L. W. Tan, M. C. J. Lee, J. Jiang, D. A. T. Luu, S. L. Ng, P. Y. Q. Li, A. Wang, R. Qi, G. J. X. Ong, T. Y. Ng, J. J. Haigh, Z. Tiang, A. M. Richards, R. S. Y. Foo, Asparagine Synthetase Marks a Distinct Dependency Threshold for Cardiomyocyte Dedifferentiation. *Circulation* **149**, 1833–1851 (2024).

77. E. Himelman, M. A. Lillo, J. Nouet, J. P. Gonzalez, Q. Zhao, L. H. Xie, H. Li, T. Liu, X. H. Wehrens, P. D. Lampe, G. I. Fishman, N. Shirokova, J. E. Contreras, D. Fraidenraich, Prevention of connexin-43 remodeling protects against Duchenne muscular dystrophy cardiomyopathy. *J Clin Invest* **130**, 1713–1727 (2020).

78. W. A. Basheer, S. Xiao, I. Epifantseva, Y. Fu, A. G. Kleber, T. Hong, R. M. Shaw, GJA1-20k Arranges Actin to Guide Cx43 Delivery to Cardiac Intercalated Discs. *Circ Res* **121**, 1069–1080 (2017).

79. H. Wickham, *ggplot2: Elegant Graphics for Data Analysis* (Springer-Verlag New York, 2016).

80. D. S. Day, B. Zhang, S. M. Stevens, F. Ferrari, E. N. Larschan, P. J. Park, W. T. Pu, Comprehensive analysis of promoter-proximal RNA polymerase II pausing across mammalian cell types. *Genome Biol* **17**, 120 (2016).

81. T. J. Aballo, J. Bae, W. G. Paltzer, E. A. Chapman, A. J. Perciaccante, M. R. Pergande, R. J. Salamon, D. J. Nuttall, M. W. Mann, Y. Ge, A. I. Mahmoud, Integrated proteomics identifies troponin I isoform switch as a regulator of a sarcomere-metabolism axis during cardiac regeneration. *Cardiovasc Res* **121**, 1240–1253 (2025).

82. J. C. Oliveros.

83. R. Patrick, V. Janbandhu, V. Tallapragada, S. S. M. Tan, E. E. McKinna, O. Contreras, S. Ghazanfar, D. T. Humphreys, N. J. Murray, Y. T. H. Tran, R. D.

Hume, J. J. H. Chong, R. P. Harvey, Integration mapping of cardiac fibroblast single-cell transcriptomes elucidates cellular principles of fibrosis in diverse pathologies. *Sci Adv* **10**, eadk8501 (2024).

84. P. Mittal, C. W. M. Roberts, The SWI/SNF complex in cancer - biology, biomarkers and therapy. *Nat Rev Clin Oncol* **17**, 435–448 (2020).
